# Lipid-based nanoparticles deliver mRNA to reverse the pathogenesis of lysosomal acid lipase deficiency in a preclinical model

**DOI:** 10.1101/2024.12.07.627318

**Authors:** Matthias Zadory, Elliot Lopez-Vince, Hamza Haddouch, Samuel Babity, Nastaran Rezaei, Fatma Moawad, Rita Maria Kenaan El Rahbani, Nemanja Vujic, Vincent Q. Trinh, Dagmar Kratky, Simon-Pierre Gravel, Davide Brambilla

## Abstract

Lysosomal acid lipase (LAL) is the only known enzyme that degrades cholesteryl esters (CEs) and triglycerides (TGs) in the lysosomes. LAL deficiency (LAL-D) results in hepatosplenomegaly with extensive accumulation of CEs and TGs, and can in the most severe cases be a life-threatening condition in early infancy. Using messenger ribonucleic acid (mRNA) for protein replacement is an innovative approach for the treatment of genetic disorders, but is challenged by a safe and efficient mRNA delivery. Here, we generated a combinatorial library of lipid-based nanoparticles (LNPs) for mRNA delivery and screened it *in vitro* and *in vivo,* which yielded a new formulation with a superior potency than an FDA-approved nanoformulation. This formulation efficiently delivered LAL mRNA and restored LAL activity in liver and spleen, mediating significant reversal of the pathological progression in an aggressive preclinical model of LAL-D. *In vivo,* the new formulation also promoted a more sustained and quantitatively higher LAL expression. In addition, repeated administration regimen mitigated hepatosplenomegaly, and targeted lipidomic analysis revealed strong diminution of CEs and TGs and of toxic lipid species in the liver and spleen. Transcriptomic analysis showed significant attenuation of inflammatory processes, fibrosis and several pathological pathways associated to LAL-D. These findings provide strong evidence that the intracellular production of LAL via mRNA-LNP is a very promising approach for the chronic treatment of LAL-D and support the clinical translation of mRNA therapy to overcome the challenges associated with traditional enzyme replacement therapies.

**One Sentence Summary:** Screening of a mRNA-LNPs library yielded a formulation with outmost potency and mitigated the progression of LAL deficiency in a preclinical model.

## INTRODUCTION

Lysosomal acid lipase (LAL) deficiency (LAL-D) combines rare autosomal recessive genetic disorders caused by the complete absence or insufficient LAL activity in patients (1). Mutations in the lipase A (*LIPA*) gene lead to impaired hydrolysis of cholesteryl esters (CEs) and triglycerides (TGs) in the lysosomes, provoking lipid accumulation in affected organs such as the liver and spleen, which in turn result in the development of severe pathological manifestations known as Wolman disease (WD) and CE storage disease (CESD) (2-4). WD is a devastating condition that manifests already in infants and is characterized by hepatosplenomegaly, progressive liver failure, cachexia, and steatorrhea. These symptoms collectively impair the patients’ ability to thrive and result in mortality within the first 6 months of life (4-6). In contrast, residual LAL activity of 1-12% in CESD patients results in a less severe form and a later onset, with patients reaching adulthood but exhibiting accelerated atherosclerosis, high risk of cardiac diseases, liver fibrosis, and cirrhosis (7, 8). The liver of LAL-D patients is characterized by marked microvesicular steatosis in hepatocytes and Kupffer cells, elevated fibrosis deposits and cirrhosis as the disease progresses, making the liver the primary target organ for the treatment strategy (9).

Multiple therapeutic strategies for the treatment of LAL-D have been investigated, with approaches varying depending on the severity of the disease. However, the majority of the clinical outcomes reported poorly favorable outcomes. Statins have been tested for late-onset LAL-D to reduce endogenous cholesterol production and cumulative cholesterol burden, though their efficacy and tolerability in clinical trials have been limited (10, 11). In 2015, the Food and Drug Administration (FDA) approved a human LAL enzyme replacement therapy (ERT) (Sebelipase alfa, Kanuma®). However, certain cases reported the development of anti-drug antibodies, particularly in patients with early-onset disease, requiring the adjustment of the dosage regimen or dose escalation (12, 13). In addition, liver transplantation and hematopoietic stem cell (HSC) transplantation have been used as a treatment option, mostly with poor outcomes (14). Therefore, the pursuit of more efficient and better-tolerated therapies represents a crucial avenue to achieve optimal treatment of LAL-D patients and innovative gene- and cell-based therapies may be a viable approach.

Messenger ribonucleic acid (mRNA) therapies have emerged as a promising approach for treating protein deficiencies. However, their development has been impeded by the challenge of achieving effective and safe *in vivo* delivery systems, for which lipid-based nanoparticles (LNPs) offer the ideal solution (15, 16). LNPs are generally generated with a four-component lipid system building a vast chemical space to optimize the formulation and promote most favorable biodistribution and mRNA-induced protein expression (17, 18). Here, we assessed the potential of mRNA-LNP therapy to reverse the pathological process of LAL-D in a highly relevant preclinical model. We first generated an *in vitro* LAL-deficient model to screen a combinatorial library of mRNA-LNP formulations and select formulations with the highest ability to promote mRNA-induced LAL restoration. *In vivo* evaluation yielded a formulation that demonstrated effective mRNA delivery and protein expression in a LAL-deficient (LAL^−/−^) mouse model. Importantly, given the chronic nature of the condition, lipidomic analysis and RNA sequencing showed that long-term administration of the formulation could reduce the accumulation of lipids in the liver and the spleen, and mitigates the inflammatory processes and fibrosis in the liver.

## RESULTS

### Development of siRNA-mediated LAL knockdown model for *in vitro* screening of mRNA-LNPs library

Short interfering RNA (siRNA) technology was utilized to mediate a knockdown (KD) of LAL at the mRNA and protein level in an immortalized hepatocyte cell line (HepG2) (**Figure 1A**). As the 3’UTR is specific to the endogenous *LIPA* mRNA and not to the exogenous mRNA sequence encapsulated in the LNPs, two siRNA sequences targeting this region were designed and tested. To thoroughly characterize KD efficiency, *LIPA* mRNA levels were assessed by reverse transcription quantitative real-time polymerase chain reaction (RT-qPCR), LAL enzymatic activity was measured with a modified fluorescence assay using the fluorogenic substrate 4-methylumbelliferyl oleate (4-MUO), and protein levels were visualized by immunoblotting. Given the transient nature of the siRNA-mediated KD, a kinetic evaluation was performed to evaluate its persistence over time. KD of 92% and 93% at the mRNA levels was achieved three days post-administration with siLIPA1 and siLIPA2, respectively, compared to the control non-targeting siRNA (siCtl), and remained between 75% and 90% during the assessed period of fifteen days (**Figure 1B**). Similarly, a drastic decrease in LAL activity between 81% and 85% persisted with no difference between both siRNA sequences. Additionally, the absence of LAL protein expression by immunoblotting confirmed the robust and stable protein KD induced by siLIPA1 and siLIPA2 (**Figure 1C**). Although cell counts of HepG2 cells treated with siLIPA1 and siLIPA2 showed reduced proliferation three days post-transfection, no differences between the three conditions were observed at subsequent time points, demonstrating that cell viability was maintained despite potent *LIPA* KD (**Figure 1D**).

**Figure 1.**
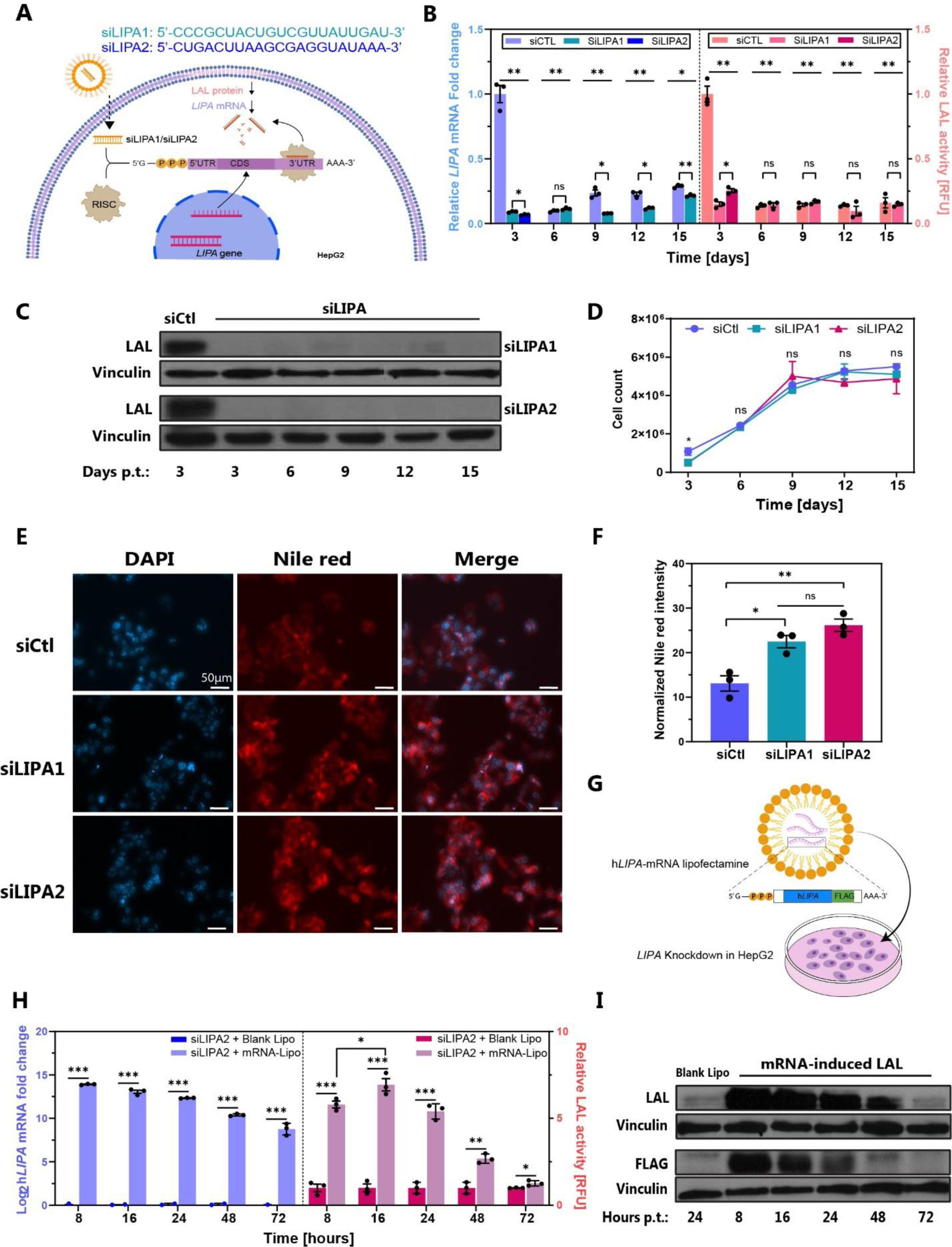
siRNA-mediated LAL knockdown *in vitro* model. **(A)** Scheme of siRNA-induced LAL knockdown (KD) in HepG2 cells. siLIPA1 and siLIPA2 bind specifically to the 3’UTR region of endogenous *LIPA* mRNA. **(B)** Kinetic evaluation of LAL KD over 15 days, with endogenous *LIPA* mRNA levels on the left y-axis in fold change and relative LAL activity on the right y-axis. Relative values are normalized on cells treated with siCtl. **(C)** Western blot of LAL in HepG2 cells treated with siCtl, siLIPA1, and siLIPA2. Vinculin served as a loading control. **(D)** Proliferation curve of siRNA-treated HepG2 cells. **(E)** Micrograph of lipid accumulation in cells treated with siCtl, siLIPA1, and siLIPA2 three days post-transfection. Cells were stained with Nile red and 4′,6-diamidino-2-phenylindole (DAPI) to visualize neutral lipids and nuclei, respectively. Magnification, 20x. Scale bars, 50 μm. **(F)** Quantification of the Nile red intensities. **(G)** Scheme of the rescue experiment with the designed exogenous human *LIPA* (h*LIPA*). h*LIPA* was transfected into HepG2 cells that had been treated with siLIPA2. As a control, cells treated with siLIPA2 were transfected with lipofectamine without mRNA (Blank Lipo). **(H)** Kinetics of the rescue experiment over three days with the exogenous mRNA levels on the left y-axis in log_2_ fold change and with relative LAL activity levels on the right y-axis. Relative values are normalized to cells treated with siLIPA2+Blank Lipo. **(I)** Western blot of LAL with vinculin as a loading control. (B, D, F) One-way ANOVA with Tukey’s post hoc test for each time points. (H) Unpaired two-tailed Student’s *t*-test for each time point. *P < 0.05, **P < 0.01, ***P < 0.001. Graphs show mean ± SEM of N=3 independent experiments. CDS, CoDing Sequence, ns, non-significant; RFU, relative fluorescence units.

Since LAL is responsible for neutral lipid degradation in the lysosome, we investigated intracellular lipid accumulation. Nile red staining revealed a more intense signal in siRNA-treated cells than in those treated with siCtl three days post-transfection, indicative of a LAL-deficient phenotype (**Figures 1E and 1F**). Although there were no statistical differences between siLIPA1 and siLIPA2 in the KD-induced lipid accumulation, siLIPA2 exhibited a stronger KD at the mRNA level, and thus was selected for the following experiments. Overall, the generated siRNA-mediated *LIPA* KD led to a marked reduction in *LIPA* mRNA expression and LAL protein expression and activity, with viable HepG2 cells suitable for *in vitro* screening of mRNA-LNPs.

Next, we designed the *LIPA* mRNA sequence to be encapsulated into LNPs based on the human *LIPA (*h*LIPA)* sequence (NCBI, NM001127605.3) and added a C-terminal FLAG-tag to distinguish exogenous from endogenous *LIPA* in subsequent experiments (**Figure 1G, Figure S1**). A rescue experiment was conducted with lipofectamine-based h*LIPA* mRNA-lipoplexes and the restoration of h*LIPA* mRNA and LAL activity was assessed over time. A primer pair specific to the FLAG-tag sequence allowed the evaluation of the intracellular delivery of exogenous h*LIPA* mRNA in HepG2, which peaked at 8 hours post-transfection, followed by a pronounced decline after 24 hours with a substantial amount of h*LIPA* mRNA remaining after 72 hours (**Figure 1H**). Interestingly, recovery of LAL activity showed a different profile, with a peak at 16 hours post-transfection, followed by a decrease with no activity measured after 72 hours (**Figure 1H**), which was corroborated by immunoblotting (**Figure 1I**) and confirmed the effective rescue of LAL in the KD model.

### Identification of hit formulations by *in vitro* screening of mRNA-LNPs library

LNPs are composed of four lipid components, namely the ionizable lipid, the helper lipid (phospholipid), cholesterol and poly(ethylene glycol) (PEG) (18, 19). To build the library, we selected distinct lipids with established mRNA delivery potency (**Figure 2A**). Dlin-MC3-DMA is the ionizable lipid of the siRNA-based approved drug Onpattro® and has been extensively used for siRNA, mRNA, and plasmid DNA delivery (17). The ionizable lipid 5A2-SC8 has been described for efficient delivery of siRNA and mRNA in preclinical models (20-22). While MC3 is commercially available, 5A2-SC8 was synthesized and characterized as a dendrimer-based ionizable lipid after purification (**Figure S2-S4**). Phospholipids have been selected based on their variation in the headgroup composition and degree of aliphatic chain saturation, since these features have been proven to impact the efficacy of nucleic acid delivery of the formulations (23, 24). We included 1,2-dioleoyl-sn-glycero-3-phosphoethanolamine (DOPE) and 1,2-dioleoyl-sn-glycero-3-phosphocholine (DOPC) as phospholipids with unsaturated aliphatic chain and distinct headgroups. 1,2-distearoyl-sn-glycero-3-phosphocholine (DSPC) present in Onpattro®, 1,2-dipalmitoyl-sn-glycero-3-phosphocholine (DPPC), 1,2-dimyristoyl-sn-glycero-3-phosphocholine (DMPC) and hydrogenated soy L-α-phosphatidylcholine (HSPC) are all choline-based phospholipids with various alkyl chain length. By combining different lipids and varying their molar ratio (**Figure 2B**), a library including 53 mRNA-LNPs was constructed (**Table S1**).

**Figure 2:**
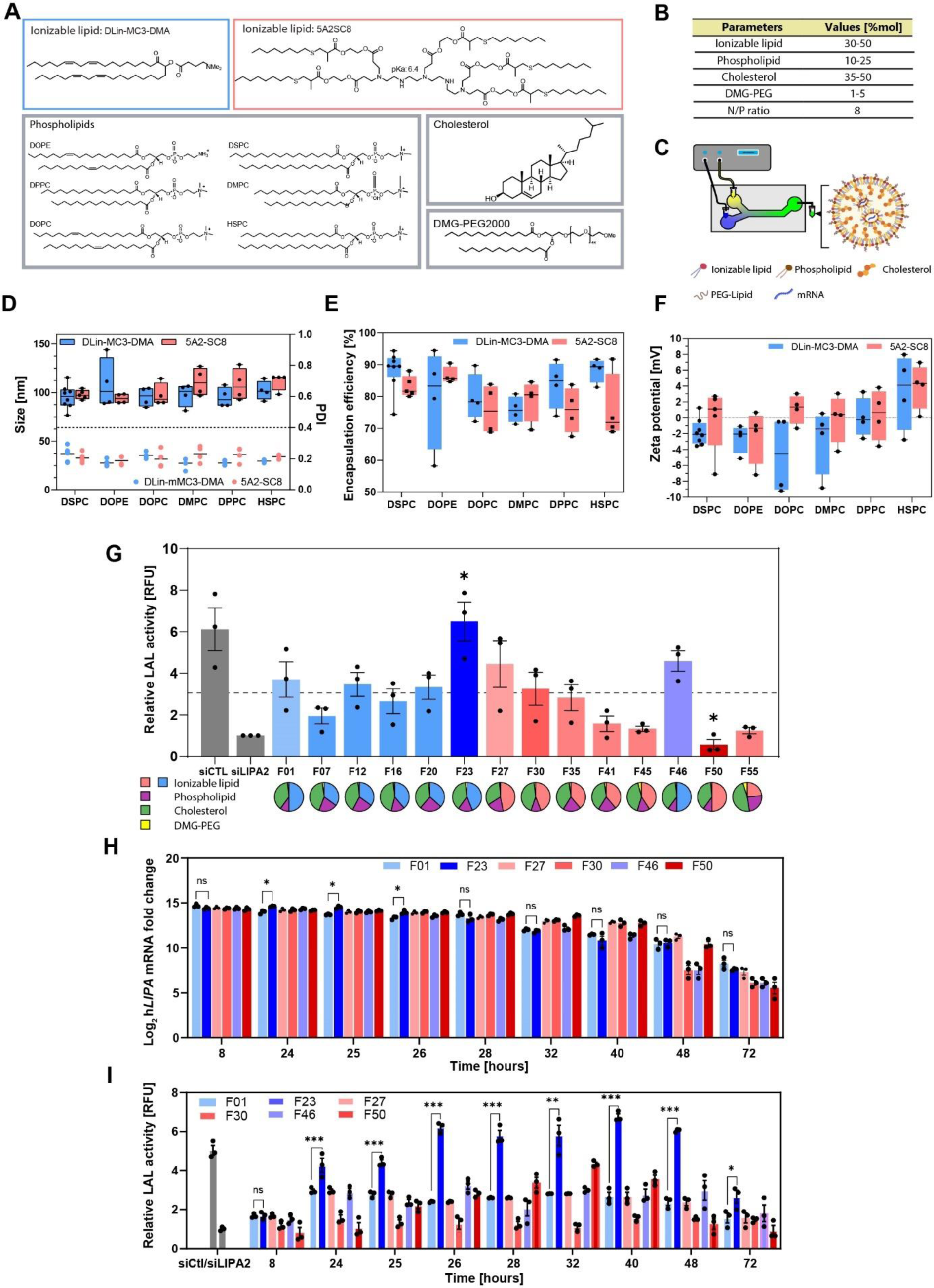
*In vitro* screening of a library of mRNA-LNP formulations. **(A)** Chemical structures and **(B)** molar percentages of lipids used in the mRNA-LNPs library. **(C)** Representation of the microfluidic mixing device setup for the standardized generation of mRNA-LNP formulations. **(D)** Physicochemical characterization of the selected formulations based on the hydrodynamic size (box plots; left y-axis) and polydispersity index (dots; right y-axis), **(E)** encapsulation efficiency, and **(F)** surface charge. Each point represents the mean of 3 independent measurements for one formulation. **(G)** Screening of selected mRNA-LNP formulations in siLIPA2 KD cells with siCtl representing the basal activity and the dotted line 50% of the basal activity. For each formulation, the mean ± SEM of 3 independent experiments is shown. **(H)** Kinetic evaluation of h*LIPA* mRNA delivery to HepG2 cells and **(I)** mRNA-induced restoration of LAL activity. Graphs show the mean ± SEM of 3 independent experiments. (D - F) Boxplots representing the median, the upper and lower quartile, and minimum and maximum whiskers in each group. (G) One-way ANOVA with Tukey’s post hoc test analysis on all formulations. (H) and (I) Unpaired two-tailed Student’s *t*-test between F01 and F23. *P < 0.05, **P < 0.01, ***P < 0.001.

Reproducible and standardized production of mRNA-LNPs was achieved by a customized microfluidic system (**Figure 2C**), enabling precise monitoring of the pressure and flow rate in a linear correlation (**Table S2, Figure S5**). To optimize the mixing output, the flow rate was gradually increased to produce particles with a hydrodynamic diameter of 100 ± 10 nm and a polydispersity index (PDI) of ≤ 0.2, demonstrating the homogeneity of the particle population and high encapsulation efficiency (**Figure S6**). Previous middle and high throughput screenings of LNP libraries typically did not include physicochemical properties of LNPs, but rather conducted directly the screening process with little knowledge of the individual size and encapsulation levels of the LNPs. Therefore, we sought to characterize the physicochemical properties of the LNP library by size, encapsulation efficiency, and charge to identify potential differences associated with the choice of lipids (**Figures 2D, 2E, 2F**). No significant trend in the physicochemical properties was observed, which may be attributed to the highly standardized mixing procedure. Based on the appropriate size (≤ 100 nm) and optimal encapsulation efficiency (≥85%), we selected a set of 14 formulations for *in vitro* screening. Prior to this, a dose-response study performed with the standard formulation F01 (based on Onpattro^®^) defined the optimal dosage for the screening process (**Figure S7)**. At a dose of ≥1.0 µg of h*LIPA* mRNA, mRNA-induced LAL expression and activity was decreased, suggesting that maximal delivery was attained, as evidenced by immunoblots. Since a strong restoration of LAL activity was already achieved at a dosage of 0.5 μg h*LIPA* mRNA, this dose was chosen for subsequent *in vitro* screening experiments. Kinetic evaluation of intracellular delivery of h*LIPA* mRNA showed an increase, reaching a plateau 24 hours post-transfection, while the mRNA amount in the medium inversely decreased (**Figure S8**). Consequently, we performed the *in vitro* screening with 0.5 µg and harvested the cells 24 hours post-transfection.

The *in vitro* screening of the selected 14 formulations revealed that the LNP composition in ionizable lipids and phospholipids considerably impacted the potency of the LNPs in restoring LAL activity (**Figure 2G**). Notably, a dramatic decrease of the enzymatic activity was observed in the 5A2-SC8-based formulations comprising DMPC (F45) and HSPC (F50, F53), compared to DSPC (F27). Similarly, DLin-MC3-DMA-based LNPs tend to profit from the phospholipid DSPC at a molar ratio of 10. From this screening, a formulation (F23) containing the helper lipid DPPC displayed significant superiority compared to the standard Onpattro® formulation (displayed as F01). In contrast, DPPC had a negative effect when associated with the ionizable lipid 5A2-SC8 (F41), demonstrating that an ideal phospholipid combination depends on the ionizable lipid and its molar ratio. In general, LNPs containing the ionizable lipid DLin-MC3-DMA demonstrated superior performance, with five formulations being able to restore 50% of the basal activity (F01, F12, F20, F23, and F46) compared to two formulations containing 5A2-SC8 (F27 and F30).

Next, we evaluated the kinetics of mRNA delivery by qPCR and the extent of mRNA-induced recovery of LAL activity after a single transfection with the control F01 and five formulations displaying high, middle, and low levels of LAL restoration. mRNA delivery was comparable between the formulations (**Figure 2H)**, but F23 demonstrated a superior and more sustained mRNA-induced LAL activity (**Figure 2I**). Formulations containing 5A2-SC8 as an ionizable lipid were comparable to MC3-based LNPs, but showed a delayed peak in protein expression. Most strikingly, F50, which failed to induce substantial protein expression 24 hours post-transfection, showed a pronounced LAL restoration with a peak at 32 hours and a more sustained expression profile over time, resulting in an increased and higher area under the curve (AUC) than F30 and F46, but not F27 (**Table S3**). Any observed differences were not biased by potential cytotoxicity of the formulations, as cell viability was unaltered by all tested LNPs at 0.5 μg mRNA (< 100 μg/mL lipids) (**Figure S9**). Based on the *in vitro* screening results, four formulations (F01, F23, F27, and F50) with the highest recovery of LAL activity and AUC in the KD model were selected for *in vivo* evaluation.

### *In vivo* evaluation in healthy mice identified a novel formulation with favorable biodistribution and high mRNA-induced LAL activity

Having selected the most promising formulations *in vitro*, we next assessed their biodistribution, pharmacokinetics, h*LIPA* mRNA delivery, and LAL protein expression in healthy mice. The biodistribution study of fluorescently labeled LNPs showed favorable accumulation in the liver and, to a lesser extent, in the spleen for all formulations, since these organs are the primary filtration sites of LNPs (**Figure 3A, 3B**) (25). Although previous studies have shown the distribution of 5A2-SC8 in splenic tissues (21), we observed that LNPs with the ionizable lipid 5A2-SC8 exhibited a more pronounced accumulation in the spleen than MC3-based LNPs. RT-qPCR analysis of FLAG-tagged h*LIPA* mRNA revealed a pronounced accumulation in the liver and spleen, with higher levels being delivered by F27 and F50 in the latter and almost no exogenous h*LIPA* mRNA delivered to the lungs, corroborating the biodistribution profile obtained with fluorescently labeled LNPs (**Figure S10**). The pharmacokinetic analysis of the formulations showed comparable profiles and pharmacokinetic parameters, with F23 having the largest AUC compared to all other formulations **(Figure S11)**.

**Figure 3.**
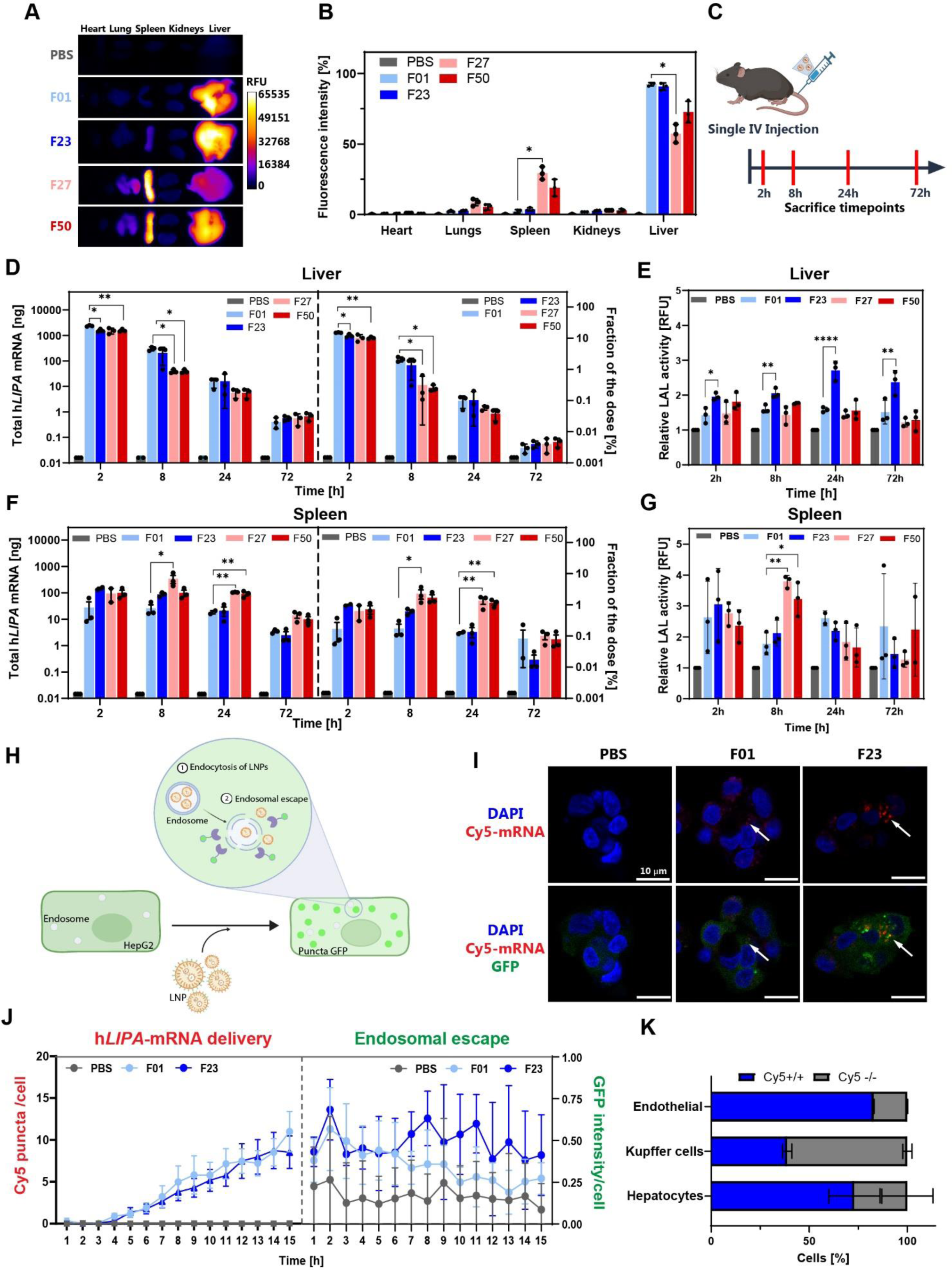
*In vivo* evaluation of selected mRNA-LNPs. **(A)** Biodistribution analysis of selected DiIC18(5); 1,1′-dioctadecyl-3,3,3′,3′-tetramethylindodicarbocyanine, 4-chlorobenzenesulfonate (DiD)-labeled LNPs in CD-1 mice following intravenous administration (0.5 μg h*LIPA* mRNA /g body weight). **(B)** Fluorescence values relative to total fluorescence. **(C)** Schematic representation of the kinetic study. **(D)** Exogenous h*LIPA* mRNA levels delivered and **(E)** mRNA-induced LAL activity in the liver and **(F, G)** spleen. Relative fluorescence values are normalized to levels in PBS-treated mice. **(H)** Schematic representation of the Gal8-GFP *in vitro* model to monitor endosomal escape in HepG2 cells. LNPs enter the cell via endocytosis and traffic through the cytoplasm within the endosome. Endosomal escape describes the fusion of the LNP and disruption of the endosomal membrane to release its cargo, triggering recruitment of galectin-8 fused with GFP (Gal8-GFP), thereby producing clear green punctae in the cytoplasm. **(I)** Confocal images of the Gal8-GFP reporter cells 8 hours post-F01 and -F23 transfection in maximum intensity projections with nuclei stained with Hoechst (blue) and mRNA labeled with Cy5 (red). Scale bars, 10 μm. **(J)** Live confocal image acquisition over 16 hours with Cy5, Hoechst, and GFP signals being recorded every 60 minutes at 5 different fields. **(K)** Flow cytometry analysis of Cy5-mRNA positive hepatic cell populations transfected with F23. Data are expressed as mean ± SD (n = 3). (B) One-way ANOVA with Tukey’s post hoc test analysis for each organ. (D), (E), (F), and (G) One-way ANOVA with Tukey’s post hoc test analysis for each time points separately. *P < 0.05, **P < 0.01, and ****P < 0.0001. (D), (E), (F), and (G) Graphs show mean ± SEM of n=3 mice. ns, non-significant; RFU, Relative Fluorescence Units.

We then examined the ability of the formulations to deliver h*LIPA* mRNA to the liver and the spleen and to promote LAL activity following a single-dose administration (**Figure 3C**). Because commonly used fluorescence or radiolabeling methods do not allow quantitative assessment of mRNA in an organ, we quantified the amount of encapsulated h*LIPA* mRNA delivered to the tissues at the different time points by RT-qPCR (**Figure S12**), as recently described (26-28). Two hours post-administration, 14 ± 2.3% of the total mRNA administered in F01 was found in the liver, with a gradual decrease to < 0.003 ± 0.0001% at 72 hours (**Figure 3D**). Similar to the *in vitro* experiments, all formulations had comparable capacity to deliver h*LIPA* mRNA to the liver. In contrast, the h*LIPA* mRNA-induced LAL activity by F23, demonstrated again superior activity in liver tissue compared to all other formulations at all time points (**Figures 3E**). In the spleen, the remaining fraction of the dose at 2-hours post-administration was ∼ 10-to 20-fold lower, indicating that the majority of the administered mRNA is filtered in the hepatic tissues (**Figure 3F**). h*LIPA* mRNA-induced LAL activity in the spleen was higher for the formulations F27 and F50 compared to F01 at 8- and 24-hours post-administration, however, the AUC was not significantly altered (**Figure 3G, Table S4**). Overall, F23 yielded the highest LAL activity in the liver with a 2.4-fold increase in the hepatic AUC compared to all formulations, along with comparable AUC in the spleen, which established F23 as the best lead in the library.

Next, we pursued to understand the underlying mechanism of F23 superiority, focusing on the endosomal escape, which is thought to drive the efficacy of LNPs to promote mRNA-induced protein expression (29). To this end, we established a cell-based fluorescent model previously described, expressing galectin-8 fused with a green fluorescent protein (Gal8-GFP) to monitor the endosomal escape of LNPs (**Figure 3H**) (30). Live confocal imaging enabled the visualization of the Gal8-GFP signal, indicative of successful endosomal escape, Cy5-labeled mRNA to track the intracellular delivery, and Hoechst to visualize cell nuclei (**Figure 3I**). By tracking only the Cy5 channel, we validated similar internalization profiles between F23 and F01, and any differences in the Gal8-GFP signal can be attributed solely to endosomal escape of the LNPs and not to the variation in delivery (**Figure 3J**). Up to 8 hours post-transfection, endosomal escape was comparable between both formulations. After that, the Gal8-GFP profiles diverged, with a stronger signal present in the cells treated with F23 compared to F01, suggesting a higher capability of F23 to escape the endosome and more efficiently deliver mRNA into the cytoplasm.

Hepatocytes, resident liver macrophages (Kupffer cells), and other liver cell subtypes are largely involved in the development of hepatic steatosis, lipid metabolism, and inflammation and thus play a pivotal role in the pathogenesis of LAL-D (3, 31). Using Cy5-labeled h*LIPA* mRNA and flow cytometry, we determined the percentage of transfection of F23 in various liver cells (**Figure 3K and S13**). Endothelial cells exhibited the highest transfection efficiency (85%), followed by hepatocytes (76%) and Kupffer cells (44%), indicating that F23 was able to transfect the hepatic cell types crucial for a successful treatment of LAL-D.

### F23 formulation promotes superior mRNA-induced LAL restoration in LAL^−/−^ mice and mitigates hepatosplenomegaly

Having selected F23 as the best-performing formulation, we next evaluated its capacity to deliver h*LIPA* mRNA and to restore LAL activity in in LAL knockout (LAL^−/−^) mice, a preclinical model of LAL-D. In general, LAL^−/−^ mice display biochemical and histopathological features of WD but resemble the CESD phenotype with a median life span of one year (32). LAL-D manifests already during fetal development with massive neutral lipid accumulation in the liver and spleen. These animals exhibit reduced body weight and yellowish liver 2 days after birth, demonstrating the early and rapid progression of the disease (33). We generated LAL^−/−^ mice on a C57BL/6J background and used them at early age, ranging from 5 to 9 weeks, in which safe and repeated injections of the LNPs were feasible. We first performed a kinetic evaluation after a single LNP injection to assess h*LIPA*-mRNA delivery and restoration of mRNA-induced LAL activity using two dosages (0.5 and 1.0 µg/g BW) (**Figure 4A**). As observed in CD-1 mice, comparable levels of delivered h*LIPA* mRNA to the liver were obtained with F01 and F23 (**Figure 4B**). A comparison of mRNA levels at the same time points revealed no significant differences in wild-type (WT) and LAL^−/−^ mice, indicating that the pathological status did not affect the delivery efficacy of the formulations (**Figure S14**). Enzymatic activity in LAL^−/−^ mice treated with LNP blank was close to 0, reflecting the severe condition of WD patients (**Figure 4C**). At the lower dosage of 0.5 µg/g BW, both formulations showed a slight recovery of < 10% of WT activity in the liver at 48 hours. Using 1.0 µg/g BW, both formulations achieved a more potent recovery, with F23 showing higher activity and AUC than F01 at 24- and 48-hours (**Table S5**). While no significant difference in the mRNA delivery to the spleen was observed (**Figure 4D**), the recovery of the LAL activity in the spleen was surpassing that in the liver, reaching almost 20% of WT levels 48 hours post-administration for the F23 formulation (**Figure 4E**). Overall, F23 exhibited a 1.6-fold higher total AUC (**Table S5**) compared to F01, and thus was selected as the formulation for the efficacy study.

**Figure 4.**
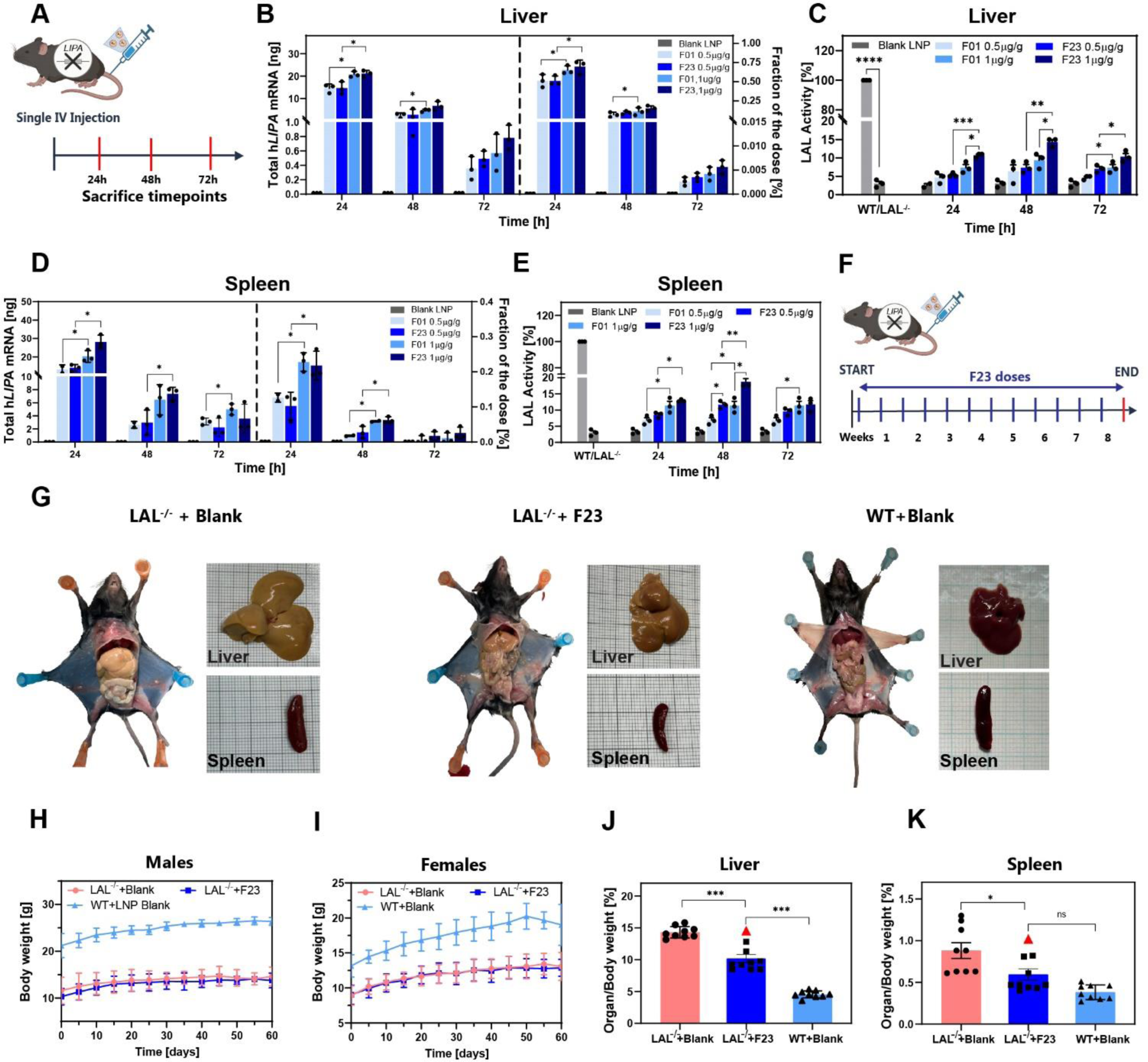
F23 formulation restores mRNA-induced LAL activity and mitigates hepatosplenomegaly in LAL^−/−^ mice. **(A)** Schematic illustration of the kinetic study in LAL^−/−^ mice. **(B)** Exogenous h*LIPA* mRNA levels delivered by F01 and F23 at 0.5 μg/g and 1.0 μg/g. **(C)** mRNA-induced LAL recovery in the liver and **(D, E)** spleen of LAL^−/−^ mice, respectively. **(F)** Schematic illustration of the efficacy study in LAL^−/−^ mice injected with F23 at 1.0 μg/g at a 5-day interval for 8 weeks. **(G)** Representative images of mice, liver (upper panel), and spleen (lower panel) from LAL^−/−^+Blank, LAL^−/−^+F23, and WT+Blank at the end of the study. **(H)** Body weight of males and **(I)** females (n=5 per group). **(J)** Liver and **(K)** spleen weights normalized to total body weight. Red triangles depict the male mouse (0629) with a later initiation of treatment. **(B-E)** One-way ANOVA with Tukey’s post hoc test analysis for each time points separately. (J) and (K) One-way ANOVA with Tukey’s post hoc test analysis *P < 0.05, **P < 0.01, ***P < 0.005. Graphs show mean ± SEM.

For the long-term efficacy study, mice were intravenously injected at 5-day intervals for 8 weeks with either empty F23 LNPs (LAL^−/−^+Blank and WT+Blank) or with F23 containing h*LIPA* mRNA (LAL^−/−^+F23) (**Figure 4F**). Body weights were monitored before each injection. One LAL^−/−^+Blank female mouse died during the study. At sacrifice, a dramatic increase in organ size and discoloration due to large fat deposition was visible in the LAL^−/−^ group treated with blank LNPs compared to the WT group, which is in line with a LAL-D phenotype (**Figure 4G**). In contrast, treatment with F23 resulted in a liver size reduction and pronounced recoloration towards WT mice, demonstrating an amelioration of the phenotype. Body weights in LNP-blank-injected LAL^−/−^ mice was significantly lower than age- and sex-matched WT controls, which was not corrected by F23 treatment. (**Figure 4H and 4I**), a trend also observed with long-term ERT (34). LAL^−/−^+Blank showed marked hepatosplenomegaly with an average of 2.8-fold increase in liver and 2.3-fold in spleen weight compared to the WT+Blank-treated mice. F23 treatment markedly reduced these increases by 25-35% and 30-47% for the liver and spleen, respectively (**Figure 4J and 4K**). It is worth mentioning that fortuitous initiation of the F23-treatment of a single male mouse (ID 0629) at 9 weeks of age failed to show improvement, indicating that early initiation of mRNA-based treatment is of paramount importance for positive outcome. Interestingly, a sex-based analysis between the groups revealed that F23 treatment of female LAL^−/−^ mice led to an almost complete normalization of the spleen and a more pronounced decrease in liver weights than in males (**Figure S15**).

### F23 formulation reduces accumulation of neutral lipids in LAL^−/−^ mice

Tissue morphology and neutral lipid accumulation in the liver and spleen were visualized by hematoxylin and eosin (H&E) and Oil Red O (ORO) staining, respectively. Numerous lipid-laden cells characterized by a pale color and macro- and microvesicular steatosis were visible in untreated LAL^−/−^ mice, which were strongly reduced in F23-treated mice (**Figure 5A and 5B**). Similarly, the intense ORO staining in untreated LAL^−/−^ mice illustrated the massive lipid accumulation, which was markedly reduced in F23-treated mice. Some cell clusters remained positive for neutral lipids despite treatment, indicating that F23 was unable to transfect all cells throughout the liver parenchyma and splenic tissue at the tested dosage. Quantification of the ORO signal intensity revealed a more prominent reduction of lipids in the spleen compared to the liver following F23 administration (**Figure 5C and 5D**).

**Figure 5.**
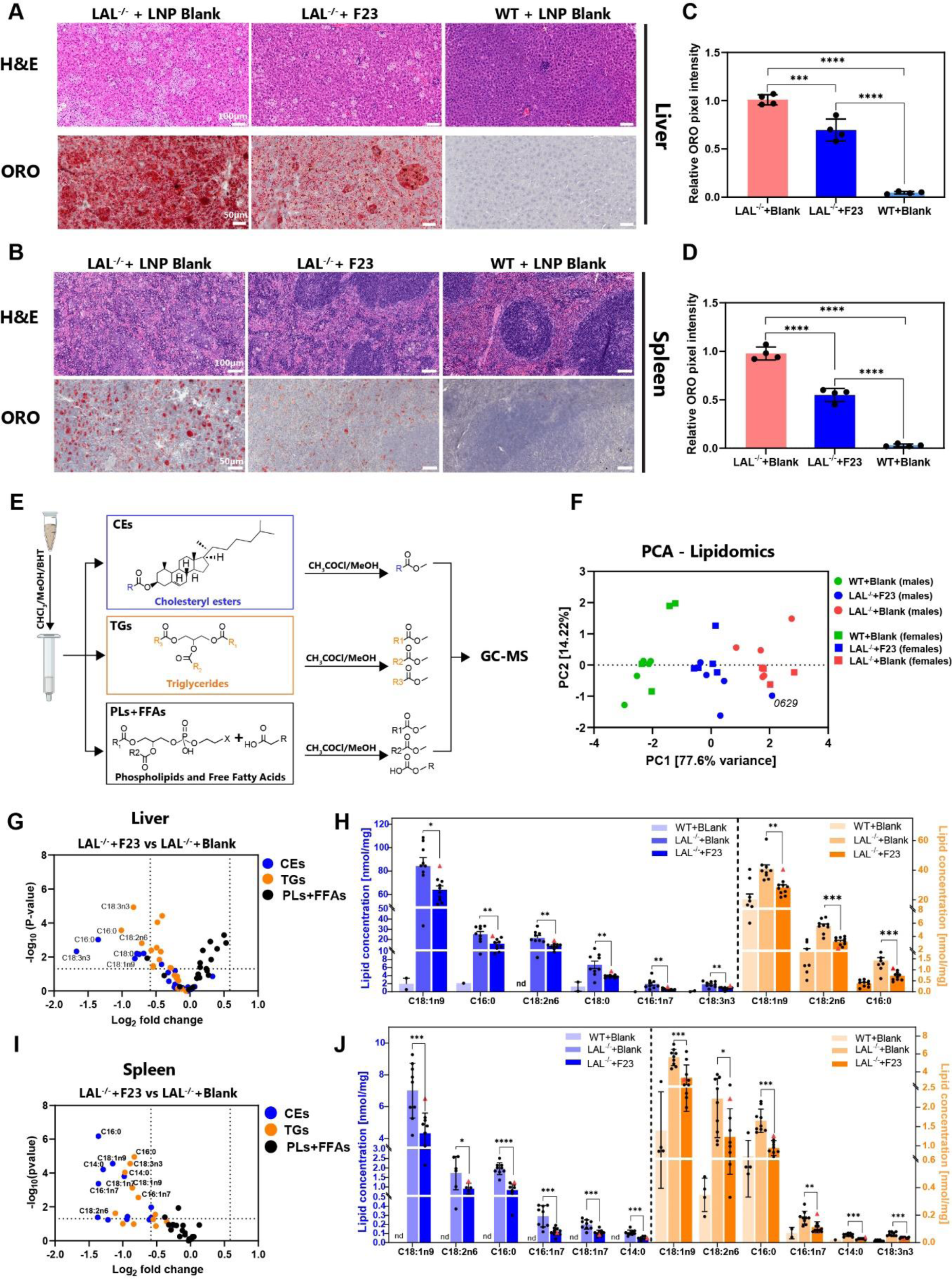
F23 formulation reduces neutral lipid accumulation in LAL^−/−^ mice. **(A)** Hematoxylin/eosin (H&E) and Oil Red O (ORO) staining of histological cross sections of liver and **(B)** spleen. Scale bars: 100 μm (H&E staining); 50 μm (ORO staining). Quantification of ORO pixel intensities in cross sections of **(C)** liver and **(D)** spleen. **(E)** Schematic overview of the lipidomic analysis in the cholesteryl ester (CE), triglyceride (TG), and phospholipid (PL)/free fatty acid (FFA) fractions. **(F)** Principal component analysis (PCA) of lipidomic data of WT+Blank, LAL^−/−^+F23, and LAL^−/−^ +Blank mice separated by sex (males in circle, females in square) with the percentage of variance for PC1 and PC2. **(G)** Volcano plots showing differential lipid levels of LAL^−/−^+F23 versus LAL^−/−^ +Blank in the liver and **(H)** Average concentrations of significantly altered CEs and TGs species, and **(I, J)** in the spleen. Plots show the fold change in log_2_ (*x*-axis) and the associated -log_10_ of the *P* values (y-axis) with dotted lines defining the threshold as ≥1.5 or ≤ -1.5 with a cutoff ≤0.05. Colored dots correspond to the isolated CE (blue), TG (orange), and PL+FFA (black) fractions. Red triangles depict the male mouse (0629) with a later initiation of treatment. (C, D) One-way ANOVA with Tukey’s post hoc test analysis. (H) and (J) unpaired two-tailed Student’s *t*-test between LAL^−/−^+blank and LAL^−/−^+F23 groups. *P < 0.05, **P < 0.01, ***P < 0.001, and ****P < 0.0001. Graphs show mean ± SEM (n≥9).

To determine which lipid species accumulated in LAL^−/−^ mice and were modulated by F23 treatment, we performed a targeted lipidomic analysis on liver and spleen tissues. After lipid extraction from the tissues, three fractions were isolated: one containing CEs, one containing TGs, and the last fraction containing phospholipids (PLs) and free fatty acids (FFAs) (**Figure 5E**). In total, 29 fatty acid methyl esters (FAMEs) in each fraction were quantified (**Table S6 and S7**). Principal component analysis (PCA) revealed distinct clusters in liver and spleen according to the condition, with a pronounced shift of the F23-treated LAL^−/−^ mice toward the WT+Blank group, indicative of a significant improvement (**Figure 5F**). In line, F23 treatment mitigated liver CEs and TGs by 30% and 25%, respectively (**Figure S16A**). Similar to ORO staining, we detected less lipid accumulation in the spleen compared to the liver, and treatment with F23 resulted in an even greater decrease of 55% and 43% of CEs and TGs, respectively (**Figure S16B**).

A Volcano plot analysis showed FAMEs that were modulated by F23 treatment with fold changes of ≥ 1.5 or ≤ -1.5 and p-values cutoff of ≤ 0.05 identifying six CE species and three TG species with the highest accumulation in the liver and strongest reduction by F23 treatment (**Figure 5G**). Interestingly, PLs and FFAs did not exhibit a noticeable increase despite the long-term injections of LNPs, indicating that the PLs contained in the LNP formulations did not affect the overall lipid load and were safely mobilized and metabolized by the cells. The most enriched CE species in the liver included oleic acid (C18:1n9), linoleic acid (C18:2n6), palmitic acid (C16:0), stearic acid (C18:0), palmitoleic acid (C16:1n7), and α-linolenic acid (C18:3n3), which were all significantly diminished by F23 treatment (**Figure 5H**). The most abundant TG species C18:1n9, C18:2n6, and C16:0 were also significantly reduced by F23. Comparison of the spleen and liver volcano plot analyses revealed that the average of log_2_ fold change value in the spleen was higher for both CEs and TGs, showing again the stronger effect of F23 in the splenic tissue with seven CE species and six TG species which were strongly reduced in the spleen following F23 treatment (**Figure 5I**). Akin to the liver, C18:1n9, C16:0, and C18:2n6 were the lipid species with the highest accumulation in the spleen and were all significantly reduced by F23 administration (**Figure 5J**). Levels of unsaturated n-6 FAs and n-3 FAs that are involved in the development of steatosis and hepatotoxicity (35-37) were significantly increased in untreated LAL^−/−^ compared to WT mice, and reduced after treatment (**Figure S17**). Furthermore, we found a 37% reduction in the proinflammatory FA arachidonic acid (AA) **(Figure S18)**. In summary, treatment of LAL^−/−^ mice with F23 decreased the accumulation of unsaturated and saturated lipotoxic lipid species, which might positively affect the pathophysiology of LAL-D.

### F23 formulation corrects liver functions and mitigates critical pathological processes in the liver of LAL^−/−^ mice

To further investigate the extent of the correction of liver metabolic functions and pathological processes, we performed RNA-sequencing (RNA-seq) on hepatic tissues. PCA analysis of the RNA-seq data showed a clear distinction in the transcriptomic profiles between the groups, with a significant shift of F23-treated LAL^−/−^ mice toward the WT group (**Figure 6A**). Of note, the score plot segregated mouse (0629) with later-onset treatment within the untreated cluster. While no distinct cluster was observed in untreated LAL^−/−^ mice according to sex, a clear separation between males and females in WT and in F23-treated LAL^−/−^ mice was noticed. The high number of up-regulated (3475) and down-regulated (1664) genes (fold change ≥2.0 or ≤-2.0, false discovery rate (FDR) ≤0.01) between the LAL^−/−^+Blank and WT+Blank groups (**Figure 6B**) indicated severe transcriptional alterations underlying the histopathological phenotype. In contrast, only 1555 up-regulated genes (55% reduction versus Blank) and 107 down-regulated genes (94% reduction versus Blank) were differentially expressed between LAL^−/−^+F23 and WT+Blank (**Figure 6C)**, demonstrating the pronounced effect of F23 on the transcriptomic profile of LAL^−/−^ mice.

**Figure 6.**
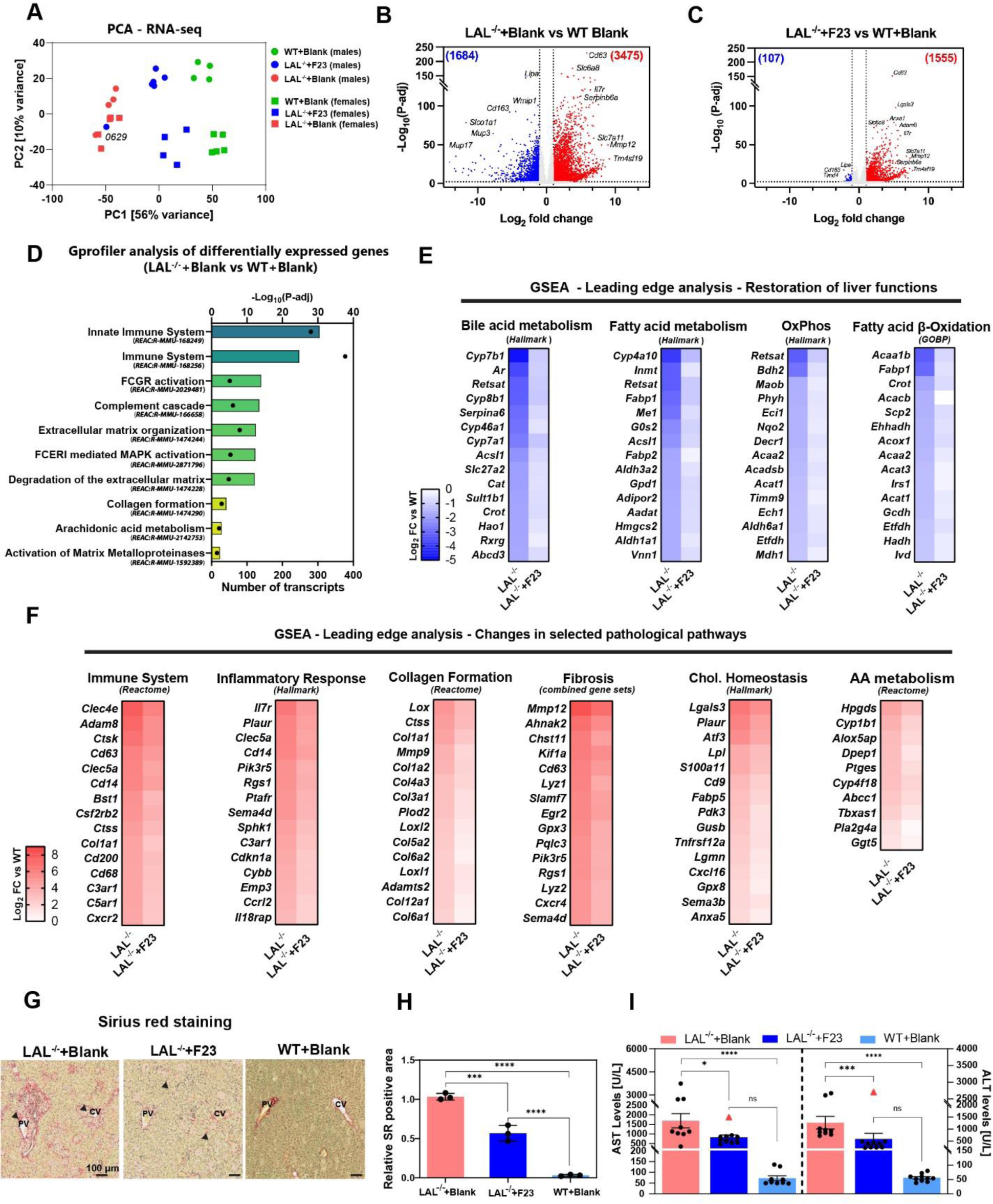
F23 formulation corrects metabolic pathways and mitigates fibrosis and inflammation-related processes in the liver of LAL^−/−^ mice. **(A)** Principal component analysis (PCA) plot of male (circles) and female (squares) WT+Blank, LAL^−/−^+F23, and LAL^−/−^+Blank, with the percentage of variance for PC1 and PC2. **(B)** Volcano plots showing up-regulated (red) and down-regulated genes (blue) hepatic gene expression of LAL^−/−^+Blank versus WT+Blank, and **(C)** LAL^−/−^+F23 versus WT+Blank mice. Dotted lines define the threshold as ≥2.0 or ≤ -2 for the average expression fold change in log_2_ (*x*-axis) and the associated -log_10_ of the adjusted *P* values (y-axis) with a cutoff ≤0.01. **(D)** Functional analysis of significantly altered genes in LAL^−/−^+Blank versus WT+Blank groups. **(E)** Heatmap representations of transcript enriched in the WT+Blank group and **(F)** enriched in the LAL^−/−^ +Blank group with fold change (Log_2_) in LAL^−/−^+Blank and LAL−/− +F23 versus WT+Blank, respectively. Transcripts are ranked according to fold change from (E) lowest (blue) to highest (white) and from (F) highest (red) to lowest (white). **(G)** Representative images of Sirius red staining of histologic sections of of LAL^−/−^ +Blank, LAL^−/−^+F23, and WT+Blank livers (scale bar, 100 μm) and **(H)** quantification. **(I)** Plasma aminotransferase concentrations. Red triangles depict the male mouse (0629) with later initiation of the treatment. Graphs show mean ± SEM (n≥9). One-way ANOVA with Tukey’s post hoc test. *P < 0.05, ***P < 0.001, and ****P < 0.0001.

Next, we performed gene functional analyses of the identified differentially expressed genes to better characterize the transcriptome signature associated with untreated LAL^−/−^ livers. Using g:Profiler, we found a strong enrichment of pathways associated with immune response, inflammation, remodeling of the extracellular matrix and fibrosis-related pathways, and sterol synthesis (**Figure 6D, Figure S19**), which are all hallmarks of LAL-D in patients and mice (38, 39). Interestingly, the main altered pathways that emerged from the analysis were not primarily related to CE or TG metabolism, but rather to the immune system and inflammation, highlighting the important role of LAL in various pathological metabolic pathways. Gene Set Enrichment Analysis (GSEA) (**Figure S20**) (40) of down-regulated gene sets (enriched in WT+Blank) identified bile and fatty acid metabolism, fatty acid ß-oxidation, and oxidative phosphorylation, which were all partially restored by F23 treatment (**Figure 6E**). Enrichment analysis of up-regulated gene sets (enriched in LAL^−/−^+Blank) included immune system, inflammatory response, collagen formation and fibrosis, cholesterol homeostasis, and AA metabolism (**Figure 6F**). F23 treatment of LAL^−/−^ mice led to a reduction in these gene sets, notably in the fibrosis pathway, for which the enrichment yielded the highest score. Sirius red staining confirmed reduced pericellular and periportal fibrosis (Metavir F2-equivalent), which was limited to the perivenular area with mild deposition surrounding the clusters (Metavir F1-equivalent) in liver sections of LAL^−/−^ mice following F23 treatment **(Figure 6G)**. Pale red staining, indicative of hepatocyte collapse/necrosis, was more prominent in the LAL^−/−^+Blank than in the F23-treated LAL^−/−^ mice. Quantification of positive SR area revealed a significant reduction of fibrosis in F23-treated mice without complete resolution to WT levels **(Figure 6H)**. H&E images revealed clusters of ballooning and necrotizing lipid-laden hepatocytes associated with a mononuclear inflammation **(Figure S21)**, which significantly decreased following F23 treatment. Despite not being restored to the WT condition, transaminase levels depicted a significant and strong reduction by F23 treatment, confirming the mitigation of both hepatocyte injury and inflammation (**Figure 6I**).

Altogether, these results demonstrate that recovery of LAL through mRNA-LNP, in addition to reducing steatosis, can restore crucial hepatic metabolic functions and attenuate many pathological processes, including liver injury, fibrosis, and immune responses triggered by the drastic accumulation of lipids.

## DISCUSSION

LAL-D exhibits a broad range of clinical manifestations that, in its severe form (WD), leads to death in early infancy (41). Over the past decades, several approaches have been clinically tested for the treatment of LAL-D, including reduction of the systemic cholesterol burden with statins and transplantation of liver or HSCs, and the most recent, ERT (5, 39). These therapeutic approaches lead in most cases to an ameliorated liver histology and decreased hepatic lipid deposition, yet display unfavorable outcomes in some cases (39, 42, 43). In a recent study, a h*LIPA-* expressing adeno-associated virus (AAV) treatment was able to restore LAL activity in LAL^−/−^ mice. However, gene expression gradually decreased over time, thus requiring re-administration that remains a challenge in AAV-based approaches (44).

Here, we described the development of an mRNA-based therapeutic approach to mediate intracellular production and restoration of LAL activity. Since LAL-D is a chronic condition with continuous and substantial accumulation of substrates within lysosomes, safe and efficient delivery of h*LIPA* mRNA to the target organ is of paramount importance to achieve sufficient enzymatic LAL activity and mitigate the pathological progression. *In vitro* and *in vivo* screening of our mRNA-LNP library identified F23 as novel formulation with superior recovery of LAL activity than the FDA-approved control formulation (F01), highlighting the importance of LNP optimization. While numerous studies focused on the modification of the ionizable lipids in LNPs, our screening data showed that largely underexplored alterations of the molar ratio and PL composition (15, 24, 45) significantly impact potency of the formulation to induce LAL activity, but not the efficacy of mRNA delivery to the cells. Furthermore, the discrepancy found between mRNA delivery and mRNA-induced LAL activity suggests a lack of direct correlation between these two processes, underlining endosomal escape as the key element that defines efficient protein expression, as indicated by the Gal8-GFP experiments. Previous studies using recombinant human LAL produced in different expression systems emphasize crucial involvement of mannose-phosphate receptors in lysosomal protein internalization (34, 46, 47). A sufficient expression of the receptors at the cell surface is thus indispensable for efficient enzyme internalization, which may limit the efficacy of ERT approaches. In contrast, F23 is able to transfect the main hepatic cell types independently of the mannose-phosphate receptor expression on the cell surface, but mainly through endocytotic internalization of entire mRNA-LNP particles (48, 49). This illustrates an additional advantage of mRNA-LNPs that may expand its therapeutic potential beyond the limitations of ERT.

Long-term administration of F23 significantly reduced hepatosplenomegaly in LAL^−/−^ mice, but similar to previous ERT-based studies (34), failed to induce improvement in body weight gain. These data suggest that the overall body weight is likely associated with extra-hepatic consequences of LAL-D pathology and a systemic dysregulation of energy homeostasis (50) that requires further studies. Comparable to ERT (34), a more pronounced weight reduction was observed for the spleen, despite a 12-fold lower mRNA delivery compared to the liver. Considering that impaired lysosomal lipid catabolism in LAL-D leads to a massive substrate deposition mostly in the liver, unfolded protein response and endoplasmic reticulum stress described in other fatty liver diseases and/or inefficient lysosomal acidification may prevent more favorable hepatic outcomes compared to the less affected spleen (51-53). Splenocytes may process mRNA more effectively to produce LAL, ultimately recovering faster from the pathological progression of LAL-D. Timely intervention is critical for therapeutic efficacy due to the early onset and rapid progression of the disease (33), as also evidenced by the lack of effect in the current study in the single LAL^−/−^ mouse with late treatment initiation.

Whereas previous studies of ERT and AAV gene-based treatments in LAL^−/−^ mice generally described the overall reduction of total cholesterol and TG burden (34, 44, 54), we presented for the first time lipidome analysis of liver and spleen from LAL^−/−^ mice and their changes induced by mRNA-LNP intervention. We observed the greatest hepatic lipid accumulation in the CE fraction of LAL^−/−^ livers with a drastic 150-fold increase, whereas TGs were elevated only 2-fold. These changes are in accordance with TG and total cholesterol concentrations in the liver of 4-week-old LAL^−/−^ mice (33) as well as adult fed and fasted LAL^−/−^ mice (55), emphasizing the crucial role of LAL particularly in CE metabolism. CEs and TGs were largely enriched with saturated (C18:0, C16:0) and unsaturated (C18:2n6, C18:1n9, C16:1n7) FAs, all of which have been previously associated with pro-apoptotic properties and involvement in the pathogenesis of metabolic dysfunction-associated steatotic liver disease (MASLD) and metabolic dysfunction-associated steatohepatitis (MASH) (36, 56, 57). The significant reduction in accumulation of these lipotoxic lipids by F23 administration and the concomitant decline in n-6 unsaturated FA and AA concentrations suggests a pronounced hepatoprotective effect promoted by regaining LAL activity.

At the transcript level, F23 improved several hepatic pathways by alleviating LAL-D-associated pathological changes in gene expression profiles. The restored expression of genes involved in ß-oxidation suggests that the hepatic cells regained the ability to metabolize the FAs generated by LAL-mediated lipid hydrolysis. Consistent with these findings, the moderate but significant increase of PLs and FAs in the liver indicates that FAs are indeed mobilized by LAL, but concomitant rescue of FA ß-oxidation limits the accumulation of its lipotoxic by-products (58, 59). These lipidomic and transcriptomic profiles have shown that LAL and its hydrolytic products are key regulators of numerous pathways and homeostatic processes and that their reactivation by mRNA-LNPs represents a promising possibility for future therapeutic avenues.

Some limitations of our study, however, need to be mentioned. First, LAL^−/−^ mice are a genetic model of WD and a pathological model of CESD, so unlike WD patients, they do not suffer from early lethality. Second, due to the degradation of transfectants by phagocytes, a more comprehensive evaluation of the dosage, dosing interval, and duration of treatment needs to be performed, as these factors strongly affect the efficacy of gene therapy (19, 60-62). Third, the lack of body weight gain after administration of mRNA-LNPs and other treatment modalities, including ERT and AAV-based approaches, highlights the great need for a better understanding of the systemic, intestinal, or other extrahepatic mechanisms that drive this pathologic trait of LAL^−/−^ mice (34, 44, 54).

In summary, our study showed that the development and optimization of mRNA-LNP from a combinatorial library yielded a formulation capable of safely and efficiently delivering mRNA encoding LAL to partially restore its function *in vitro* and *in vivo* in target organs. Long-term administration of this formulation significantly reduced lipid accumulation, attenuated the progressive disease phenotype, and restored crucial metabolic functions in the liver of a preclinical model of LAL-D. We conclude that these findings pave the way for investigating and improving novel treatment strategies to overcome the limitations of current therapeutic approaches, and offer perspectives for targeted therapies that could transform the outcome for LAL-D patients.

## MATERIALS AND METHODS

### Lysosomal acid lipase enzymatic activity assay

LAL activity was determined using the fluorogenic substrate 4-methylumbelliferone oleate (4-MUO) (63). with minor modifications. Briefly, 5 μL of protein lysates (1 μg/μL) were mixed with 75 μL of 200 mM sodium acetate buffer, pH 5.5, and preincubated for 10 minutes in the presence or absence of Lalistat-2 (final assay concentration 0.2 μM) in DMSO. 25 µL of substrate (4-MUO 1 mg/mL in 4% Triton X-100) was added, incubated for 30 minutes at 37 °C, and the reaction was stopped with 50 μL of 1 M Tris pH 8.0. Fluorescence (λex = 360 nm, λem = 460 nm) was measured on a Spark microplate reader (Tecan, Maennedorf, Switzerland) and LAL activity was calculated as the difference between triplicates incubated with and without Lalistat-2. Relative fluorescence values were normalized to the controls (siCtl- or siLIPA2-treated cells, PBS- or LNP Blank-treated mice).

### Galectin 8-GFP reporter cell model and confocal imaging

The Galectin 8-GFP (GFP-Gal8) reporter fusion model was achieved in HepG2 cells by adapting previously described protocol (29). Briefly, Piggybac transposon plasmid encoding GFP-Gal8 fusion protein (#127191, Addgene) was co-transfected with a Super Piggybac Transposase expression vector (#V012800, NovoProlabs). GFP-positive cells were selected at 8 days post-transfections by fluorescence-activated cell sorting using a FACSAria™ IIIU Fusion (BD Biosciences, Franklin Lakes, NJ) and BD FACS Diva v8.0.1 software. For confocal imaging, 30,000 HepG2 cells expressing Gal8-GFP were plated out 24 hours before image acquisition in an Ibidi µ-Slide 8-well high glass bottom plate. Cell nuclei were stained with Hoechst (H3570, Invitrogen, Waltham, MA) prior the addition of 200 µL of DMEM and 100 µL of F01 or F23 (0.5ug h*LIPA* mRNA/well). Images were taken at 5 focal planes per view at hourly intervals for 17 hours using an LSM700 laser confocal microscope (Zeiss) and quantified with ImageJ software (NIH, version 1.54h and 1.54i).

### Animal studies

Wild-type CD-1 mice (6-8 weeks of age) were purchased from Charles River (Montreal, QC) and housed under a specific-pathogen-free conditions with a 12-hour light / dark cycle and maintained on standard rodent chow (Teklad global, 18% protein rodent diet) with free access to water. All experiments with CD-1 animals were conducted in accordance with the protocols approved by the Ethics Committee for Animal Experiments of the Université de Montréal (appr. 23-001).

Kinetic and efficacy studies in LAL^−/−^ mice were performed according to the European Directive 2010/63/EU in compliance with national laws and approved by the Austrian Federal Ministry of Education, Science and Research, Vienna, Austria (2020-0.129.904). LAL^−/−^ mice were generated using embryonic stem cells from EUCOMM (*Lipa^tm1a(EUCOMM)Hmgu^/Biat).* The *Lipa-tm1a* mice were crossbred with transgenic mice expressing Cre recombinase under the control of the cytomegalovirus promotor, resulting in *Lipa-tm1b^+/−^*mice. These mice were intercrossed to generate LAL^−/−^ mice and their WT littermates. Male and female mice were enrolled in the studies at equal numbers.

### Efficacy studies in LAL^−/−^ mice

Female and male LAL^−/−mice^ aged 4-9 weeks received either F23 mRNA-LNPs (1.0 μg RNA/g BW) or empty F23 LNPs and WT littermates received empty F23 LNPs via retro-orbital injection every 5 days over a period of 2 months, totaling 13 injections. BW was recorded before each injection. One day after the last injection, the mice were fasted for 4 hours, and euthanized. Blood was drawn and plasma was isolated by centrifugation at 3500 rpm for 10 minutes at room temperature. Livers and spleens were thoroughly rinsed in ice-cold PBS and snap-frozen for further analyses or fixed in 4% neutral-buffered paraformaldehyde for 24 hours for histological preparations.

### Transcriptomics

cDNA libraries were prepared using the Kapa mRNA Hyper Prep kit (KapaBiosystem) according to the manufacturer’s instructions, and sequenced on an Illumina NextSeq 500 to generate ∼24 million 84-bp single-end reads per sample. Trimmomatic was used for adapter trimming and removal of low-quality 3’ bases. Processed reads were mapped to the GRCm39 mouse reference genome via STAR using gene annotations from GENCODE M29. Normalization and differential gene expression analyses were done using the DESeq2 R package (v1.30.1). G:Profiler web-based software served for functional gene enrichment analysis based on gene sets from the Reactome pathway database. Gene set enrichment analyses (GSEA) were done with the GSEA software (v. 4.3.0) using fold change (Log_2_) values from DESeq2 R package. Hallmarks, Reactome and GoBP MSigDB gene set collections were used in the analysis. Genes ranked with a positive or negative enrichment were selected and the 15 genes with the highest fold change (Log_2_) were then computed.

### Targeted lipidomics

Pulverized tissues were incubated overnight at 4°C in a solution of chloroform/methanol (2:1) containing 0.004% butylated hydroxytoluene (BHT), filtered and dried under nitrogen gas. Cholesteryl esters and triglycerides were eluted on an aminopropyl column (Bond Elut LRC-NH2, 500mg) (Agilent Technologies). Fatty acids from each fraction were analyzed as their methyl esters after a direct transesterification with acetyl chloride/methanol on a 7890B gas chromatograph coupled to a 5977A Mass Selective Detector (Agilent Technologies, USA) equipped with a capillary column (J&W Select FAME CP7420; 100 m x 250 µm inner diameter; Agilent Technologies Inc.) and operated in the PCI mode using ammonia as the reagent gas. Samples were analyzed under the following conditions: injection at 270 ⁰C in a split mode (split ratio: 50:1) using high-purity helium as the carrier gas (constant flow rate: 0.44 mL/min) and the following temperature gradient: 190 ⁰C for 25 min, increased by 1.5⁰C/min until 236⁰C. Fatty acids were analyzed as their [M+NH3]+ ion by selective ion monitoring and concentrations were calculated using standard curves and isotope-labeled internal standards. PCA was performed on a web-based software PCA calculator https://www.statskingdom.com/pca-calculator.html.

### Statistical analysis

Data were analyzed and presented using GraphPad Prism (8.4.0). Principal component analysis was performed by plotting each value for PC1 and PC2 in a scatter dot plot in Prism. Figures show mean ± SEM. Unpairwise comparison of means between two groups was performed using Student’s *t*-tests (two sided) with the assumption of normal distribution of the data. Multiple groups were compared using one-way analysis of variance (ANOVA) with Tukey’s post hoc correction. Values and error bars are expressed as means ± SEM. Degrees of significance were defined as *P* < 0.05 (\**P* < 0.05, \*\**P* < 0.01, and \*\*\**P* < 0.001).

## Supporting information

Supplementary Informations

## Acknowledgments

We would like to thank the Metabolomic core facility (Montreal Heart Institute), notably Prof. M. Ruiz, C. Daneault, and I. Robillard, for performing the targeted lipidomic analysis; the genomic core facility at the Institute for Research in Immunology and Cancer in Montreal for performing the RNAseq analysis. We greatly appreciate the assistance of Nassim Oumansour (Faculty of Pharmacy, University of Montreal) for characterization of the LNP library, B. Schwarz (Gottfried Schatz Research Center, Medical University of Graz) for ORO and H&E staining, and T. Moustafa (Division of Gastroenterology and Hepatology, Medical University of Graz) for Sirius red staining.

## Funding

This work was supported by:

Phospholipid Research Council grant DBR-2022-102/1-1 (DB, SPG)

Canadian Institutes of Health Research grant PJT 190141 (DB, SPG)

Natural Sciences and Engineering Research Council of Canada grant RGPIN-2018-05076 (DB)

Natural Sciences and Engineering Research Council of Canada grant RGPIN-2023-03828 (SPG)

Natural Sciences and Engineering Research Council of Canada Alliance International Catalyst Grant (ALLRP) grant ALLRP 586015 – 23 (DB, SPG)

Canadian Generic Pharmaceutical Association and Biosimilars Canada Biotherapy Chair (DB)

FRQS research scholarship grant number 282186 (DB)

FRQS doctorate student scholarship number 29869 (MZ)

Canada First Research Excellence Fund through the TransMedTech Institute (MZ)

Austrian Science Fund (FWF) grants SFB 10.55776/F73 and 10.55776/P32400 (both to DK)

Province of Styria (DK)

City of Graz (DK)

The authors sincerely acknowledge the financial support from the Faculty of Pharmacy at the Université de Montréal

## Author contributions

Conceptualization: SPG, DB, DK designed and conceptualized the study.

Methodology: DB, SPG, DK, NV, MZ designed the experimental work, HH performed the Gal8-GFP experiments, image acquisition and analysis.

Investigation: MZ, performed most of the experimental work, ELV installed, optimized the microfluidic and assisted in *in vitro* experiments and for the synthesis of the ionizable lipid 5A2-SC8, NV performed retro-orbital injection and assisted in all experiments in LAL^-/-^ mice, SB performed the synthesis and characterization of 5A2-SC8, RMKR assisted with RNA extraction of in vivo samples

Visualization: MZ created all the figures, HH assisted in the visualization of confocal images in the Gal8-GFP experiments, NR and FM assisted in the data visualization.

Funding acquisition: DB, SPG, DK acquired funding

Project administration: DB, SPG, DK. DK managed ethical approval for experiments in LAL^-/-^ mice

Supervision: DB, SPG oversighted the planning, the execution of the study and managed the collaboration with the external core facilities.

Writing – original draft: MZ wrote the original draft with the help of the coauthors.

Writing – review & editing: DB, DK, SPG, NV reviewed and corrected the manuscript.

## Data and materials availability

Numerical values of the mRNA-LNPs library and lipidomic analysis can be found in the supplementary information. RNA-seq dataset is available on NCBI Gene Expression Omnibus, accession number GSE280933. All data associated with this study are available in the main text or the supplementary materials.

## Supplementary Materials

### Materials and Methods

#### Cell culture

HepG2 cells (ATCC; HB 8065, passages 10–25) were maintained in high-glucose Dulbecco’s modified Eagle’s medium (DMEM plus sodium pyruvate, supplemented with 10% fetal bovine serum (FBS) and 1% penicillin/streptomycin. Cells were harvested using trypsin (all reagents from Wisent) at 70-90% and reseeded to 50% confluency.

#### RNA extraction from cells and tissues

Total RNA from cells was isolated using Monarch® Total RNA Miniprep Kit (New England Biolabs, Ipswich, MA) following manufacturer’s protocol. Total RNA from ∼20 mg of tissues was isolated by homogenization in TRIzol reagent (Millipore-Sigma, Burlington, MA) using the TissueLyzer® II, extracted as described (64), and resuspended in diethyl pyrocarbonate (DEPC)-treated water.

#### Reverse transcription quantitative real-time PCR

Reverse transcription was performed with 1 µg of RNA using the LunaScript RT SuperMix Kit (E3010, New England Biolabs, Ipswich, MA) on a SimpliAmp thermal cycler (A24812, Applied Biosystems) according to the manufacturer’s protocol. Primers used for the amplicon of the encapsulated mRNA (hLIPA) were specific to the FLAG sequence: Forward (Fwd) primer 5’-CTGGGGAAGCAGTGCCAAGAATTA-3’, Reverse (Rv) primer 5’-GTCGTCATCGTCTTTGTAGTCCTG-3’. For the endogenous human LIPA (LIPA) in HepG2 cells, the 3’UTR region was amplified using the Fwd primer 5’-CTCTGGACCCTGCATTCTGA-3’ and Rv primer 5’-AGACAACTGGTTTGGGACCTTTG-3’. Quantitative RT-PCR was performed with the Luna Universal qPCR Master Mix (M3003, New England Biolabs) on a QuantStudio 5 Real-time PCR system (A28133, Applied Biosystems). Fold change expression levels were calculated using the 2-ΔΔCt method and normalized to endogenous human TAT-Box Binding Protein (TBP) for HepG2 cell extracts (Fwd: 5’-TGCCACGCCAGCTTCGGAGA-3’, Rv: 5’-ACCGCAGCAAACCGCTTGGG-3’, and to mouse TBP for in vivo experiments (Fwd: 5’-CTGGCGGTTTGGCTAGGTTT-3’, Rv: 5’-ACCATGAAATAGTGATGCTGGG-3’.

#### Protein extraction from cells and tissues

Cells were harvested by scraping in Triton X-100 lysis buffer (150 mM NaCl, 50 mM Tris-HCl pH 7.4, 5mM EDTA, 1mM EGTA, 1% Triton X-100, with protease inhibitor cocktail (PIC; 1:100 dilution; ab65621, Abcam, Cambridge, UK), NaF (5 mM), and Na3VO4 (1 mM) added before use), left on ice for 30 minutes with occasional vortexing, centrifuged (17,000 x g for 20 minutes at 4 °C), and the infranatants were stored at −80 °C until use. The snap-frozen tissues were pulverized using a liquid N2-cooled mortar and pestle, weighed (∼20 mg), and homogenized in Triton X-100 lysis buffer using the TissueLyzer® II equiped with steel beads (69997; Qiagen, Hilden, Germany) for 2 × 30 Hz, with 30 s per cycle. The homogenized samples were incubated on ice for 30 minutes with occasional vortexing. Total protein content of the lysates was determined using the Pierce microBCA protein assay kit (23235, Thermo Fisher Scientific, Waltham, MA), following the manufacturer’s instructions.

#### Immunobloting

Fifteen and thirty micrograms of proteins for in vitro and in vivo experiments, respectively, were separated on a 10% SDS-PAGE, transferred onto polyvinylidene difluoride membranes (PVDF; Millipore-Sigma, Burlington, MA) using a semi-dry Trans-Blot SD (Bio-Rad, Hercules, CA), blocked, and incubated with anti-LAL (sc-58374, clone 9G7F12, Santa Cruz Biotechnology, Dallas, TX) or anti-FLAG antibody (Millipore-Sigma, F7425) both at a 1:1000 dilution overnight at 4 °C. The following day, the membranes were incubated for 1 hour at room temperature with the corresponding horseradish peroxidase-conjugated secondary antibodies (KP-5220-0367, KP-5220-0336, Mandel scientific, Guelph, ON) and incubated with the SuperSignal West Pico PLUS Chemiluminescent substrate (34579, Thermo Fisher Scientific). Chemiluminescence was detected on Blu-Lite films (A8815, Dutscher, Bernolsheim, France). Vinculin (14-9777-82, clone 7F9, Thermo Fisher Scientific) expression was determined as a loading control.

#### Nile red staining of neutral lipids

HepG2 cells were seeded in 24-well plates (75,000/well) and incubated for 24 hours prior to staining with a commercially available Nile red kit (500001, Cayman chemicals, Ann Arbor, MI) and DAPI (1 μg/mL, Millipore-Sigma). The fixed cells were imaged with a 20x objective on an Axio Observer Zeiss Z1 microscope (Zeiss, Oberkochen, Germany) and quantified using ImageJ software (NIH, version 1.54h and 1.54i).

#### Small interfering RNA treatments in HepG2 cells

At a density of 150,000 cells per well, HepG2 cells were plated in 6-well plates for 24 hours prior the transfection with 10 nM of siLIPA1 (#SI00005082), siLIPA2 (#SI00005103) (both from Qiagen) or siCtl (#1027292,AllStars, Qiagen), and 5 µL of RNAiMAX (Thermo Fisher Scientific) following the manufacturer’s protocol. For screening of the mRNA-LNPs, 1,800,000 cells were plated in a 10-cm2 petri dish, and transfection was performed with 15 μL of siLIPA2 and 30uL of lipofectamine RNAiMax following an adapted version of the manufacturer’s protocol.

#### h*LIPA* mRNA and Nanoparticle formulation

hLIPA mRNA was synthesized at TriLink Biotechnology (San Diego, CA, USA) and included the coding region (NM001127605.3) flanked with the 5’UTR and 3’UTR optimized for mRNA translation, the poly-adenosine tail at the 3’end, a N7-methyl guanosine and adenosine (m7GpppAG) cap, and 5-methoxyuridine as modified base. A C’ termina FLAG sequence was added prior the STOP codon. DLin-MC3-DMA (Caymans chemicals), DMG-PEG (Millipore-Sigma), cholesterol (Millipore-Sigma) DSPC, DPPC, DOPE, DMPC, HSPC, DOPC (all were from Lipoid) and stock solutions were prepared in RNAse-free absolute ethanol. 5A2-SC8 ionizable lipid was synthesized and characterized as previously described (20), The stock solutions of each lipid were pipetted and mixed at the given molar ratio listed in Table S3. mRNA was diluted in sodium acetate buffer (25 mM, pH 4.0). The molar ratio of the amine groups of the ionizable lipid to the phosphate groups on mRNA (N/P ratio) was kept at 8 for all the formulations of the library. Ethanol and aqueous phases were mixed using a microfluidic polycarbonate herringbone chip (FLUIDIC 187, Chipshop, Germany) with the aid of a pressure flow controller (OB1 MK3+, Inside Therapeutics, France) to generate the LNPs. For all the formulations, the pressure of the organic phase and of the aqueous phase were kept at 400 mbar and 480 mbar, respectively. For in vitro experiments, LNPs formulations were diluted with 1X RNAse-free PBS to 0.5 µg hLIPA mRNA for transfection into HepG2. For animal experiments, mRNA-LNP formulations were dialyzed (Pur-A-Lyzer Midi Dialysis Kits, WMCO 3.5kDa, Thermo Fisher Scientific) against 1X RNase free PBS for 1h then diluted with PBS for i.v. injection.

#### LNP characterization

LNP size was assessed by dynamic light scattering (Zetasizer Nano ZS; Malvern Instruments, Worcestershire, UK). The surface charge was measured by zetametry (NanoBrook Omni; Brookhaven instruments, Nashua, NH). The encapsulation efficiency was determined using the Quant-iT RiboGreen RNA Assay (R11490, Thermo Fisher Scientific) based on a modified protocol to allow measurements in a 96-well plate. Briefly, 10 μL of LNP formulation were diluted (1:10) in 10 mM Tris-HCl, 1 mM EDTA, pH 7.5 (TE) buffer and either 90 μL of TE buffer or 2% Triton X-100 (Tx) were added and the samples were incubated for 10 minutes at 37 °C. Ribogreen solution was added to the samples and incubated at room temperature for 10 minutes andfluorescence was assessed with a microplate reader (TECAN model SPARK, Switzerland) at λexc = 485 nm, λem = 528 nm.

#### Cell viability

HepG2 cells were seeded at a density of 15,000 cells/well in a 96-well plate. After 24 hours, cells were transfected for 24 hours with either blank or mRNA-loaded LNPs at the desired lipid concentration. After 30 minutes of incubation with PrestoBlue (Thermo Fisher Scientific, A13261), the fluorescence (λexc = 560 nm, λem = 590 nm) was measured using a Spark microplate reader (Tecan).

#### Biodistribution and pharmacokinetics studies

DiIC18(5); 1,1′-dioctadecyl-3,3,3′,3′-tetramethylindodicarbocyanine, 4-chlorobenzenesulfonate salt (DiD) (D7757, Thermo Fisher Scientific) was incorporated into the selected mRNA-LNP formulations at a 0.2 molar ratio, and CD-1 mice (3−4 per group) were injected with either 0.5 μg mRNA /g body weight (BW) or PBS through the tail vein. Blood was collected at 0.08, 0.5, 1, 2, 4, 8, 24, and 48 hours and fluorescence (λexc/em: 630/675 nm) in the plasma was measured with a Spark microplate reader (Tecan). Noncompartmental pharmacokinetic parameters were calculated using Kinetica software (version 5.1 SP1, Thermo Fisher Scientific). Animals were euthanized after 48 hours, the organs were collected and imaged with the OiS300 Lighttrack in vivo imaging system (Labeo Technologies Inc., City, Canada) using a Cy5.5 filter (λexc/em: 655/716 nm).

#### Quantification of hLIPA mRNA organ delivery

F01, F23, F27, and F50 were administered i.v. at 0.5 or 1.0 μg/g BW. WT mice were euthanized 2, 8, 24, and 72 hours, LAL−/− mice 24-, 48-, and 72-hours post-injection (3 animals per time point and formulation), along with corresponding PBS-injected controls. RNA was extracted from livers and spleens as described above. A qPCR calibration curve was generated by spiking known amounts of serially diluted exogenous hLIPA mRNA to tissue homogenates and the delivered hLIPA mRNA per mg of the pre-weighed tissue powder was extrapolated to the total amount of hLIPA mRNA per organ.

#### Cy5 labeling of h*LIPA* mRNA and flow cytometry

hLIPA mRNA was labeled with Cy5 using the Label IT Nucleic Acid Labeling Kit (MIR3100, Mirus Bio), encapsulated in LNPs, and injected i.v. at a dose of 0.5 μg/g BW mRNA. Controls were injected with PBS. After 2 hours, mice were perfused with PBS and the liver was isolated and digested in DMEM containing 0.15 U/mL Liberase TL (0540102000, Millipore Sigma) at 37°C under shaking for 45 minutes. The mixture was neutralized at 4 °C with HEPES buffer containing 2% FBS and 25 mM EDTA, the digest was passed through a 70-μm cell strainer and the red blood cells lysed using lysis buffer (00-4333, Thermo Fisher Scientific). One million cells were blocked with TruStain FcX™ PLUS (156603, clone S17011E, BioLegend), and distinct hepatic cell populations were sorted by flow cytometry using a BD LSRFortessa cell analyzer and BD FACS Diva v8.0.1 software. The following antibodies were used: Brilliant Violet 421 anti-CD31 (102424, clone 390), Alexa Fluor 488 anti-CD45.2 (103122, clone 104), Phycoerythrin/Cyanine7 anti-F4/80 (123114, clone BM8, all BioLegend), and Coralite 594 anti-ASGR-1 (CL594-11739, Proteintech). Live/Dead fixable Near IR (L34992, Invitrogen) was used to evaluate cell viability. Endothelial cells were defined as CD31+, CD45-, F4/80-, Kupffer cells as CD31-, CD45+, F4/80+, and hepatocytes as CD31-, CD45-, ASGR1+.

#### Oil red O (ORO), hematoxylin-eosin (H&E), and Sirius red (SR) staining

Paraformaldehyde-fixed tissues were paraffin-embedded for H&E and SR or sucrose-cryoprotected, OCT-embedded, and frozen-cut for ORO staining. Tissues were sectioned at 5-µm thickness, and staining was performed as previously described (65). The sections were imaged with an Olympus Slide View Digital Slide Scanner VS200 (Olympus Life Sciences, Tokyo, Japan) and quantified using QuPath-0.5.1 software.

#### ALT and AST measurement

Whole blood was collected in heparin tubes (450536, Greiner Bio one) and plasma samples were diluted 1:4 in PBS. Aspartate aminotransferase (AST) and alanine aminotransferase (ALT) measurements were performed on a Fujifilm Dry Chem NX500i machine (Fujifilm) and colorimetric multilayered slides (GOT/AST and GPT/ALT slides, Fujifilm).

**Figure S1.**
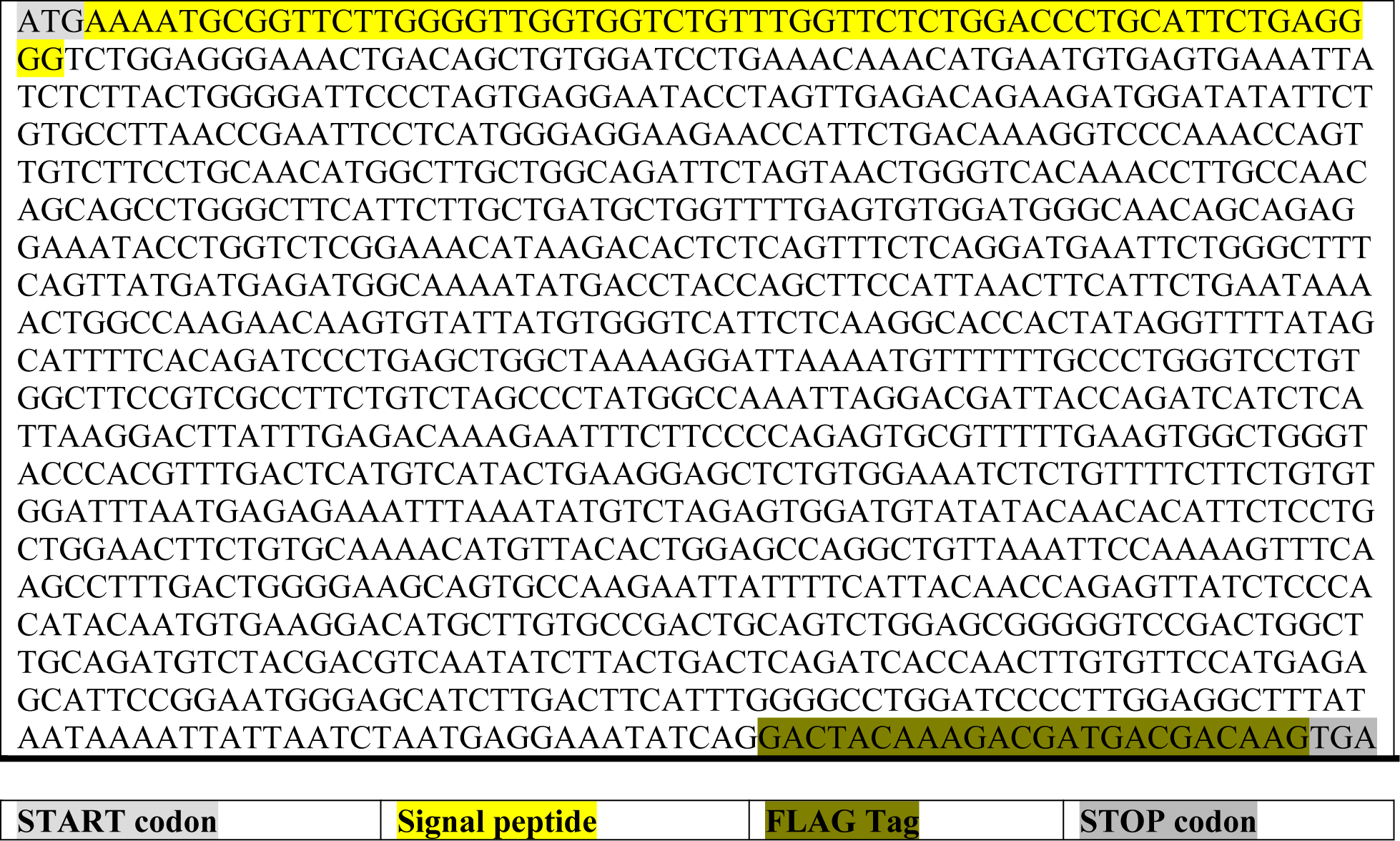
Designed exogenous h*LIPA* mRNA. LIPA lipase A, lysosomal acid type Homo sapiens (human) NM001127605.3 from the NCBI nucleotide database. A C-terminal FLAG TAG was added to distinguish between endogenous *LIPA* mRNA and encapsulated exogenous h*LIPA* mRNA.

**Figure S2.**
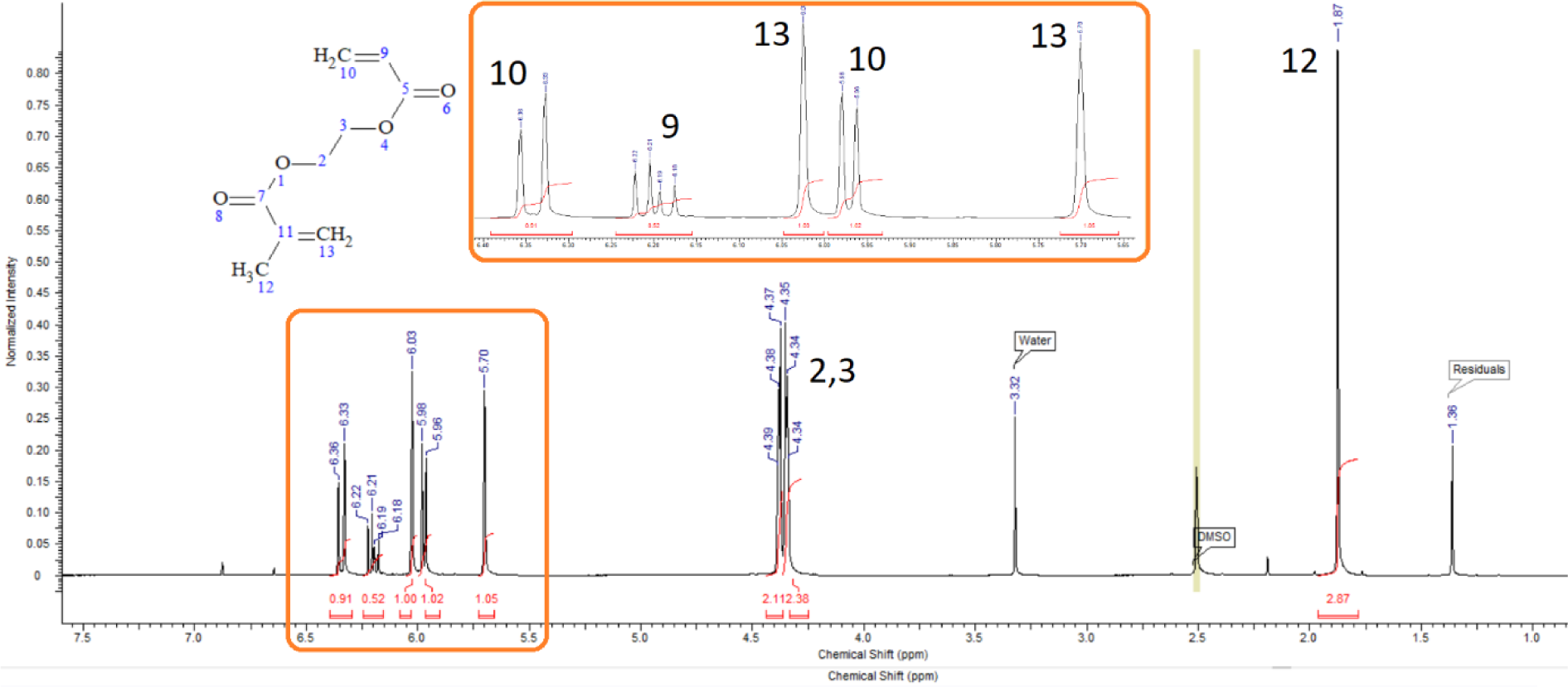
^1^H-NMR spectrum of AEMA (600 MHz, D6-DMSO, δ).

**Figure S3.**
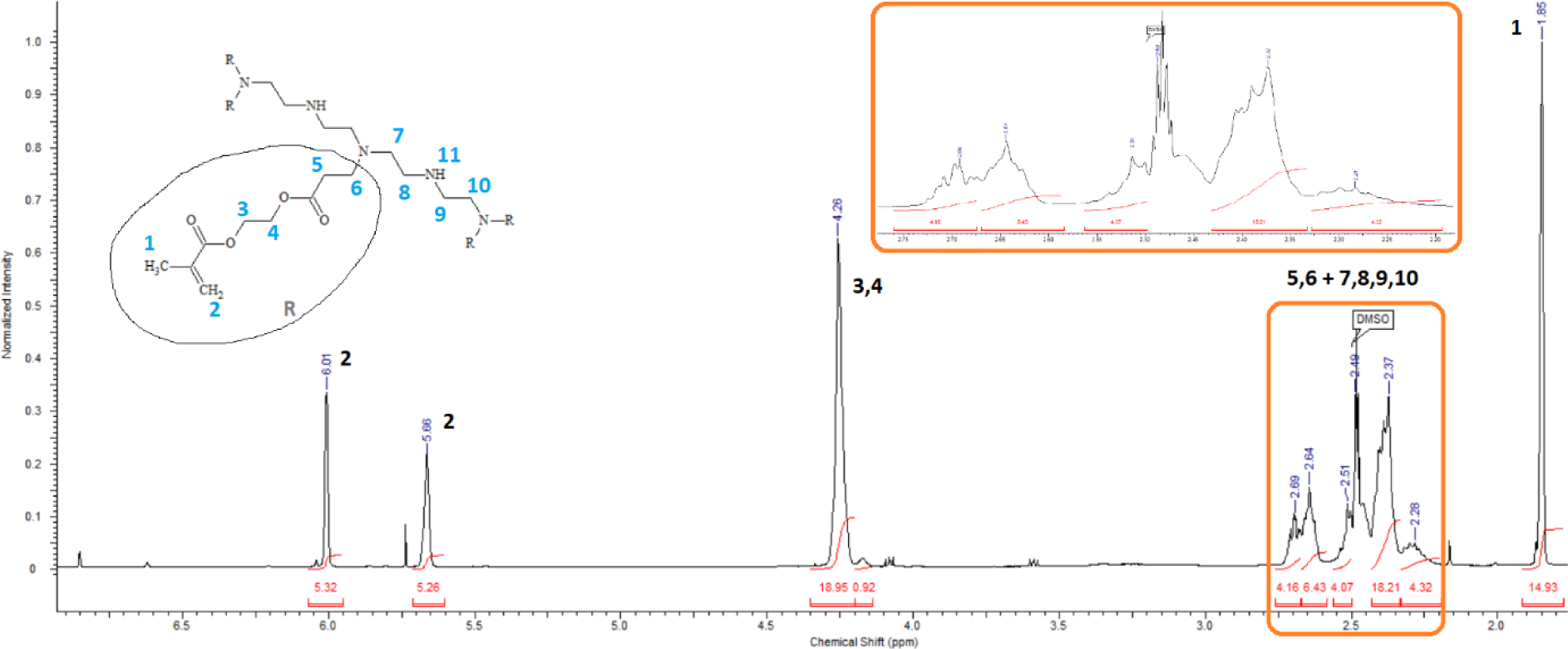
^1^H-NMR spectrum of the non-purified intermediate “5A2-meth” (600 MHz, D6-DMSO, δ).

**Figure S4.**
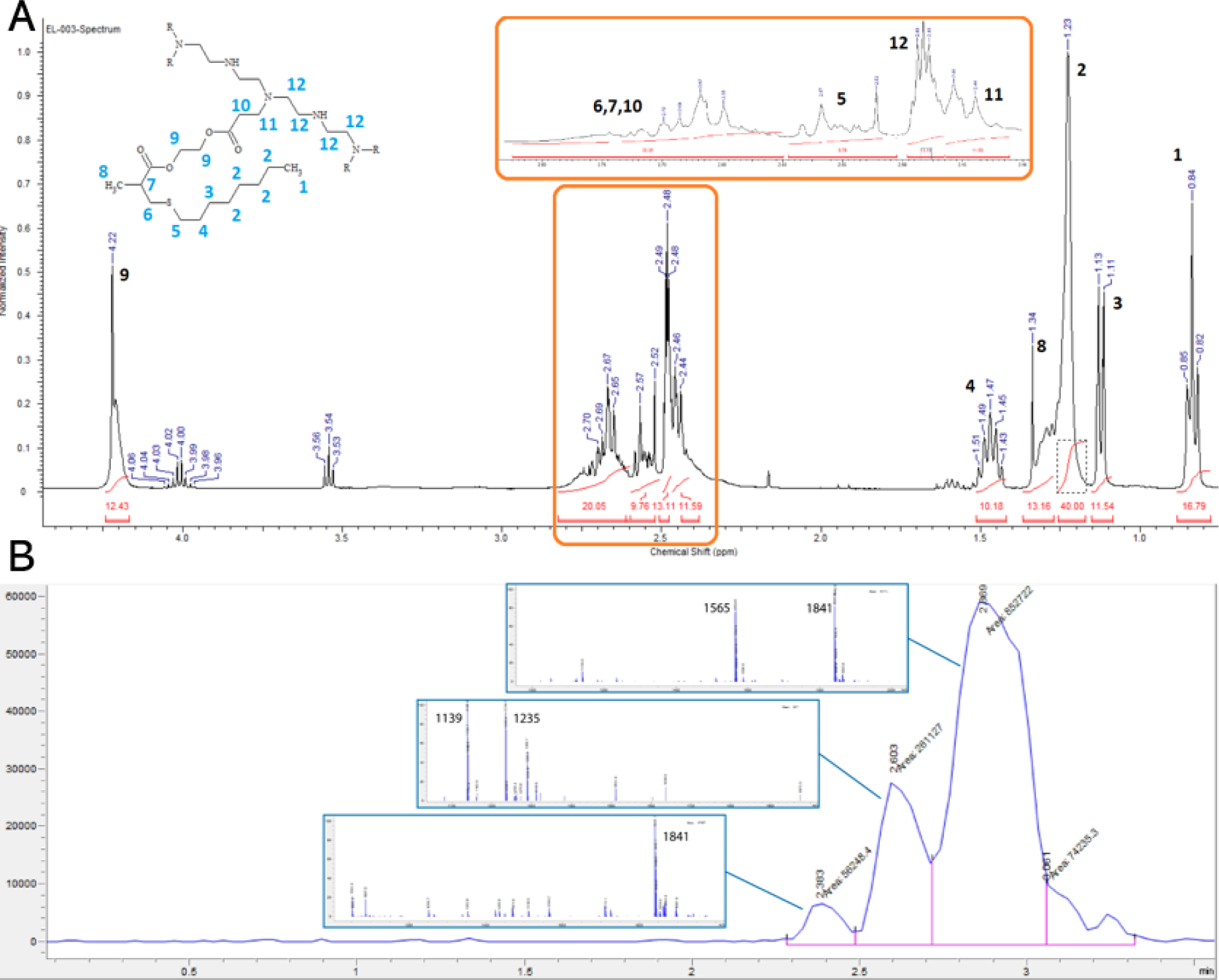
Characterization of the synthesized ionizable lipid 5A2-SC8. **(A)** ^1^H-NMR spectrum of the synthesized lipid 5A2-SC8 (600 MHz, D6-DMSO, δ) after purification. 4.22 (s, 12H, -OC*H2*C*H2*O-), 2.70-2.65 (m, 20H, -C*H*C*H2*SC8H17; -NCH2C*H2*COO-), 2.60-2.52 (m, 10H, -SC*H2*C7H15), 2.49-2.47 (m, 14H, -NC*H2*C*H2*NC*H2*C*H2*N-), 2.46-2.44 (m, 12H, - NC*H2*CH2COO-), 1.51-1.43 (c, 10H, -SCH2C*H2*C6H13), 1.34-1.25 (m, 14H, - OCOCH(C*H3*)CH2S-), 1.23-1.16 (s, 40H, -SCH2CH2CH2C*H2*C*H2*C*H2*C*H2*CH3), 1.13-1.11 (d, 12H, -SCH2CH2C*H2*C5H11), 0.85-0.82 (t, 17H, -SC6H12C*H3*). **(B)** LC spectrum of the purified fractions after column chromatography including associated MS spectrum for each peak with the specified m/z. 5A2-SC8 is expected to appear at m/z = 1841.

**Figure S5.**
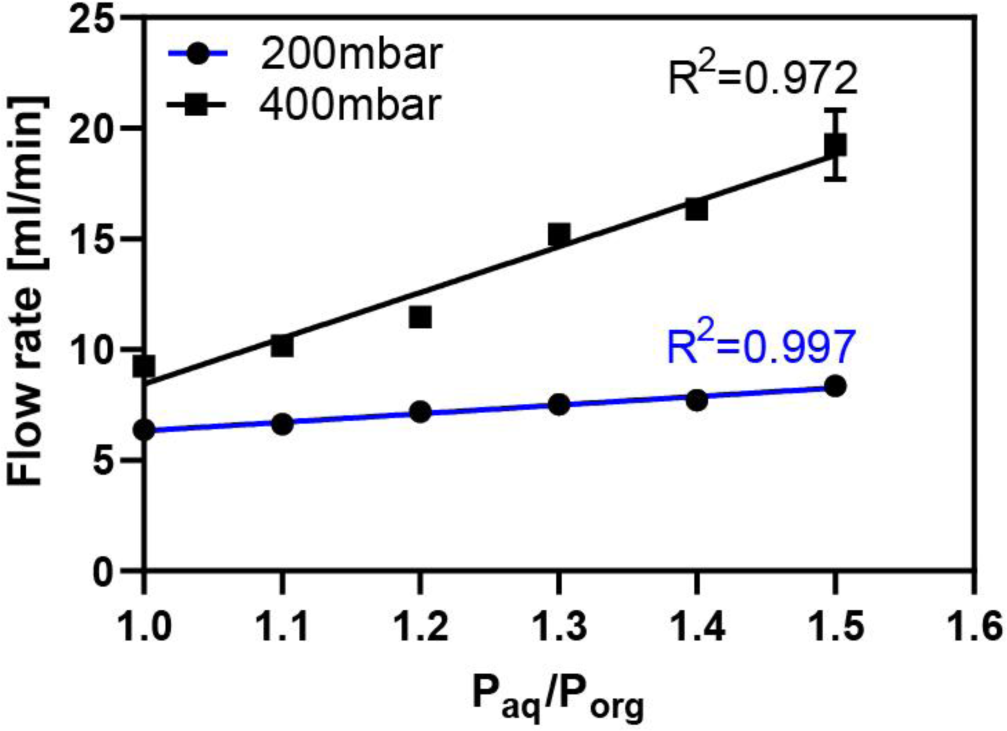
Linear relation between pressure ratio and flow rate. Increase of the ratio between the aqueous channel pressure (P_aq_) versus the pressure of the organic phase (P_org_) mediates an increase in the total flow rate in a linear relationship. The ratio was defined by maintaining P_org_ constant and increasing P_aq_.

**Figure S6.**
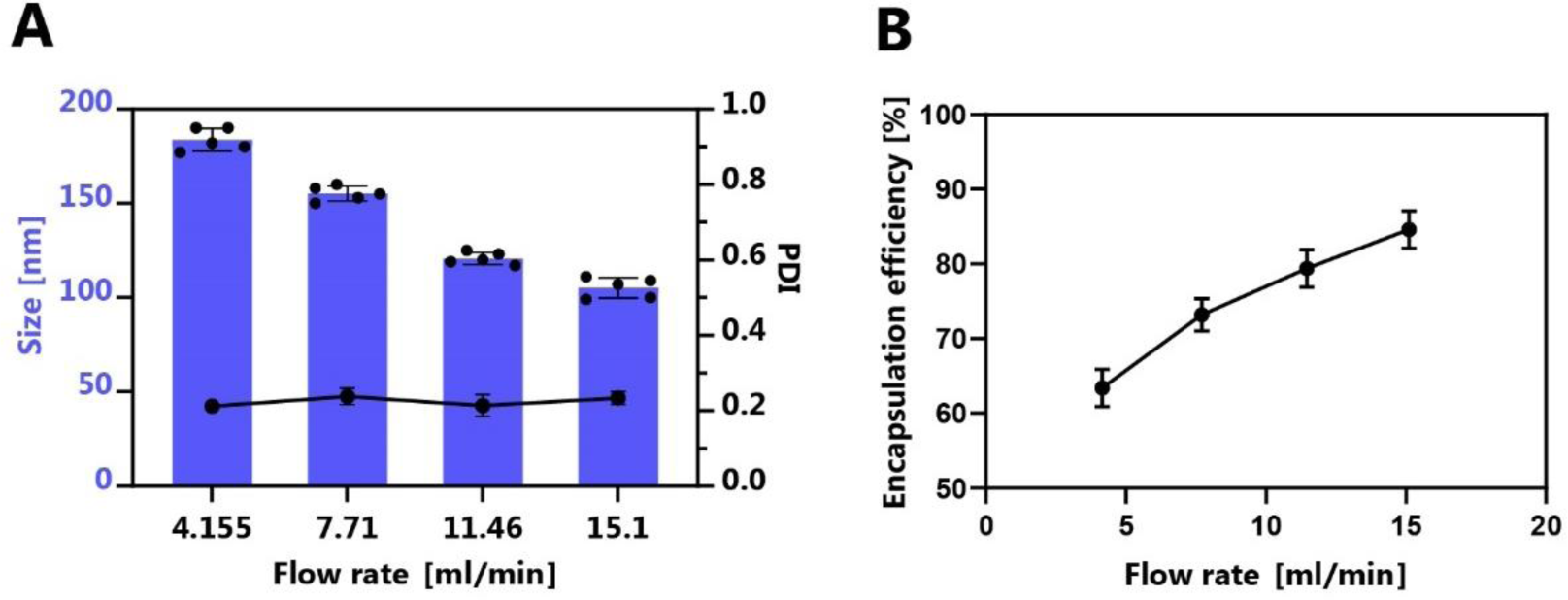
Optimization of the microfluidic setup. Dependence of the **(A)** hydrodynamic size (left axis) and the polydispersity index (right axis) as well as **(B)** the encapsulation efficiency on the flow rate of the microfluidic system. Data represent mean ± SEM of n=5 measurements.

**Figure S7.**
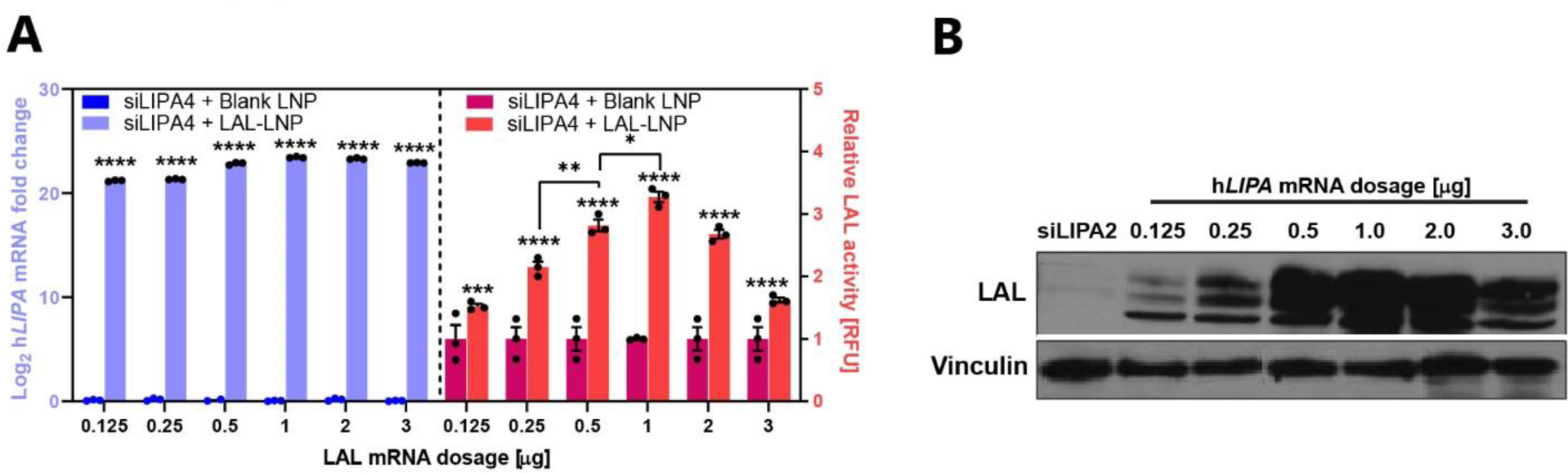
*In vitro* dose-response study with F01 formulation. **(A)** Exogenous h*LIPA* mRNA fold change in Log_2_ (left y-axis) and relative h*LIPA* mRNA-induced LAL activity (right y-axis) in HepG2 cells upon transfection with increasing amounts of encapsulated h*LIPA* mRNA. Data represent mean ± SEM of n=3 independent experiments. **(B)** Immunoblotting showing the gradual increase of the band intensity with increasing h*LIPA* mRNA dosage. Vinculin was used as a loading control.

**Figure S8.**
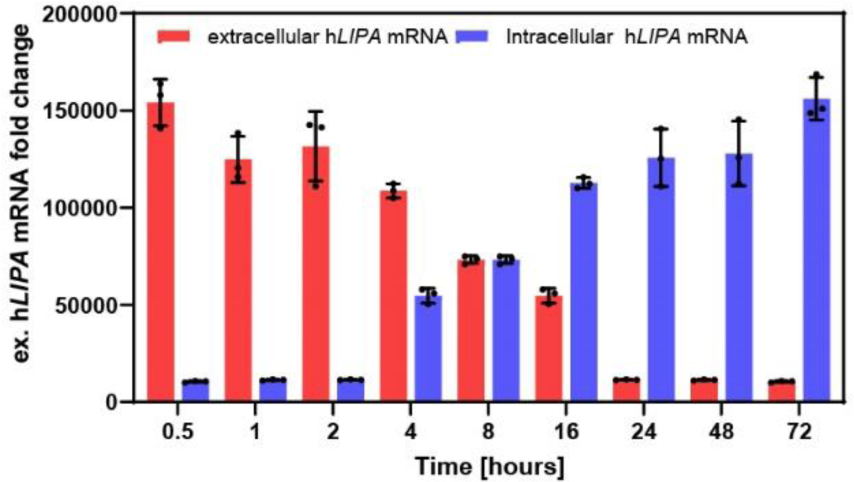
*In vitro* kinetic evaluation of encapsulated h*LIPA* mRNA delivery. F01 was transfected in HepG2 and the intracellular h*LIPA* mRNA levels were determined in the cell lysate and the medium at the indicated time points. The graph shows mean ± SEM of 3 independent experiments.

**Figure S9.**
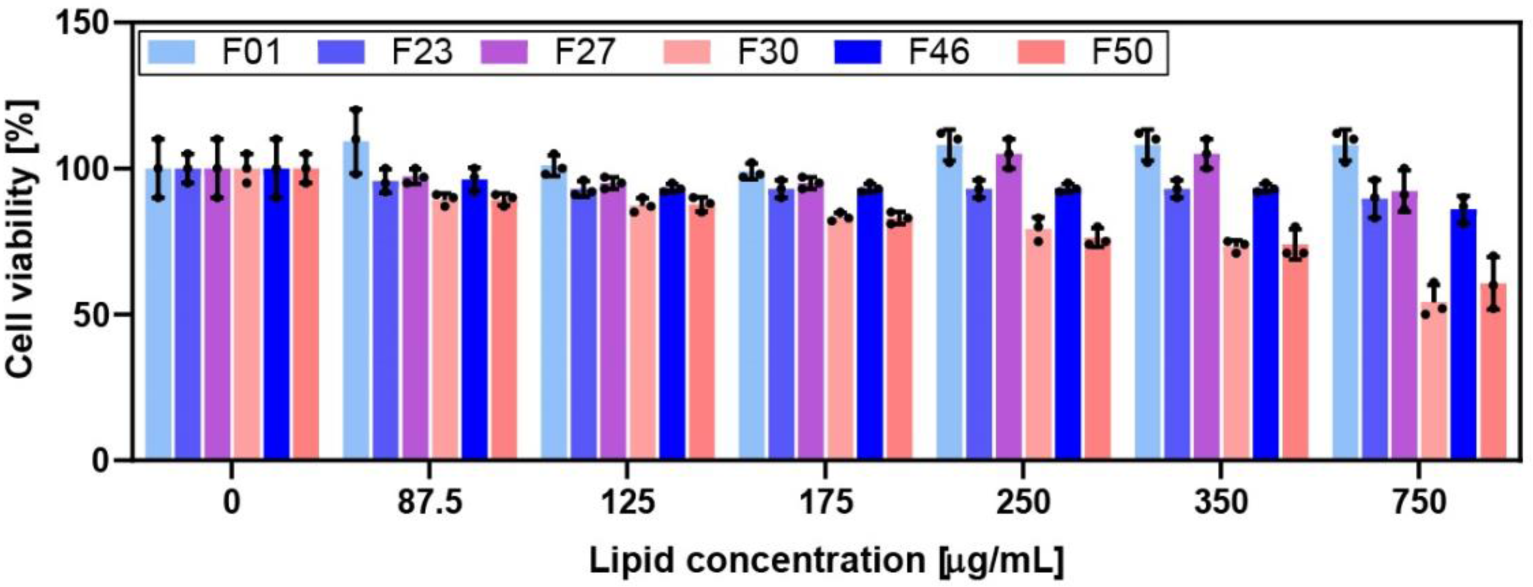
Cell viability of the tested formulations. Viability measurement 24 hours post-exposure of HepG2 cells to the mRNA-LNP formulations compared to cells grown in the absence of lipids. Data represent mean ± SEM of 3 independent experiments.

**Figure S10.**
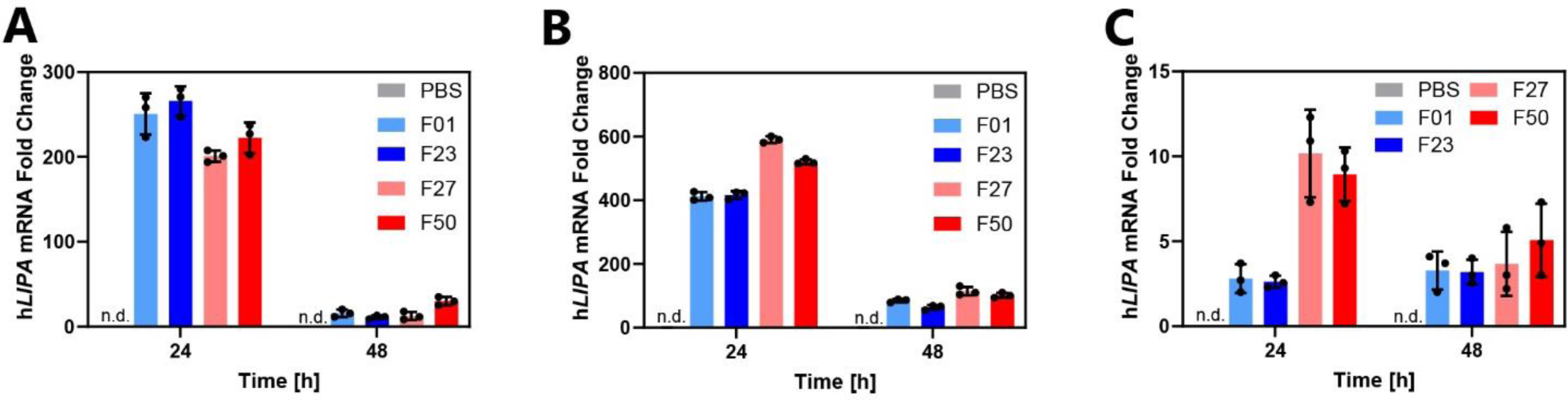
h*LIPA* mRNA levels delivered to tissues by selected formulations. **(A)** Exogenous h*LIPA*-mRNA delivered in the liver, **(B)** the spleen, and **(C)** in the lungs. Fold change have been normalized on murine TAT-binding protein (TBP). Data represent mean ± SEM (n = 3).

**Figure S11.**
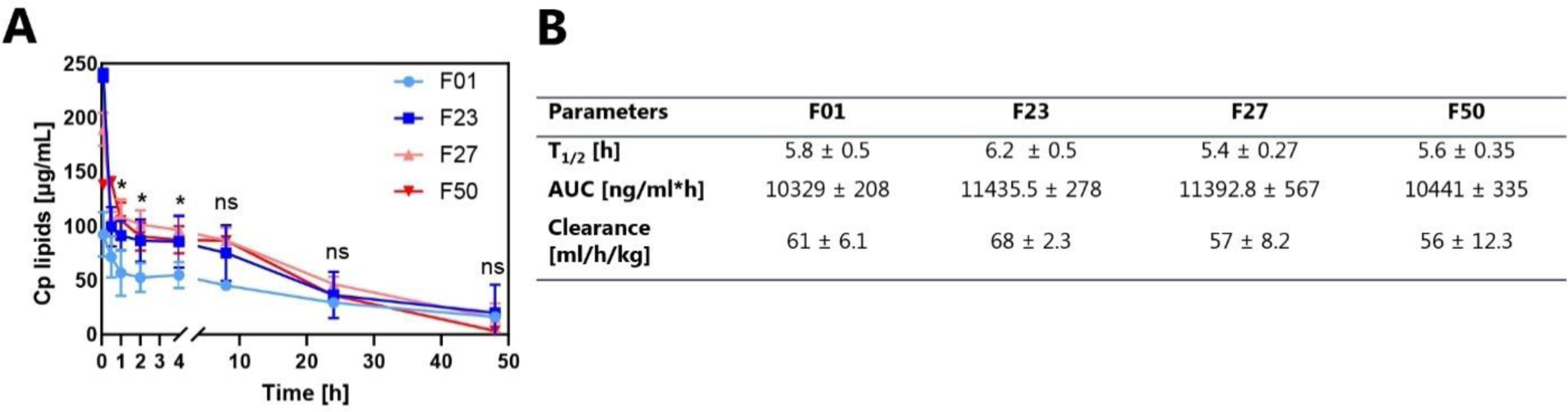
Pharmacokinetic analysis of the selected LNPs in healthy mice following i.v. administration. **(A)** Time-dependent changes of the plasma total lipid concentration (Cp) of the selected LNPs. **(B)** Pharmacokinetic parameters of the selected formulations. Data represent mean ± SEM (n = 3).

**Figure S12.**
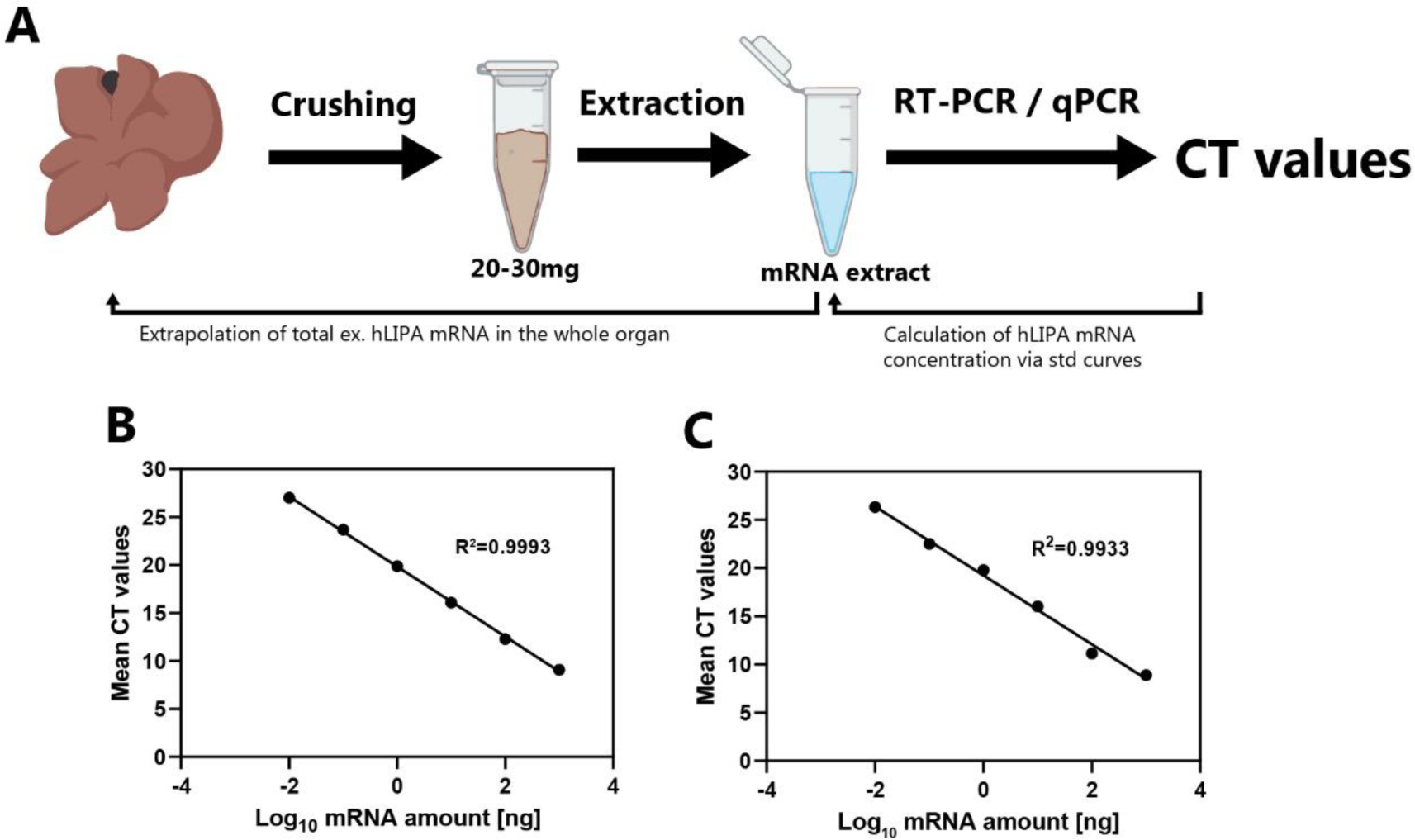
Quantification of the absolute amount of exogenous h*LIPA* mRNA delivered to the liver or the spleen. **(A)** Process workflow for the extrapolation of the absolute h*LIPA* mRNA amount delivered to the liver or spleen. **(B)** Standard curves with the cycle threshold (CT) values versus the h*LIPA* mRNA amount of the standards in logarithmic scale for liver and **(C)** spleen.

**Figure S13.**
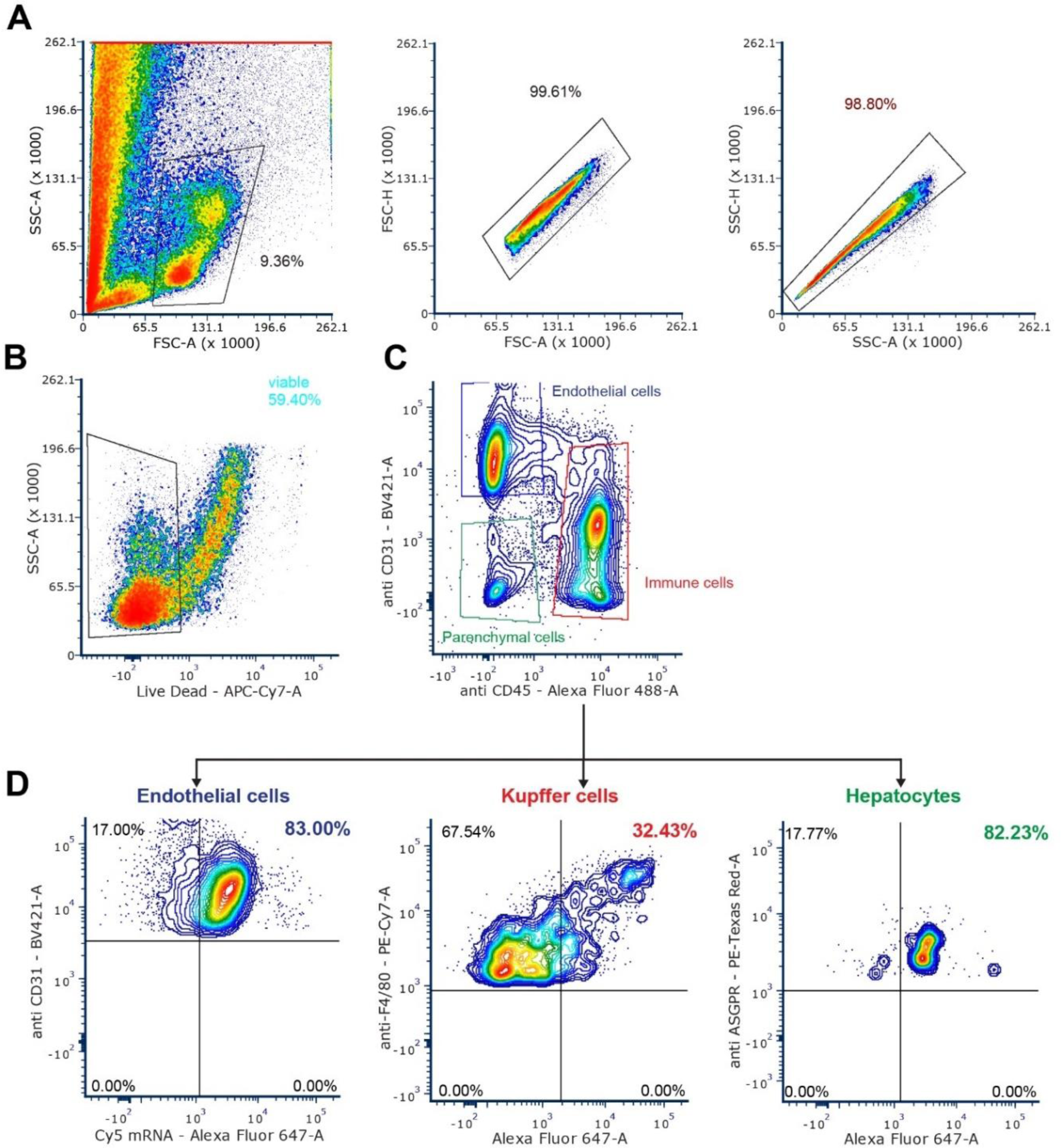
Flow cytometry gating strategy for immunophenotyping and transfection efficiency of F23 in hepatic cell subtypes. **(A)** All events recorded and subsequent isolation of single cells. **(B)** isolation of viable cells based on negative Live-dead staining. **(C)** preliminary separation of hepatic subpopulation defined as: immune cells CD31**^-^** and CD45^+,^ endothelial cells CD31^+^ and CD45^-^, parenchymal cells CD31^-^, CD45. **(D)** From **C**, Kupffer cells were specifically isolated as F4/80^+^ from the immune cells gate and hepatocytes as ASGPR^+^ from the parenchymal gate. In all subpopulation Cy5 mRNA signal was measured. In total n=3 mice.

**Figure S14.**
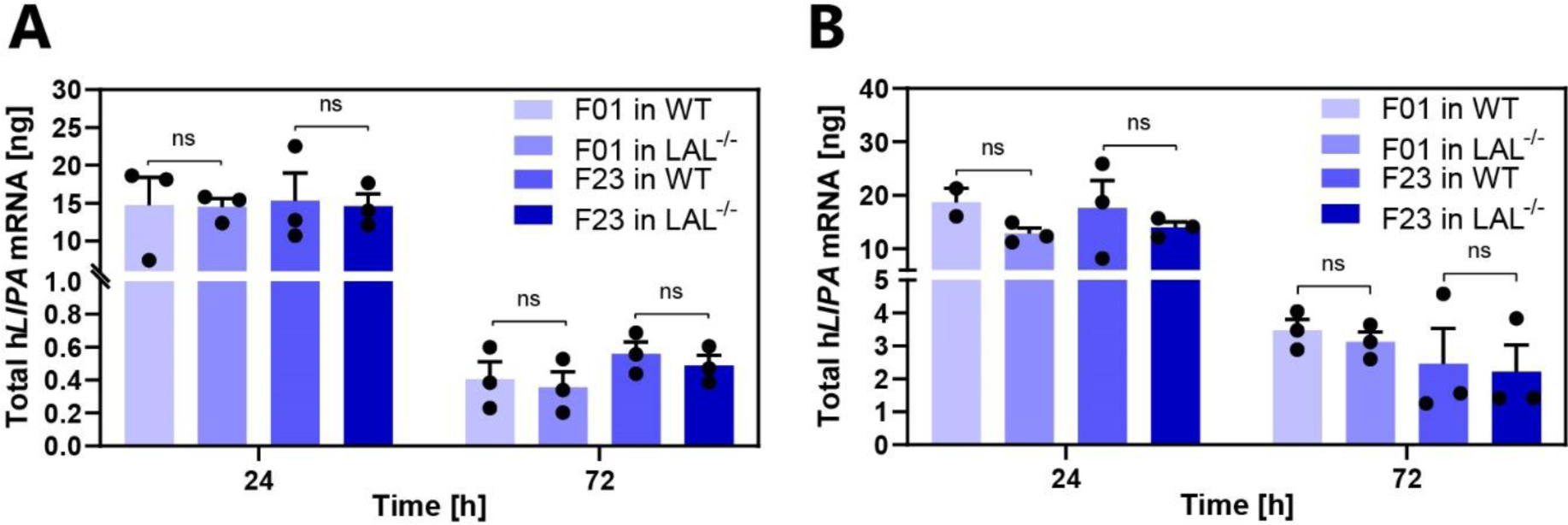
Comparison of mRNA delivery in WT and LAL^-/-^ mice. **(A)** mRNA levels in the liver and **(B)** spleen. Unpaired two-tailed Student’s *t*-test. Graphs show mean ± SEM (n=3). ns: non-significant.

**Figure S15.**
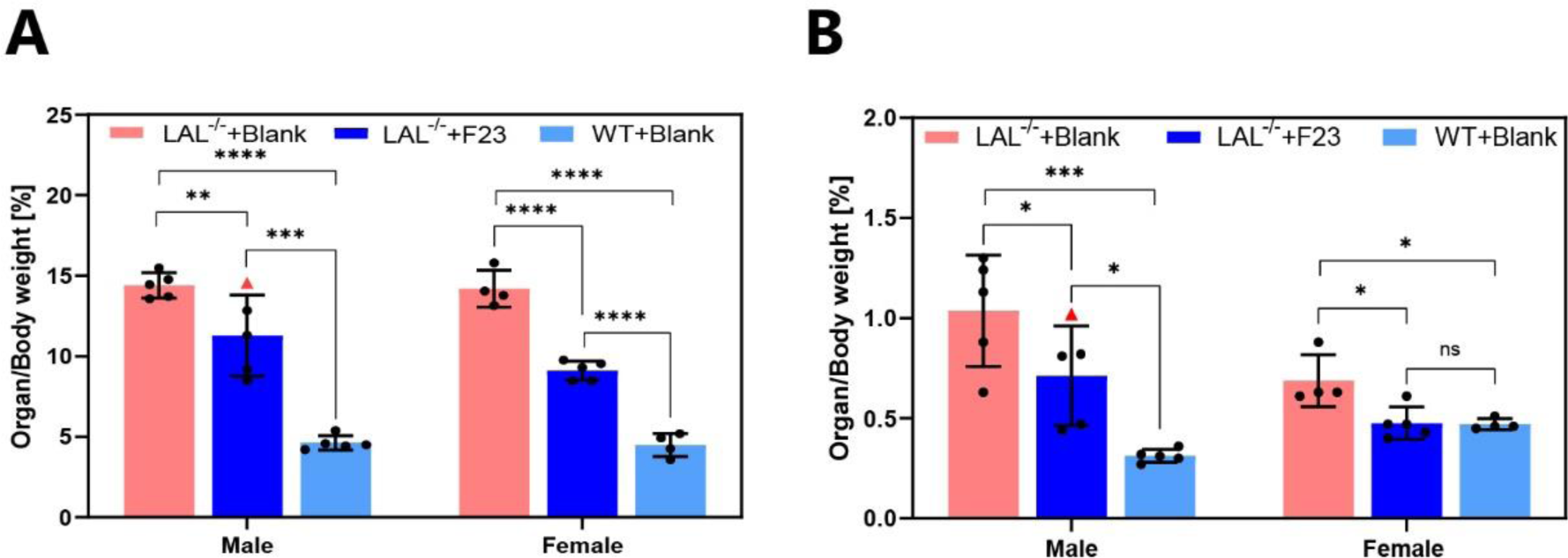
Organ weights in the efficacy study separated by sex. **(A)** Weight of the liver and **(B)** of the spleen from the three cohorts; untreated LAL^-/-^ mice (LAL^-/-^+Blank), F23-treated LAL^-/-^ mice (LAL^-/-^+F23) and wild-type control mice (WT+Blank). Red triangles depict the male mouse (0629) with a later initiation of treatment. One-way ANOVA with Tukey’s post hoc test analysis between each group in each sex *P < 0.05, **P < 0.01, ***P < 0.005, and ****P < 0.0001. Graphs show mean ± SEM (n≥4).

**Figure S16.**
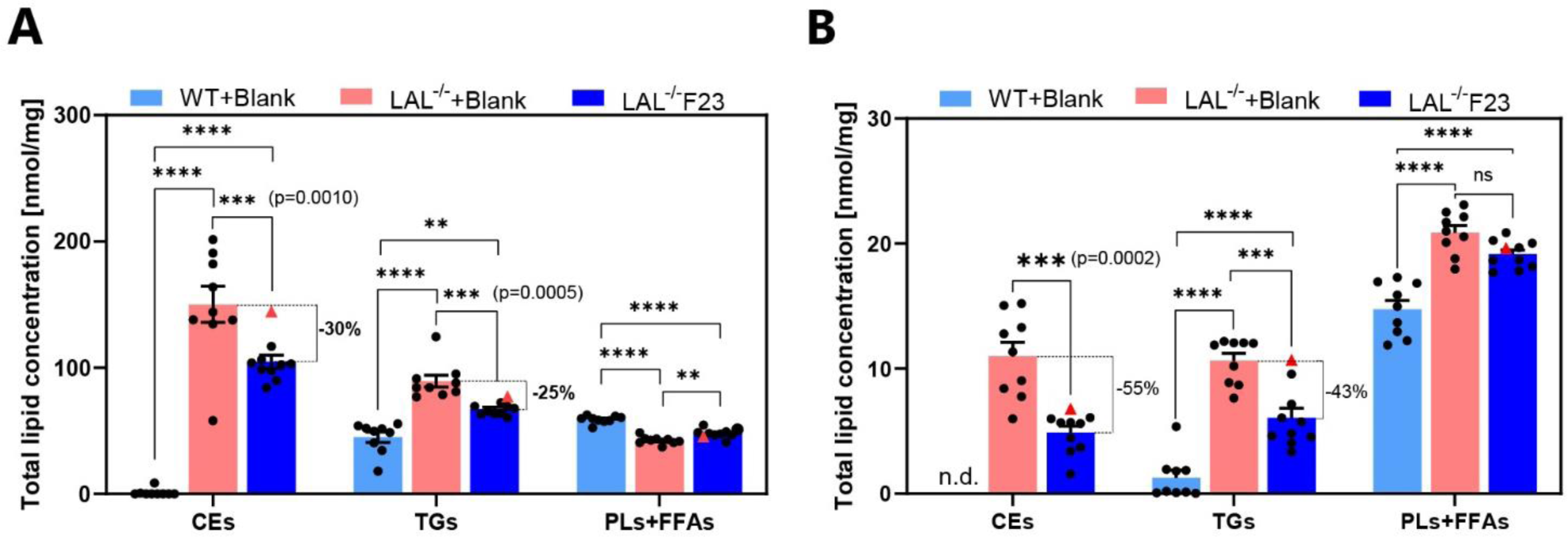
Total lipid concentrations in liver and spleen in the CE, TG, and PL + FFA fractions. **(A)** Total concentration of all 29 fatty acid methyl esters in each fraction in liver and **(B)** spleen. Red triangles depict the male mouse (0629) with a later initiation of treatment. One-way ANOVA with Tukey’s post hoc test analysis. *P < 0.05, **P < 0.01, ***P < 0.001, and ****P < 0.0001. Graphs show mean ± SEM (n≥9).

**Figure S17.**
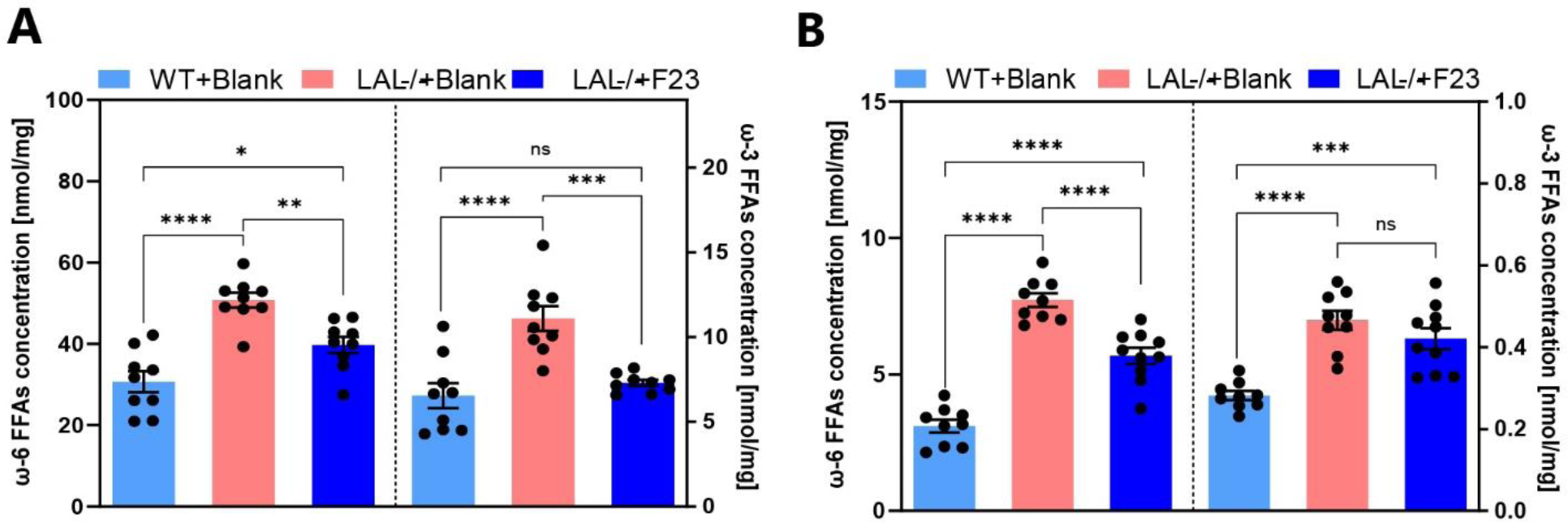
Omega-6 and omega-3 lipid concentrations. **(A)** Total omega-6 FAs in the CE, TG, and PL+FFA fractions in liver and **(B)** spleen. Omega-6 FAs include linoleic acid (C18:2n6), γ-linolenic acid (C18:3n6), eicosadienoic acid (C20:2n6), dihomo γ-linolenic acid (C20:3n6), and arachidonic acid (C20:4n6). Omega-3 fatty acids include α-linolenic acid (C18:3n3), stearidonic acid (C18:4n3), eicosatrienoic acid (C20:3n3), eicosatetraenoic acid (C20:4n3), eicosapentaenoic acid (C20:5n3), docosapentaenoic acid (C22:5n3), and docosahexaenoic acid (C22:6n3). One-way ANOVA with Tukey’s post hoc test analysis. *P < 0.05, **P < 0.01, ***P < 0.001, and ****P < 0.0001. Graphs show mean ± SEM (≥9)

**Figure S18.**
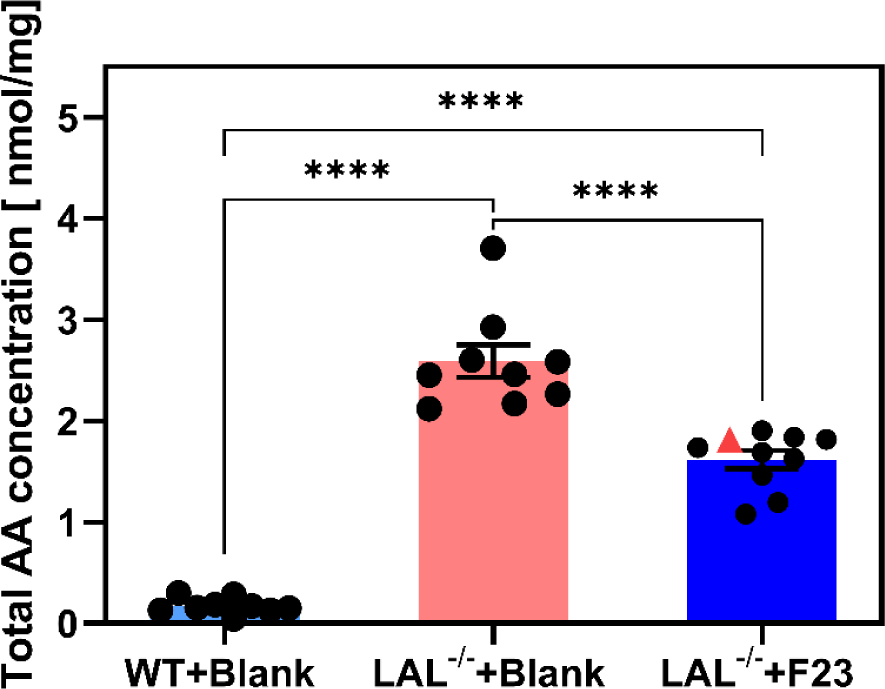
Total arachidonic acid (AA) concentrations. Total AA concentrations were pooled from the CE, TG, and PL+FFA fractions for each mouse in the cohorts. Red triangles depict the male mouse (0629) with later initiation of F23 treatment. One-way ANOVA post Tukey’s post hoc test comparison. ****P < 0.0001. Graphs show mean ± SEM (n≥9).

**Figure S19.**
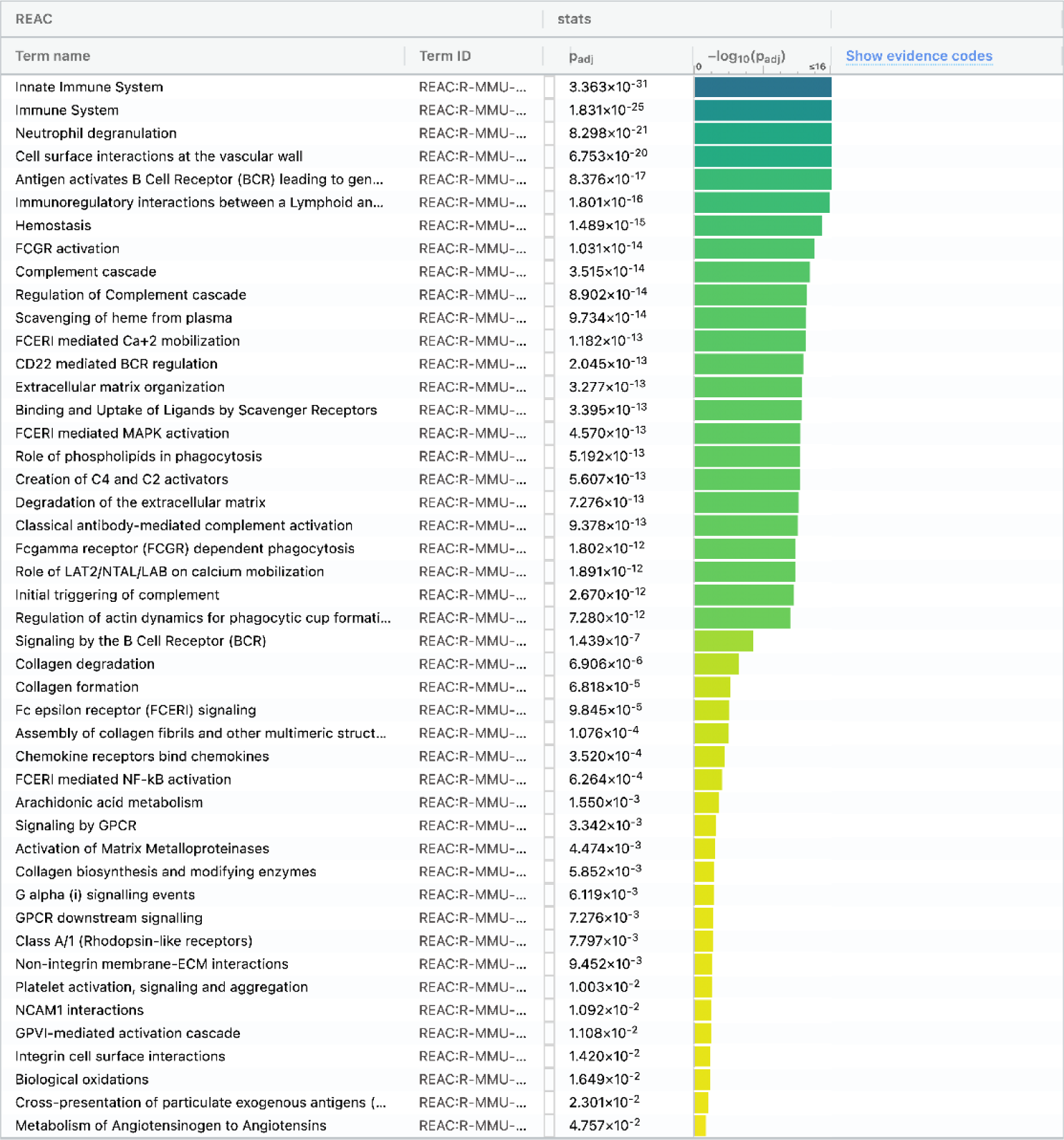
Full g:Profiler functional gene analysis for Reactome pathways. Differentially expressed genes were selected from the LAL^-/-^+Blank vs WT+Blank groups with a fold change of ≥2.0 and an FDR cutoff of ≤0.01, yielding a total of 2440 up-regulated genes and 823 down-regulated genes. The 3263 genes were computed in the web server g:Profiler.

**Figure S20.**
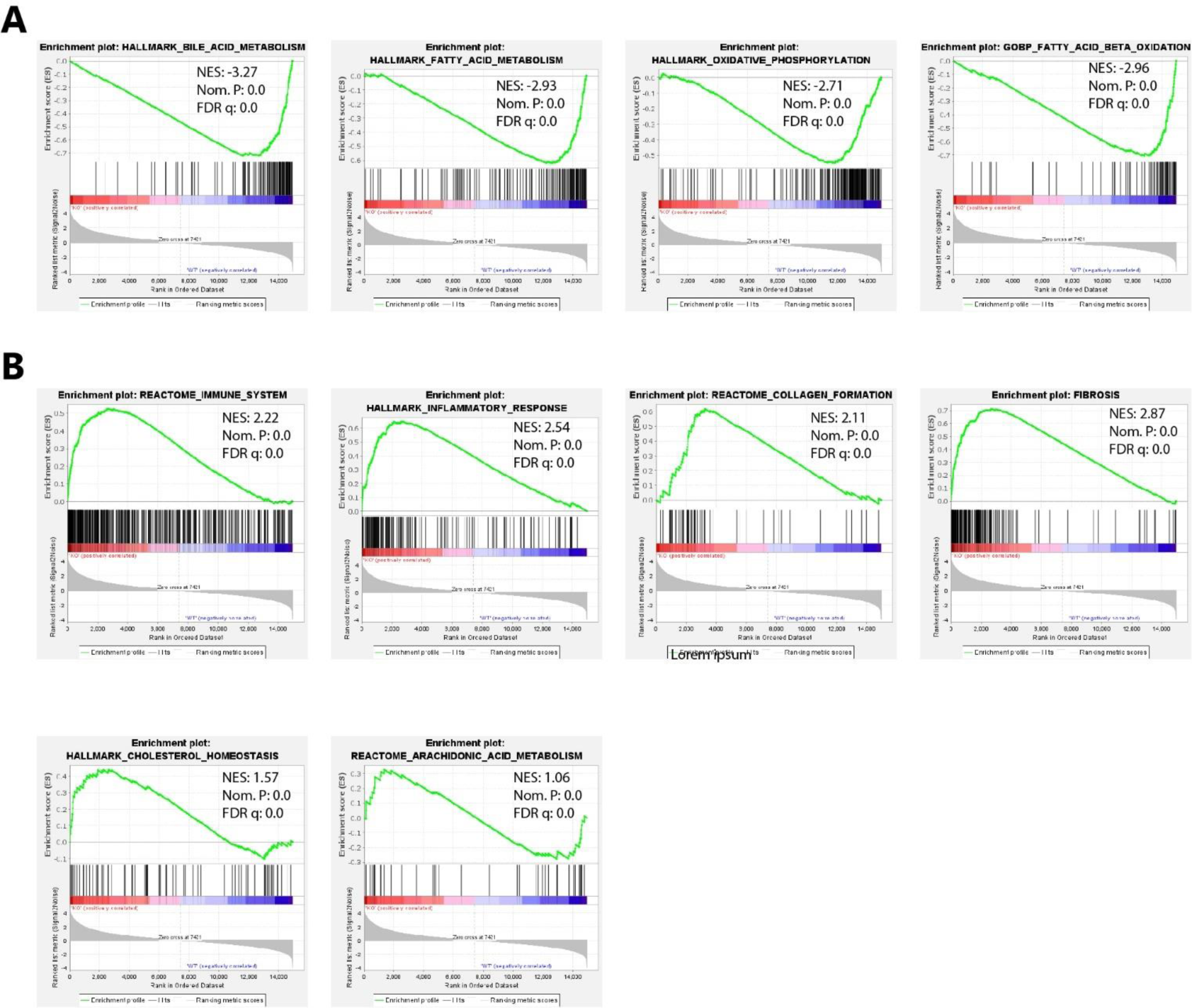
Enrichment plots from Gene Set Enrichment Analysis (GSEA) for the selected functional pathways. **(A)** down-regulated pathways (enriched in the WT+Blank group). **(B)** Up-regulated pathways (enriched in the LAL-/-+Blank group).

**Figure S21.**
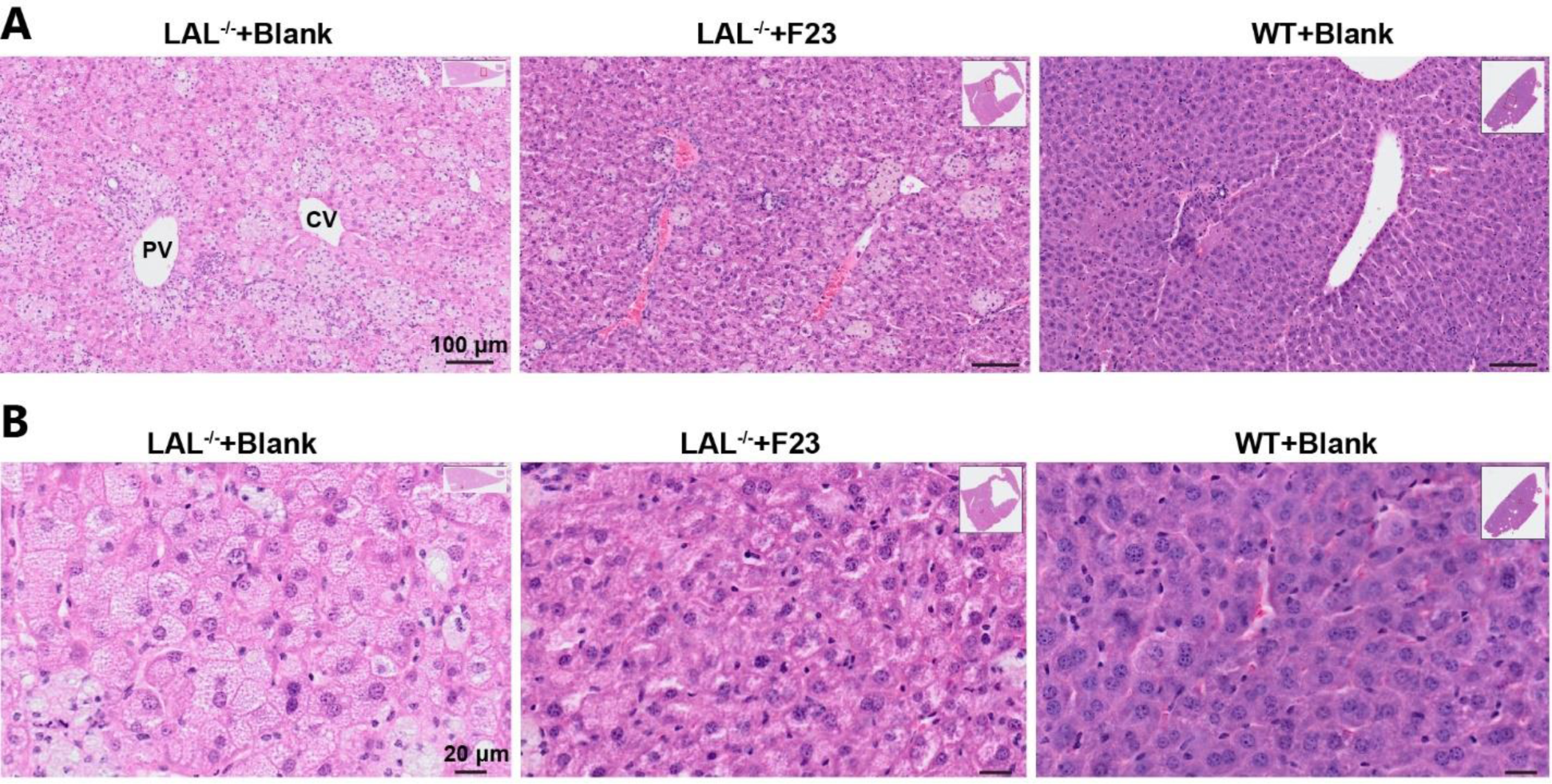
H&E staining of liver tissue. **(A)** Representative image with lower power view depicting the portal vein (PV) and centrilobular vein (CV) locations in the liver showing the presence of necrotic hepatocytes, microvesicular steatosis, and immune infiltrates, which are reduced in F23-treated LAL^−/−^ mice and not present in WT mice. Scale bar, 100 μm. **(B)** Magnification to better visualize the microvesicular steatosis spots in hepatic tissues and ballooning in hepatocytes, which is reduced by F23 treatment. Scale bar, 20 μm.

**Table S1:**
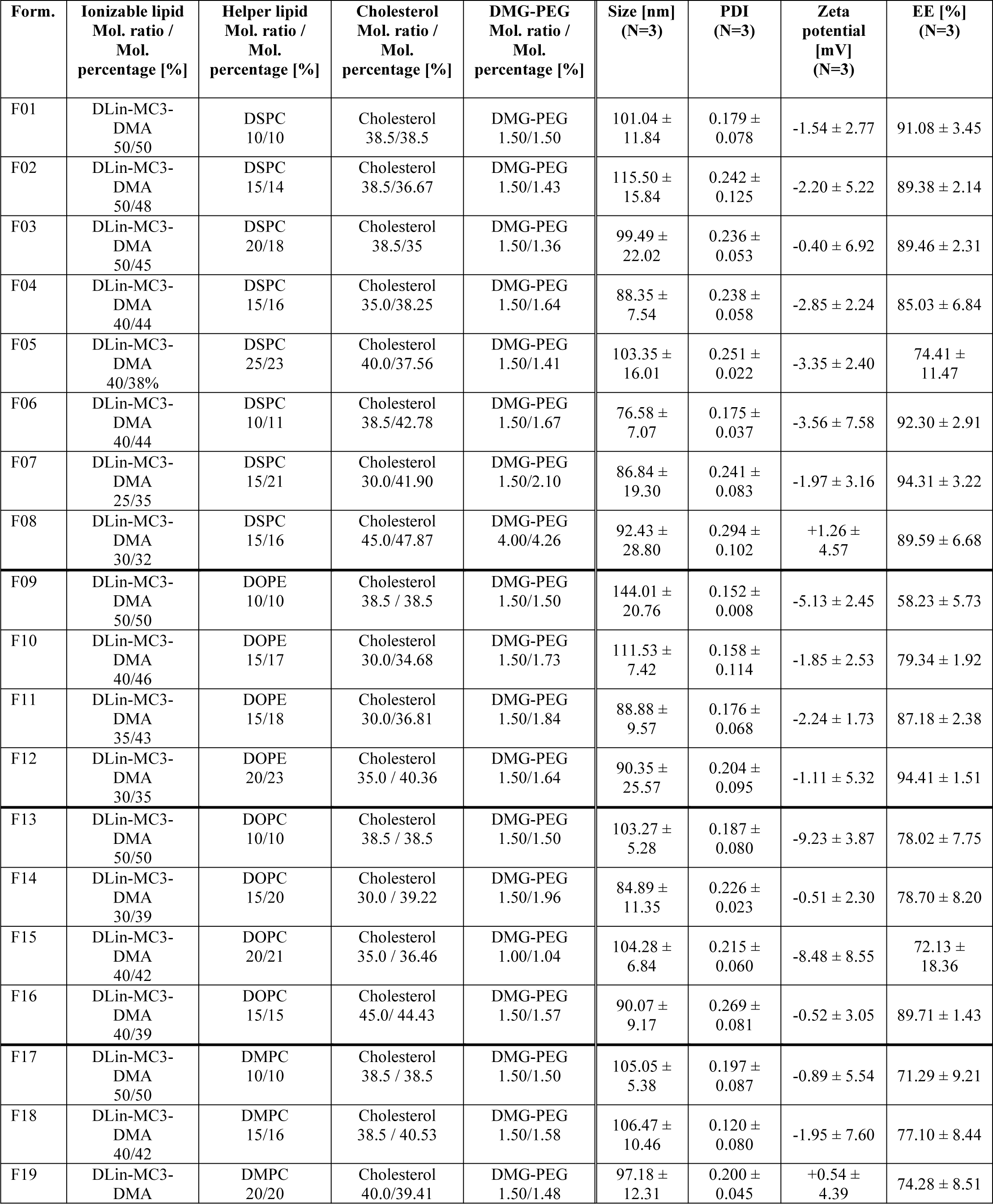
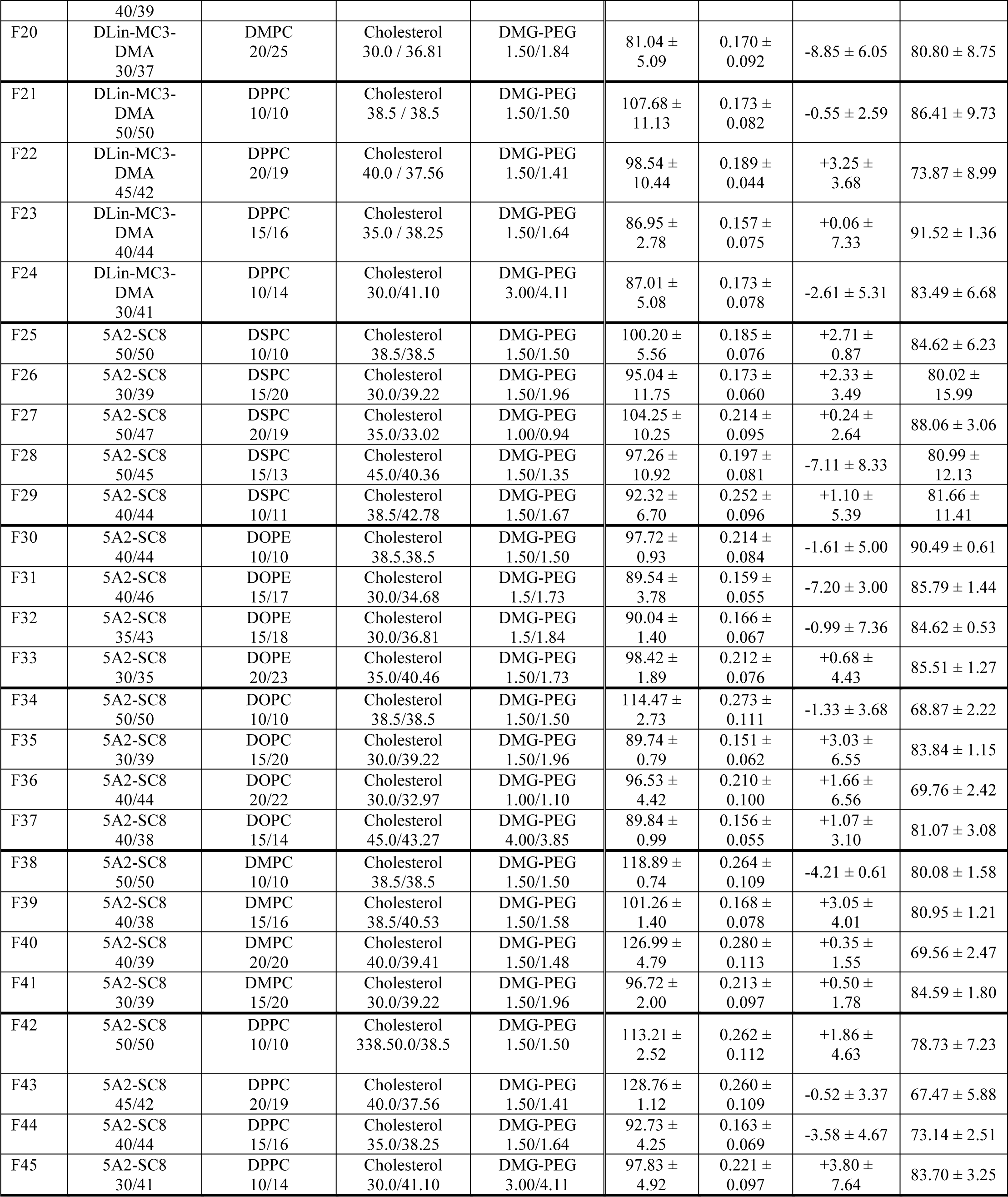
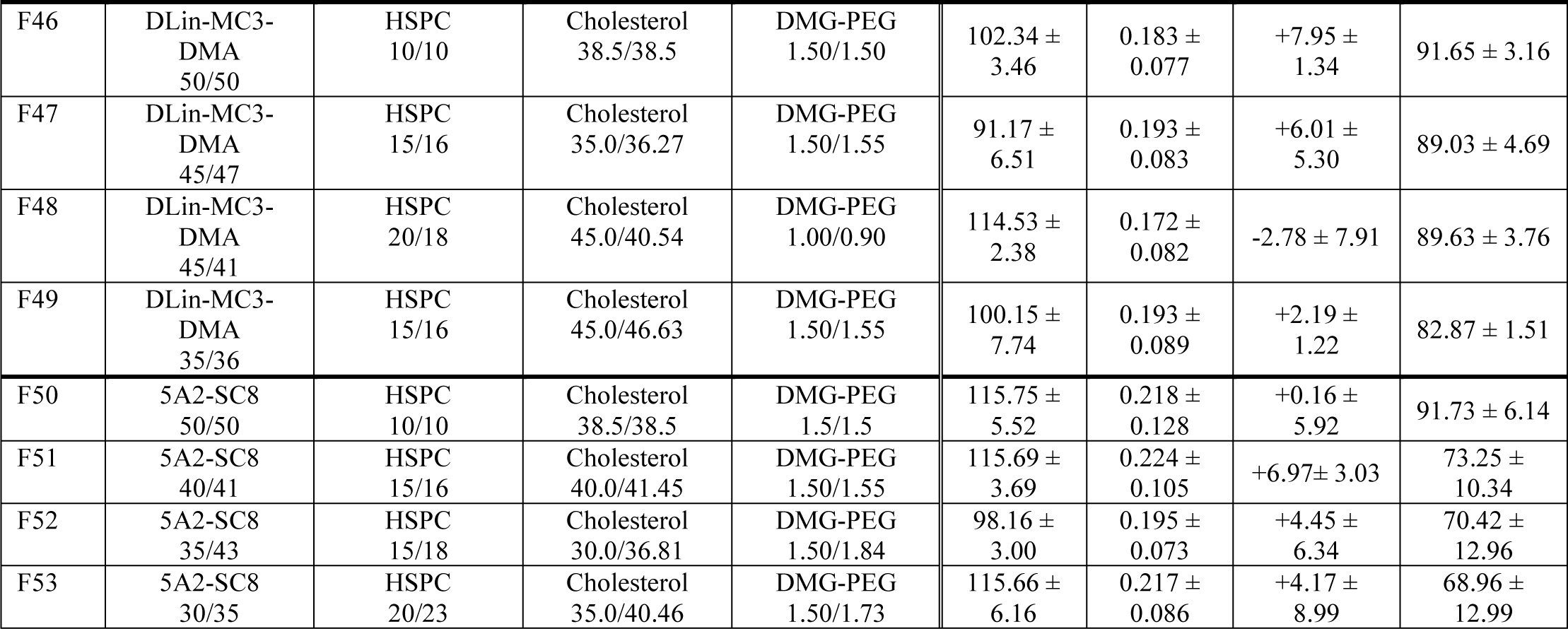
Detailed description of the 53 formulations of the combinatorial h*LIPA* mRNA-LNPs library. The composition of each lipid is given in molar ratio and molar percentage. The values of Size, PDI, encapsulation efficiency, and zeta potential are the average and standard deviations from 3 independent repeated production of each formulation.

**Table S2:**
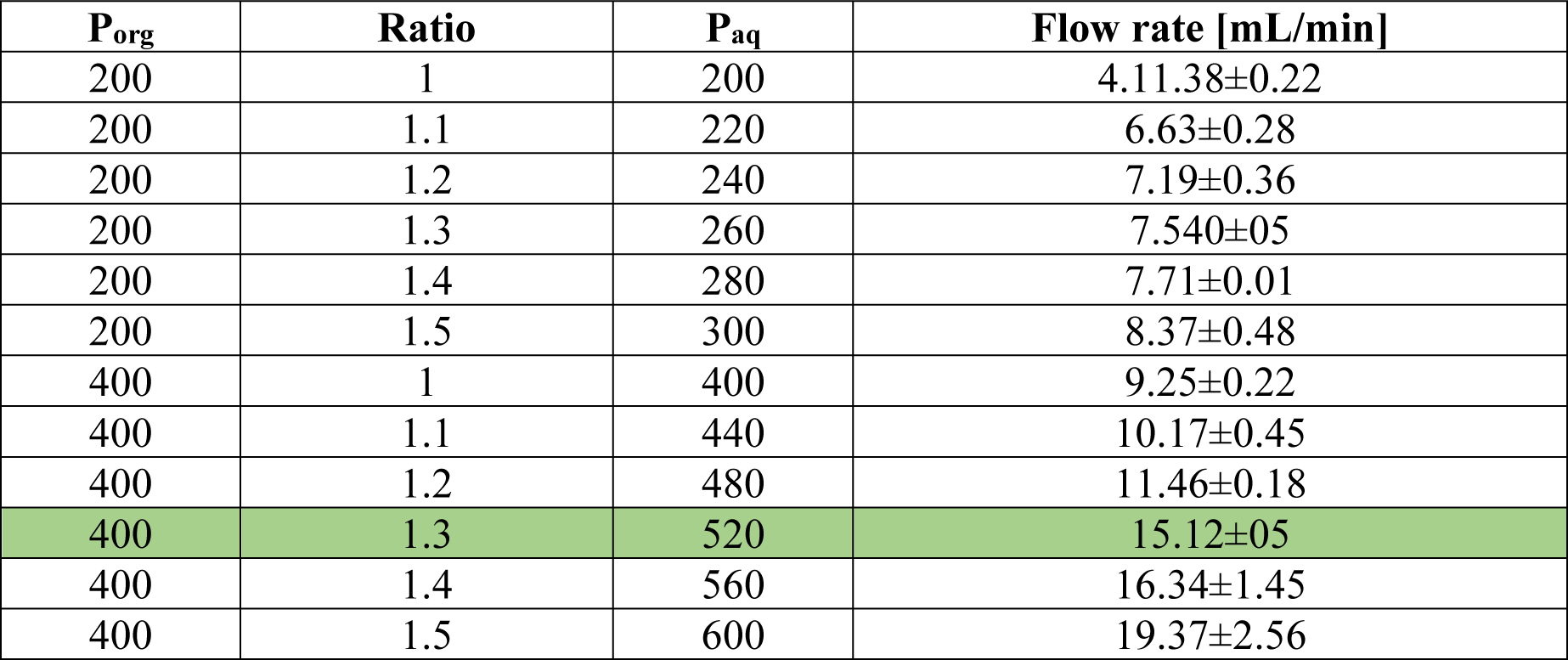
Microfluidic set up. Gradual increase of the pressure and yielded flow rate used for the optimization of the microfluidic mixing. The highlighted cells represent the chosen setup for the microfluidic mixing. Data show mean ± SD, n=3.

**Table S3:**
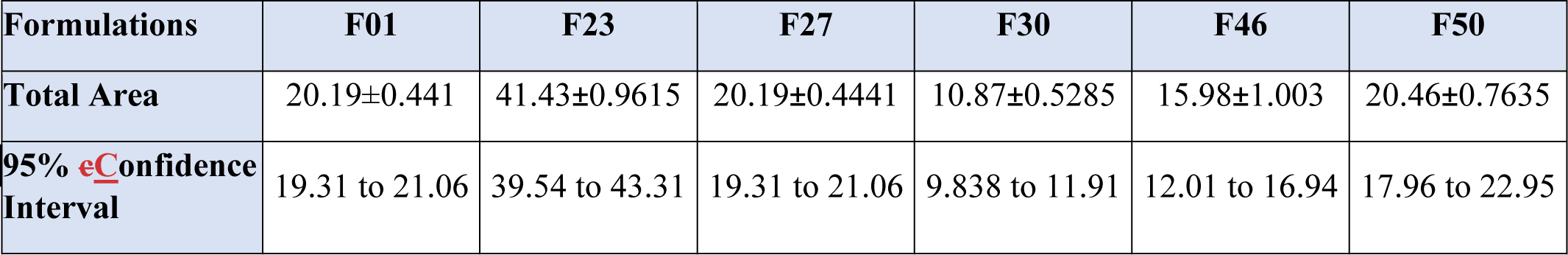
Area under the curve of mRNA-induced LAL activity of the formulations assessed in the *in vitro* kinetic analysis.

**Table S4:**
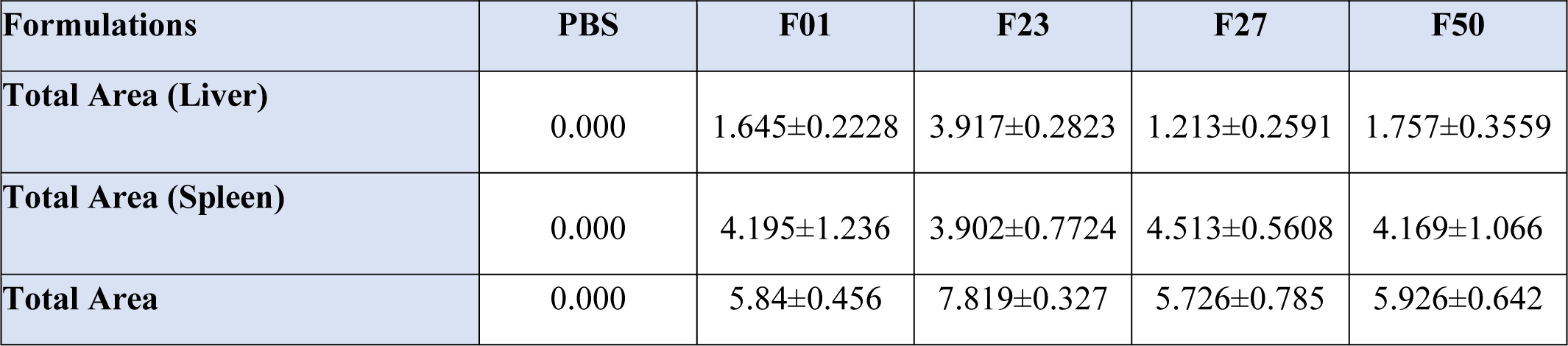
Area under the curve of mRNA-induced LAL activity in healthy mice of selected mRNA-LNP formulations. The values represent the MEAN ± SD of 3 AUC profiles per formulation.

**Table S5:**
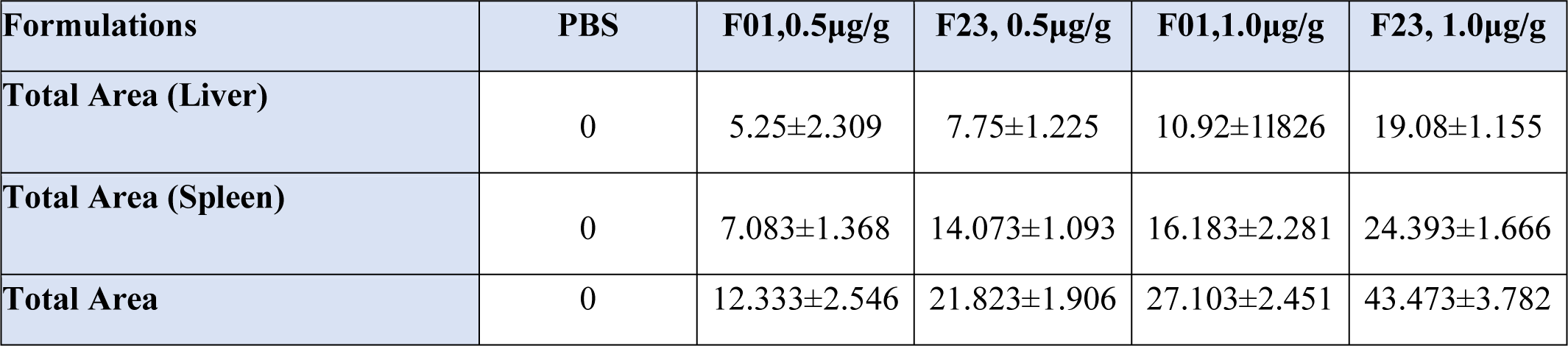
Area under the curve of mRNA-induced LAL activity in LAL^-/-^ mice. The values represent the MEAN ± SD of 3 AUC profiles per formulation.

**Table S6:**
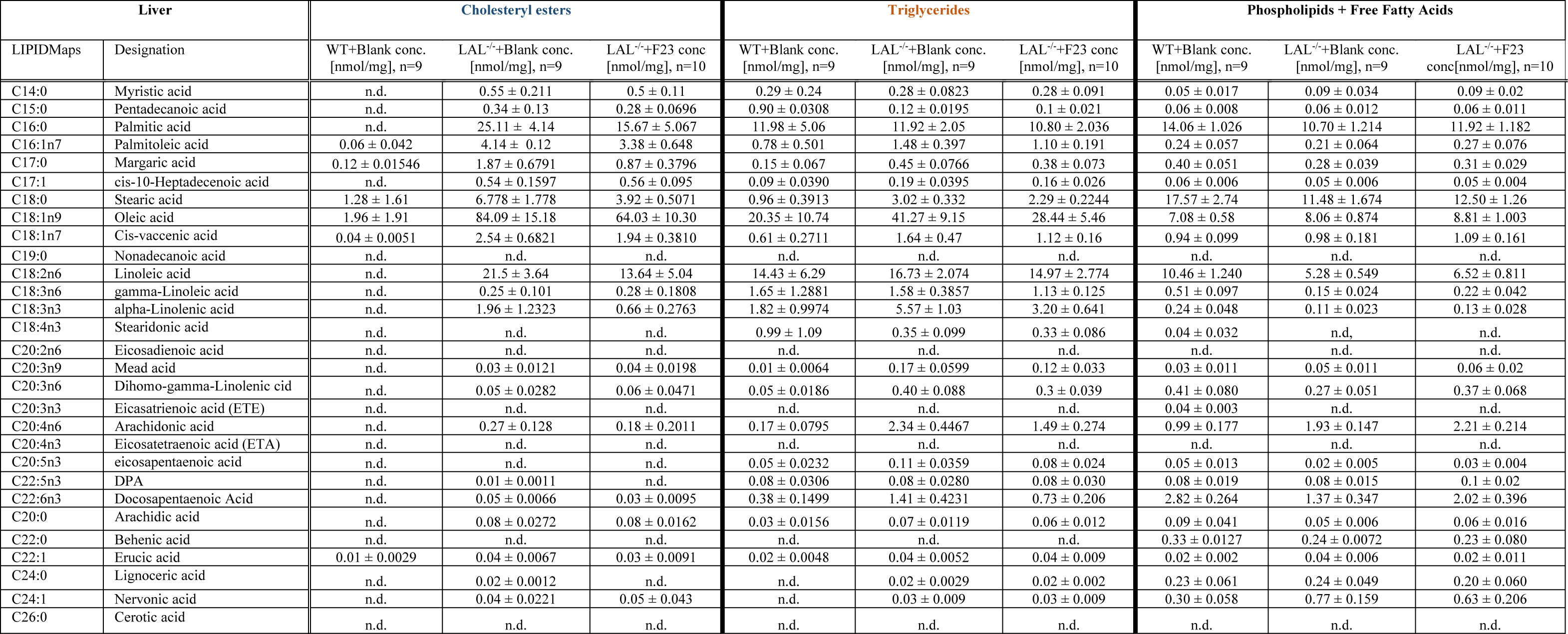
Targeted lipidomic analysis of hepatic samples from the efficacy study. Cholesteryl esters (CEs), triglycerides (TGs), and phospholipids + free fatty acids (PLs+FFAs) fractions were obtained by the separation of the liver lipid extracts of LAL^-/-^+Blank (n=9), LAL^-/-^+F23 (n=10), and WT+Blank (n=9) mice. The values depict the average ± SD for each species in the associated cohort. N.d.: not detected.

**Table S7:**
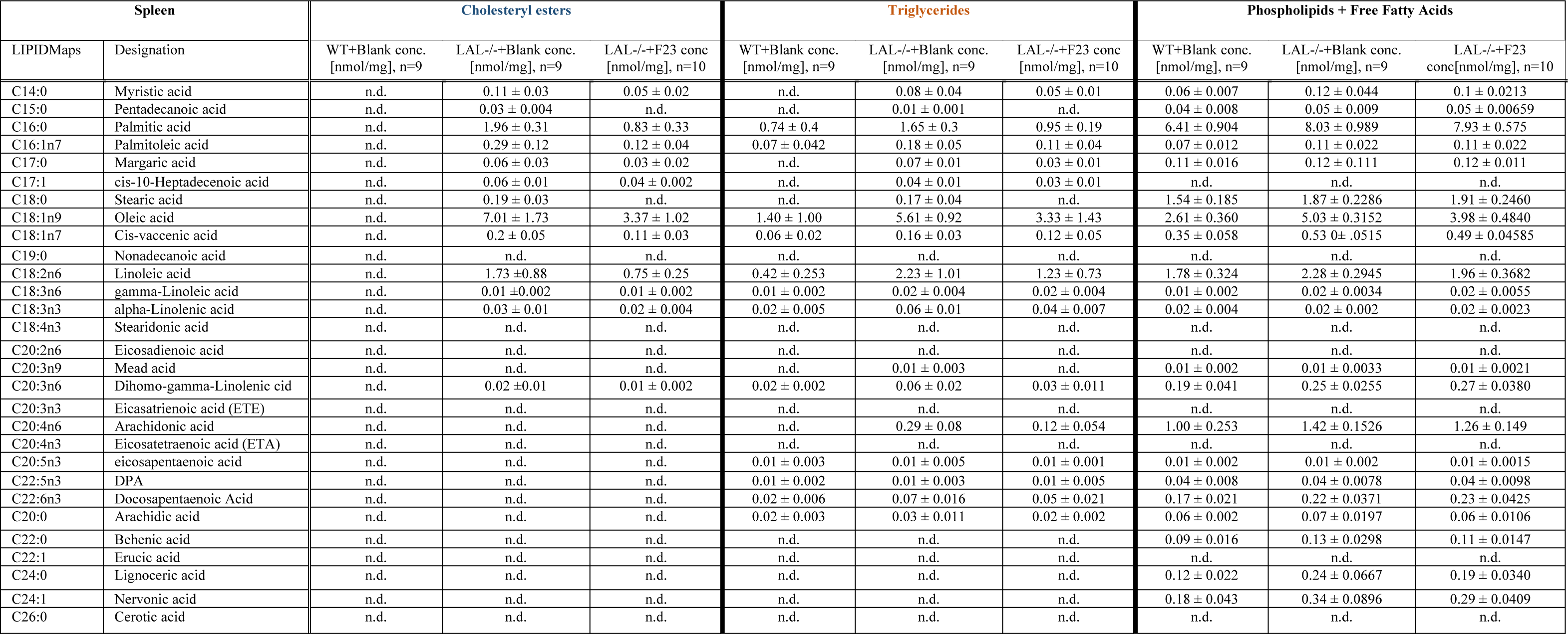
Targeted lipidomic analysis of splenic samples from the efficacy study. Cholesteryl esters (CEs), triglycerides (TGs), and phospholipids + free fatty acids (PLs+FFAs) fractions were obtained by the separation of the spleen lipid extracts of LAL^-/-^+Blank (n=9), LAL^-/-^+F23 (n=10), and WT+Blank (n=9) mice. The values depict the average ± SD for each species in the associated cohort. N.d.: not detected.

