## Supplementary Informations for "Lipid-based nanoparticles deliver mRNA to reverse the pathogenesis of lysosomal acid lipase deficiency in a preclinical model"

**Matthias Zadory et al.**

List of Supplementary Materials
Materials and Methods
Fig S1 to S21
Tables S1 to S7
References (20,64,65)

### Materials and Methods

**Cell culture**
HepG2 cells (ATCC; HB 8065, passages 10–25) were maintained in high-glucose Dulbecco's modified Eagle's medium (DMEM plus sodium pyruvate, supplemented with 10% fetal bovine serum (FBS) and 1% penicillin/streptomycin. Cells were harvested using trypsin (all reagents from Wisent) at 70-90% and reseeded to 50% confluency.

| |
|---|
| ATGAAAATGCGGTTCTTGGGGTTGGTGGTCTGTTTGGTTCTCTGGACCCTGCATTCTGAGG<br>GGTCTGGAGGGAAACTGACAGCTGTGGATCCTGAAACAAACATGAATGTGAGTGAAATTA<br>TCTCTTACTGGGGATTCCCTAGTGAGGAATACCTAGTTGAGACAGAAGATGGATATATTCT<br>GTGCCTTAACCGAATTCCTCATGGGAGGAAGAACCATTCTGACAAAGGTCCCAAACCAGT<br>TGTCTTCCTGCAACATGGCTTGCTGGCAGATTCTAGTAACTGGGTCACAAACCTTGCCAAC<br>AGCAGCCTGGGCTTCATTCTTGCTGATGCTGGTTTTGAGTGTGGATGGGCAACAGCAGAG<br>GAAATACCTGGTCTCGGAAACATAAGACACTCTCAGTTTCTCAGGATGAATTCTGGGCTTT<br>CAGTTATGATGAGATGGCAAAATATGACCTACCAGCTTCCATTAACTTCATTCTGAATAAA<br>ACTGGCCAAGAACAAGTGTATTATGTGGGTCATTCTCAAGGCACCACTATAGGTTTTATAG<br>CATTTTCACAGATCCCTGAGCTGGCTAAAAGGATTAAAATGTTTTTTGCCCTGGGTCCTGT<br>GGCTTCCGTCGCCTTCTGTCTAGCCCTATGGCCAAATTAGGACGATTACCAGATCATCTCA<br>TTAAGGACTTATTTGAGACAAAGAATTTCTTCCCCAGAGTGCGTTTTTGAAGTGGCTGGGT<br>ACCCACGTTTGACTCATGTCATACTGAAGGAGCTCTGTGGAAATCTCTGTTTTCTTCTGTGT<br>GGATTTAATGAGAGAAATTTAAATATGTCTAGAGTGGATGTATATACAACACATTCTCCTG<br>CTGGAACTTCTGTGCAAAACATGTTACACTGGAGCCAGGCTGTTAAATTCCAAAAGTTTCA<br>AGCCTTTGACTGGGGAAGCAGTGCCAAGAATTATTTTCATTACAACCAGAGTTATCTCCCA<br>CATACAATGTGAAGGACATGCTTGTGCCGACTGCAGTCTGGAGCGGGGGTCCGACTGGCT<br>TGCAGATGTCTACGACGTCAATATCTTACTGACTCAGATCACCAACTTGTGTTCCATGAGA<br>GCATTCCGGAATGGGAGCATCTTGACTTCATTTGGGGCCTGGATCCCCTTGGAGGCTTTAT<br>AATAAAATTATTAATCTAATGAGGAAATATCAGGACTACAAAGACGATGACGACAAGTGA |

| | | | |
|---|---|---|---|
| **START codon** | **Signal peptide** | **FLAG Tag** | **STOP codon** |

**Figure S1. Designed exogenous h*LIPA* mRNA.** LIPA lipase A, lysosomal acid type Homo sapiens (human) NM001127605.3 from the NCBI nucleotide database. A C-terminal FLAG TAG was added to distinguish between endogenous *LIPA* mRNA and encapsulated exogenous h*LIPA* mRNA.

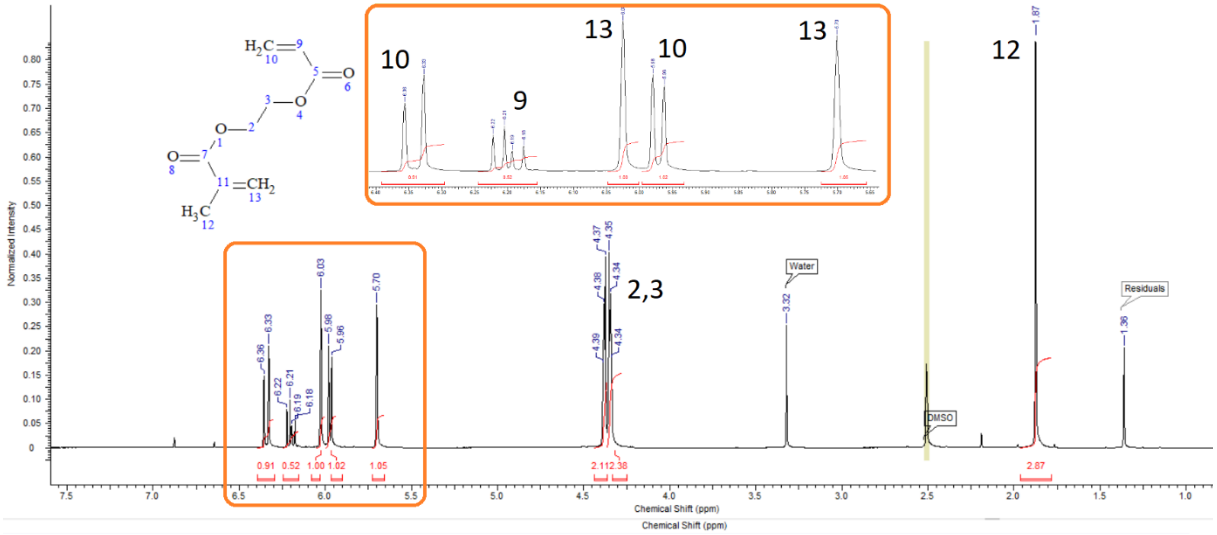


**Figure S2.** $^{1}$H-NMR spectrum of AEMA (600 MHz, D6-DMSO, δ).

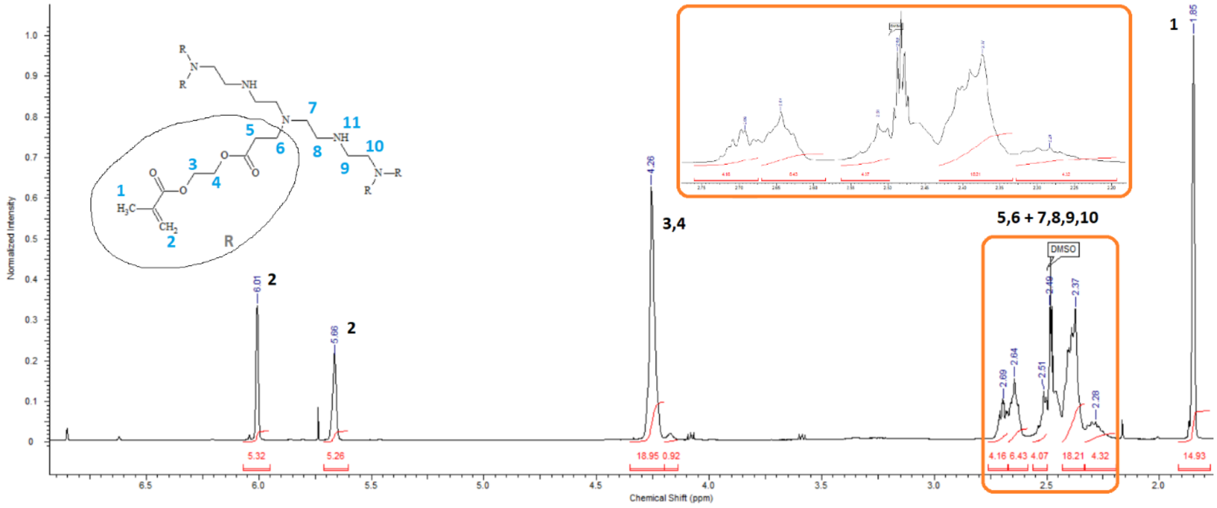


**Figure S3.** $^{1}$H-NMR spectrum of the non-purified intermediate "5A2-meth" (600 MHz, D6-DMSO, δ).

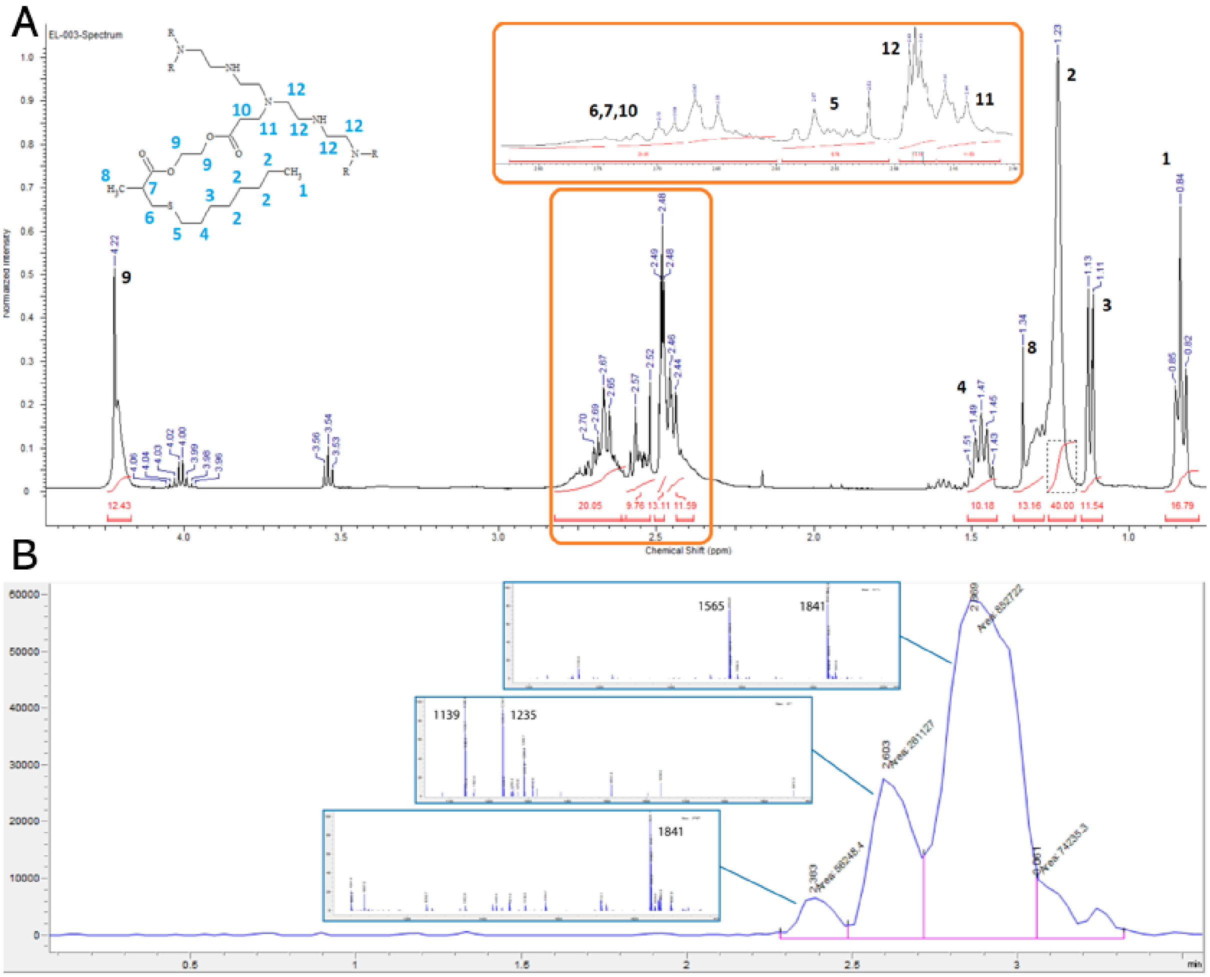


**Figure S4. Characterization of the synthesized ionizable lipid 5A2-SC8. (A)** [1]H-NMR spectrum of the synthesized lipid 5A2-SC8 (600 MHz, D6-DMSO, δ) after purification. 4.22 (s, 12H, -OC*H2*C*H2*O-), 2.70-2.65 (m, 20H, -C*H*C*H2*SC8H17; -NCH2C*H2*COO-), 2.60-2.52 (m, 10H, -SC*H2*C7H15), 2.49-2.47 (m, 14H, -NC*H2*C*H2*NC*H2*C*H2*N-), 2.46-2.44 (m, 12H, -NC*H2*CH2COO-), 1.51-1.43 (c, 10H, -SCH2C*H2*C6H13), 1.34-1.25 (m, 14H, -OCOCH(C*H3*)CH2S-), 1.23-1.16 (s, 40H, -SCH2CH2CH2C*H2*C*H2*C*H2*C*H2*CH3), 1.13-1.11 (d, 12H, -SCH2CH2C*H2*C5H11), 0.85-0.82 (t, 17H, -SC6H12C*H3*). **(B)** LC spectrum of the purified fractions after column chromatography including associated MS spectrum for each peak with the specified m/z. 5A2-SC8 is expected to appear at m/z = 1841.

.

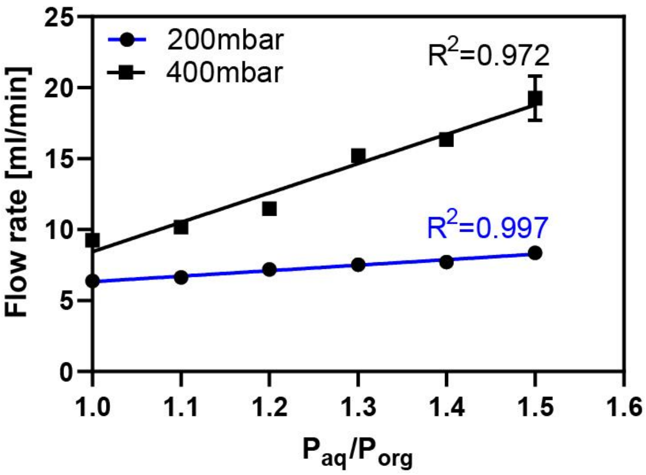


**Figure S5. Linear relation between pressure ratio and flow rate**. Increase of the ratio between the aqueous channel pressure ($P_{aq}$) versus the pressure of the organic phase ($P_{org}$) mediates an increase in the total flow rate in a linear relationship. The ratio was defined by maintaining $P_{org}$ constant and increasing $P_{aq}$.

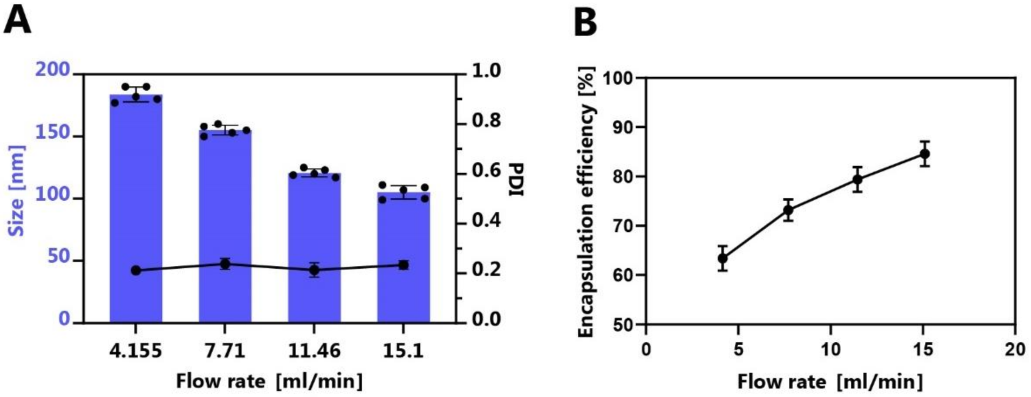


**Figure S6. Optimization of the microfluidic setup.** Dependence of the **(A)** hydrodynamic size (left axis) and the polydispersity index (right axis) as well as **(B)** the encapsulation efficiency on the flow rate of the microfluidic system. Data represent mean ± SEM of n=5 measurements.

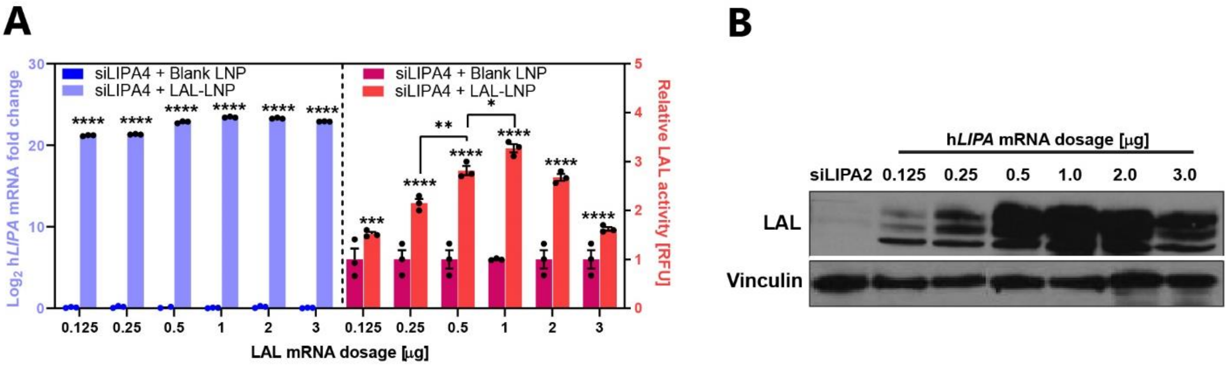


**Figure S7. *In vitro* dose-response study with F01 formulation. (A)** Exogenous h*LIPA* mRNA fold change in $Log_2$ (left y-axis) and relative h*LIPA* mRNA-induced LAL activity (right y-axis) in HepG2 cells upon transfection with increasing amounts of encapsulated h*LIPA* mRNA. Data represent mean ± SEM of n=3 independent experiments. **(B)** Immunoblotting showing the gradual increase of the band intensity with increasing h*LIPA* mRNA dosage. Vinculin was used as a loading control.

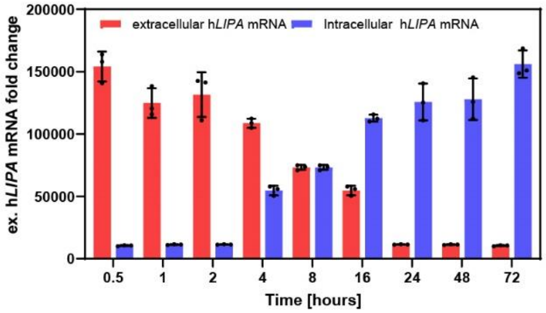


**Figure S8. *In vitro* kinetic evaluation of encapsulated h*LIPA* mRNA delivery.** F01 was transfected in HepG2 and the intracellular h*LIPA* mRNA levels were determined in the cell lysate and the medium at the indicated time points. The graph shows mean ± SEM of 3 independent experiments.

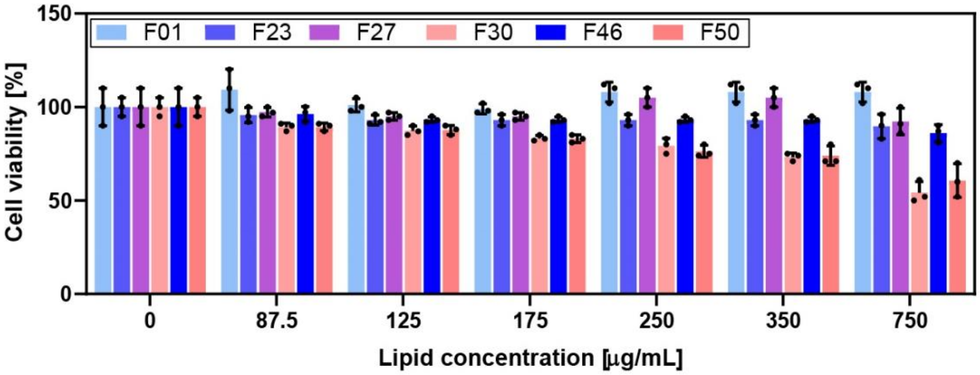


**Figure S9. Cell viability of the tested formulations.** Viability measurement 24 hours post-exposure of HepG2 cells to the mRNA-LNP formulations compared to cells grown in the absence of lipids. Data represent mean ± SEM of 3 independent experiments.

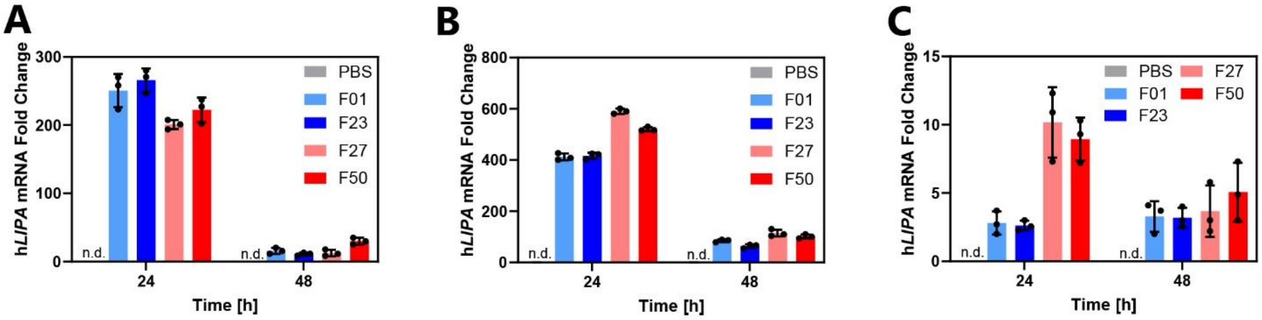


**Figure S10. h*LIPA* mRNA levels delivered to tissues by selected formulations. (A)** Exogenous h*LIPA*-mRNA delivered in the liver, **(B)** the spleen, and **(C)** in the lungs. Fold change have been normalized on murine TAT-binding protein (TBP). Data represent mean ± SEM (n = 3).

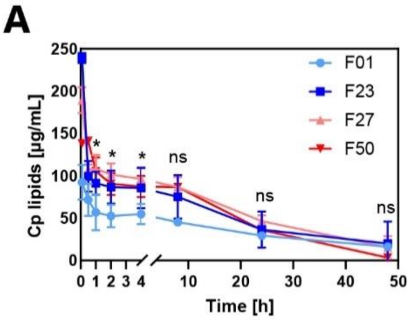


**B**

| Parameters | F01 | F23 | F27 | F50 |
|---|---|---|---|---|
| $T_{1/2}$ [h] | 5.8 ± 0.5 | 6.2 ± 0.5 | 5.4 ± 0.27 | 5.6 ± 0.35 |
| AUC [ng/ml*h] | 10329 ± 208 | 11435.5 ± 278 | 11392.8 ± 567 | 10441 ± 335 |
| Clearance [ml/h/kg] | 61 ± 6.1 | 68 ± 2.3 | 57 ± 8.2 | 56 ± 12.3 |

**Figure S11. Pharmacokinetic analysis of the selected LNPs in healthy mice following i.v. administration. (A)** Time-dependent changes of the plasma total lipid concentration (Cp) of the selected LNPs. **(B)** Pharmacokinetic parameters of the selected formulations. Data represent mean ± SEM (n = 3).

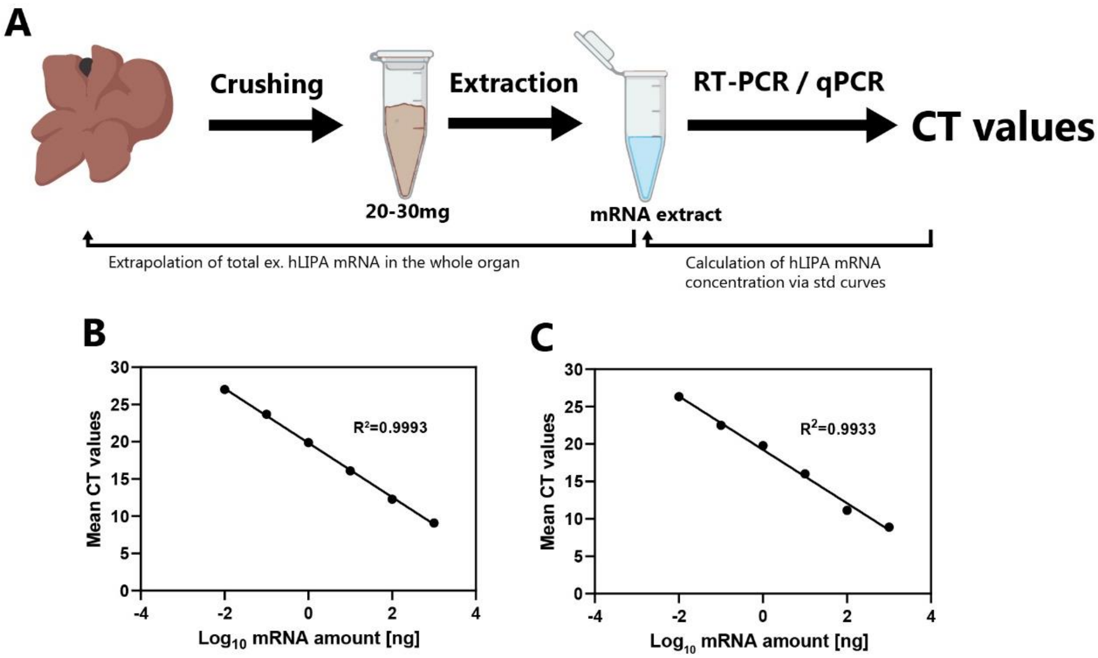


**Figure S12. Quantification of the absolute amount of exogenous h*LIPA* mRNA delivered to the liver or the spleen. (A)** Process workflow for the extrapolation of the absolute h*LIPA* mRNA amount delivered to the liver or spleen. **(B)** Standard curves with the cycle threshold (CT) values versus the h*LIPA* mRNA amount of the standards in logarithmic scale for liver and **(C)** spleen.

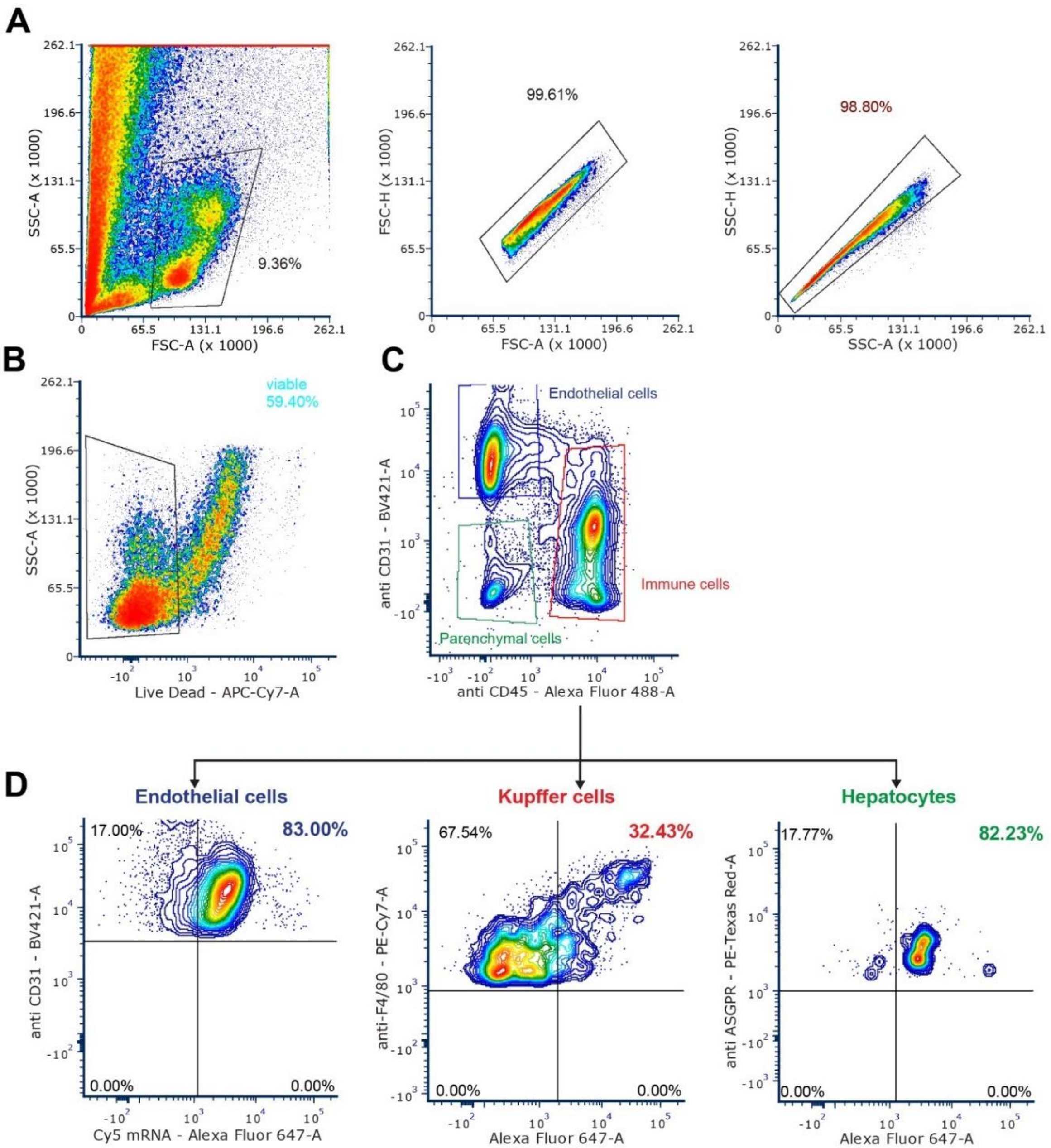


**Figure S13. Flow cytometry gating strategy for immunophenotyping and transfection efficiency of F23 in hepatic cell subtypes. (A)** All events recorded and subsequent isolation of single cells. **(B)** isolation of viable cells based on negative Live-dead staining. **(C)** preliminary separation of hepatic subpopulation defined as: immune cells CD31$^-$ and CD45$^{+,}$ endothelial cells CD31$^+$ and CD45$^-$, parenchymal cells CD31$^-$, CD45. **(D)** From **C**, Kupffer cells were specifically isolated as F4/80$^+$ from the immune cells gate and hepatocytes as ASGPR$^+$ from the parenchymal gate. In all subpopulation Cy5 mRNA signal was measured. In total n=3 mice.

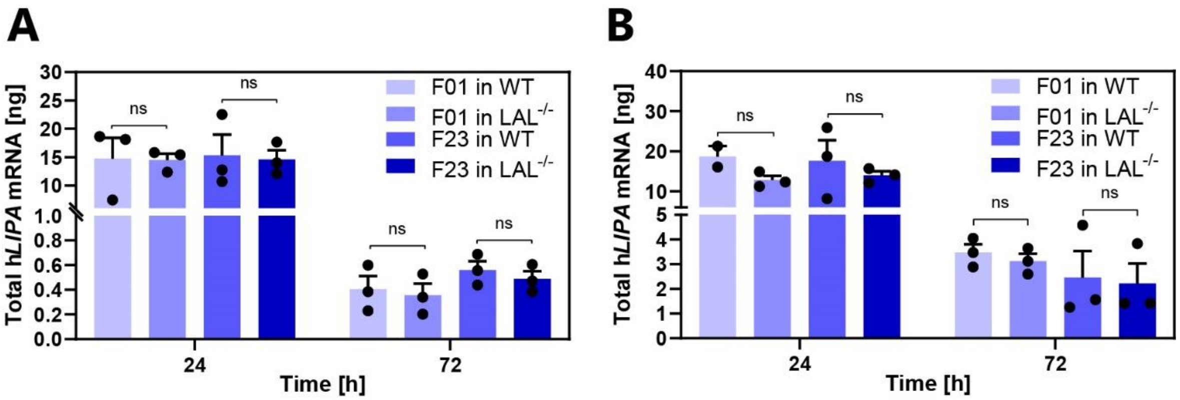


**Figure S14. Comparison of mRNA delivery in WT and LAL$^{-/-}$ mice. (A)** mRNA levels in the liver and **(B)** spleen. Unpaired two-tailed Student's *t*-test. Graphs show mean ± SEM (n=3). ns: non-significant.

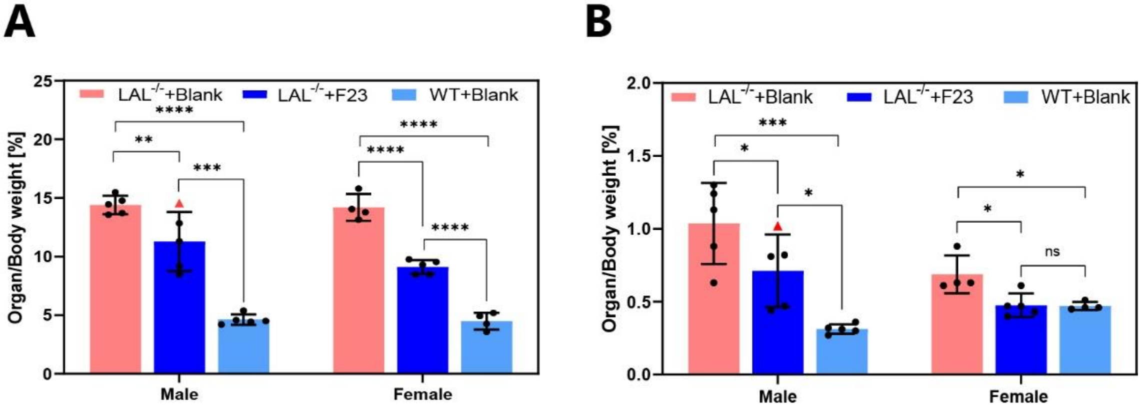


**Figure S15. Organ weights in the efficacy study separated by sex. (A)** Weight of the liver and **(B)** of the spleen from the three cohorts; untreated LAL$^{-/-}$ mice (LAL$^{-/-}$+Blank), F23-treated LAL$^{-/-}$ mice (LAL$^{-/-}$+F23) and wild-type control mice (WT+Blank). Red triangles depict the male mouse (0629) with a later initiation of treatment. One-way ANOVA with Tukey's post hoc test analysis between each group in each sex *P < 0.05, **P < 0.01, ***P < 0.005, and ****P < 0.0001. Graphs show mean ± SEM (n≥4).

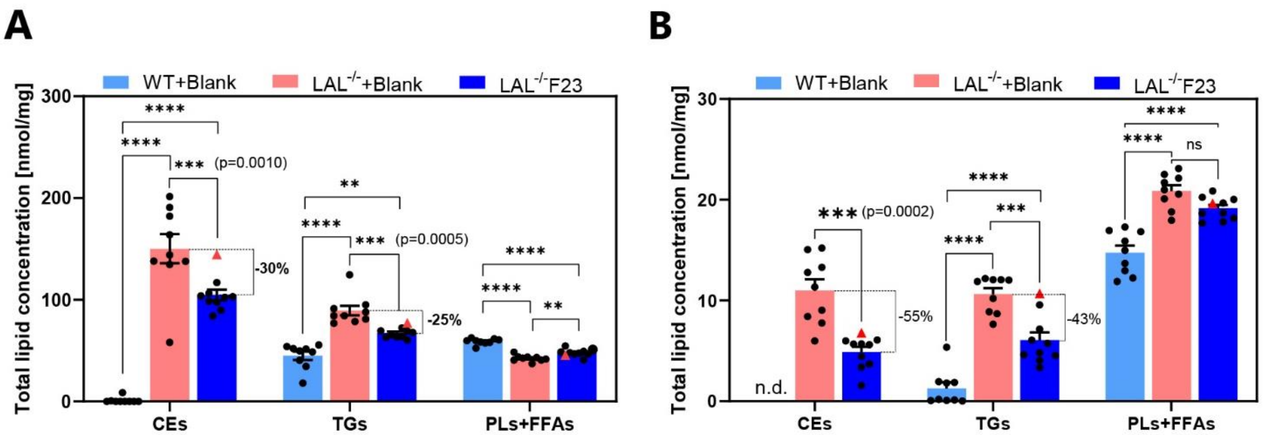


**Figure S16. Total lipid concentrations in liver and spleen in the CE, TG, and PL + FFA fractions. (A)** Total concentration of all 29 fatty acid methyl esters in each fraction in liver and **(B)** spleen. Red triangles depict the male mouse (0629) with a later initiation of treatment. One-way ANOVA with Tukey's post hoc test analysis. *P < 0.05, **P < 0.01, ***P < 0.001, and ****P < 0.0001. Graphs show mean ± SEM (n≥9).

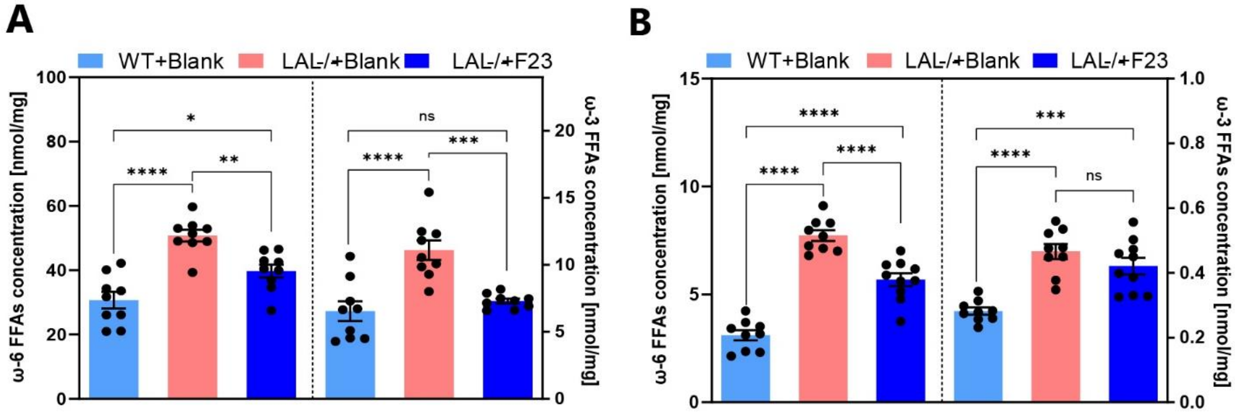


**Figure S17. Omega-6 and omega-3 lipid concentrations. (A)** Total omega-6 FAs in the CE, TG, and PL+FFA fractions in liver and **(B)** spleen. Omega-6 FAs include linoleic acid (C18:2n6), γ-linolenic acid (C18:3n6), eicosadienoic acid (C20:2n6), dihomo γ-linolenic acid (C20:3n6), and arachidonic acid (C20:4n6). Omega-3 fatty acids include α-linolenic acid (C18:3n3), stearidonic acid (C18:4n3), eicosatrienoic acid (C20:3n3), eicosatetraenoic acid (C20:4n3), eicosapentaenoic acid (C20:5n3), docosapentaenoic acid (C22:5n3), and docosahexaenoic acid (C22:6n3). One-way ANOVA with Tukey's post hoc test analysis. *P < 0.05, **P < 0.01, ***P < 0.001, and ****P < 0.0001. Graphs show mean ± SEM (≥9)

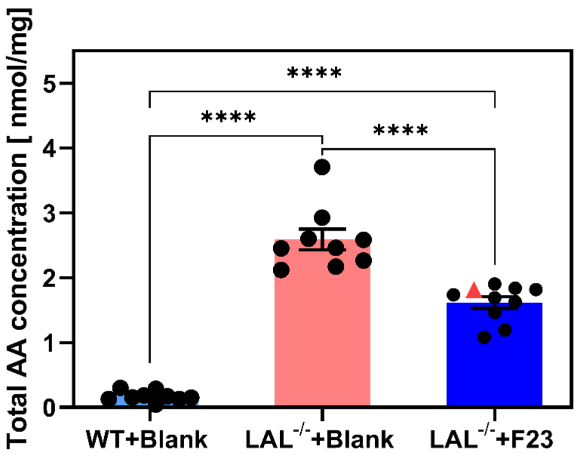


**Figure S18. Total** arachidonic acid (**AA) concentrations.** Total AA concentrations were pooled from the CE, TG, and PL+FFA fractions for each mouse in the cohorts. Red triangles depict the male mouse (0629) with later initiation of F23 treatment. One-way ANOVA post Tukey's post hoc test comparison. ****P < 0.0001. Graphs show mean ± SEM (n≥9).

| REAC | | stats | | |
|---|---|---|---|---|
| Term name | Term ID | $p_{adj}$ | $-\log_{10}(p_{adj})$ 0 … ≤16 | Show evidence codes |
| Innate Immune System | REAC:R-MMU-... | $3.363\times10^{-31}$ | | |
| Immune System | REAC:R-MMU-... | $1.831\times10^{-25}$ | | |
| Neutrophil degranulation | REAC:R-MMU-... | $8.298\times10^{-21}$ | | |
| Cell surface interactions at the vascular wall | REAC:R-MMU-... | $6.753\times10^{-20}$ | | |
| Antigen activates B Cell Receptor (BCR) leading to gen... | REAC:R-MMU-... | $8.376\times10^{-17}$ | | |
| Immunoregulatory interactions between a Lymphoid an... | REAC:R-MMU-... | $1.801\times10^{-16}$ | | |
| Hemostasis | REAC:R-MMU-... | $1.489\times10^{-15}$ | | |
| FCGR activation | REAC:R-MMU-... | $1.031\times10^{-14}$ | | |
| Complement cascade | REAC:R-MMU-... | $3.515\times10^{-14}$ | | |
| Regulation of Complement cascade | REAC:R-MMU-... | $8.902\times10^{-14}$ | | |
| Scavenging of heme from plasma | REAC:R-MMU-... | $9.734\times10^{-14}$ | | |
| FCERI mediated Ca+2 mobilization | REAC:R-MMU-... | $1.182\times10^{-13}$ | | |
| CD22 mediated BCR regulation | REAC:R-MMU-... | $2.045\times10^{-13}$ | | |
| Extracellular matrix organization | REAC:R-MMU-... | $3.277\times10^{-13}$ | | |
| Binding and Uptake of Ligands by Scavenger Receptors | REAC:R-MMU-... | $3.395\times10^{-13}$ | | |
| FCERI mediated MAPK activation | REAC:R-MMU-... | $4.570\times10^{-13}$ | | |
| Role of phospholipids in phagocytosis | REAC:R-MMU-... | $5.192\times10^{-13}$ | | |
| Creation of C4 and C2 activators | REAC:R-MMU-... | $5.607\times10^{-13}$ | | |
| Degradation of the extracellular matrix | REAC:R-MMU-... | $7.276\times10^{-13}$ | | |
| Classical antibody-mediated complement activation | REAC:R-MMU-... | $9.378\times10^{-13}$ | | |
| Fcgamma receptor (FCGR) dependent phagocytosis | REAC:R-MMU-... | $1.802\times10^{-12}$ | | |
| Role of LAT2/NTAL/LAB on calcium mobilization | REAC:R-MMU-... | $1.891\times10^{-12}$ | | |
| Initial triggering of complement | REAC:R-MMU-... | $2.670\times10^{-12}$ | | |
| Regulation of actin dynamics for phagocytic cup formati... | REAC:R-MMU-... | $7.280\times10^{-12}$ | | |
| Signaling by the B Cell Receptor (BCR) | REAC:R-MMU-... | $1.439\times10^{-7}$ | | |
| Collagen degradation | REAC:R-MMU-... | $6.906\times10^{-6}$ | | |
| Collagen formation | REAC:R-MMU-... | $6.818\times10^{-5}$ | | |
| Fc epsilon receptor (FCERI) signaling | REAC:R-MMU-... | $9.845\times10^{-5}$ | | |
| Assembly of collagen fibrils and other multimeric struct... | REAC:R-MMU-... | $1.076\times10^{-4}$ | | |
| Chemokine receptors bind chemokines | REAC:R-MMU-... | $3.520\times10^{-4}$ | | |
| FCERI mediated NF-kB activation | REAC:R-MMU-... | $6.264\times10^{-4}$ | | |
| Arachidonic acid metabolism | REAC:R-MMU-... | $1.550\times10^{-3}$ | | |
| Signaling by GPCR | REAC:R-MMU-... | $3.342\times10^{-3}$ | | |
| Activation of Matrix Metalloproteinases | REAC:R-MMU-... | $4.474\times10^{-3}$ | | |
| Collagen biosynthesis and modifying enzymes | REAC:R-MMU-... | $5.852\times10^{-3}$ | | |
| G alpha (i) signalling events | REAC:R-MMU-... | $6.119\times10^{-3}$ | | |
| GPCR downstream signalling | REAC:R-MMU-... | $7.276\times10^{-3}$ | | |
| Class A/1 (Rhodopsin-like receptors) | REAC:R-MMU-... | $7.797\times10^{-3}$ | | |
| Non-integrin membrane-ECM interactions | REAC:R-MMU-... | $9.452\times10^{-3}$ | | |
| Platelet activation, signaling and aggregation | REAC:R-MMU-... | $1.003\times10^{-2}$ | | |
| NCAM1 interactions | REAC:R-MMU-... | $1.092\times10^{-2}$ | | |
| GPVI-mediated activation cascade | REAC:R-MMU-... | $1.108\times10^{-2}$ | | |
| Integrin cell surface interactions | REAC:R-MMU-... | $1.420\times10^{-2}$ | | |
| Biological oxidations | REAC:R-MMU-... | $1.649\times10^{-2}$ | | |
| Cross-presentation of particulate exogenous antigens (... | REAC:R-MMU-... | $2.301\times10^{-2}$ | | |
| Metabolism of Angiotensinogen to Angiotensins | REAC:R-MMU-... | $4.757\times10^{-2}$ | | |

**Figure S19. Full g:Profiler functional gene analysis for Reactome pathways.** Differentially expressed genes were selected from the LAL$^{-/-}$+Blank vs WT+Blank groups with a fold change of ≥2.0 and an FDR cutoff of ≤0.01, yielding a total of 2440 up-regulated genes and 823 down-regulated genes. The 3263 genes were computed in the web server g:Profiler.

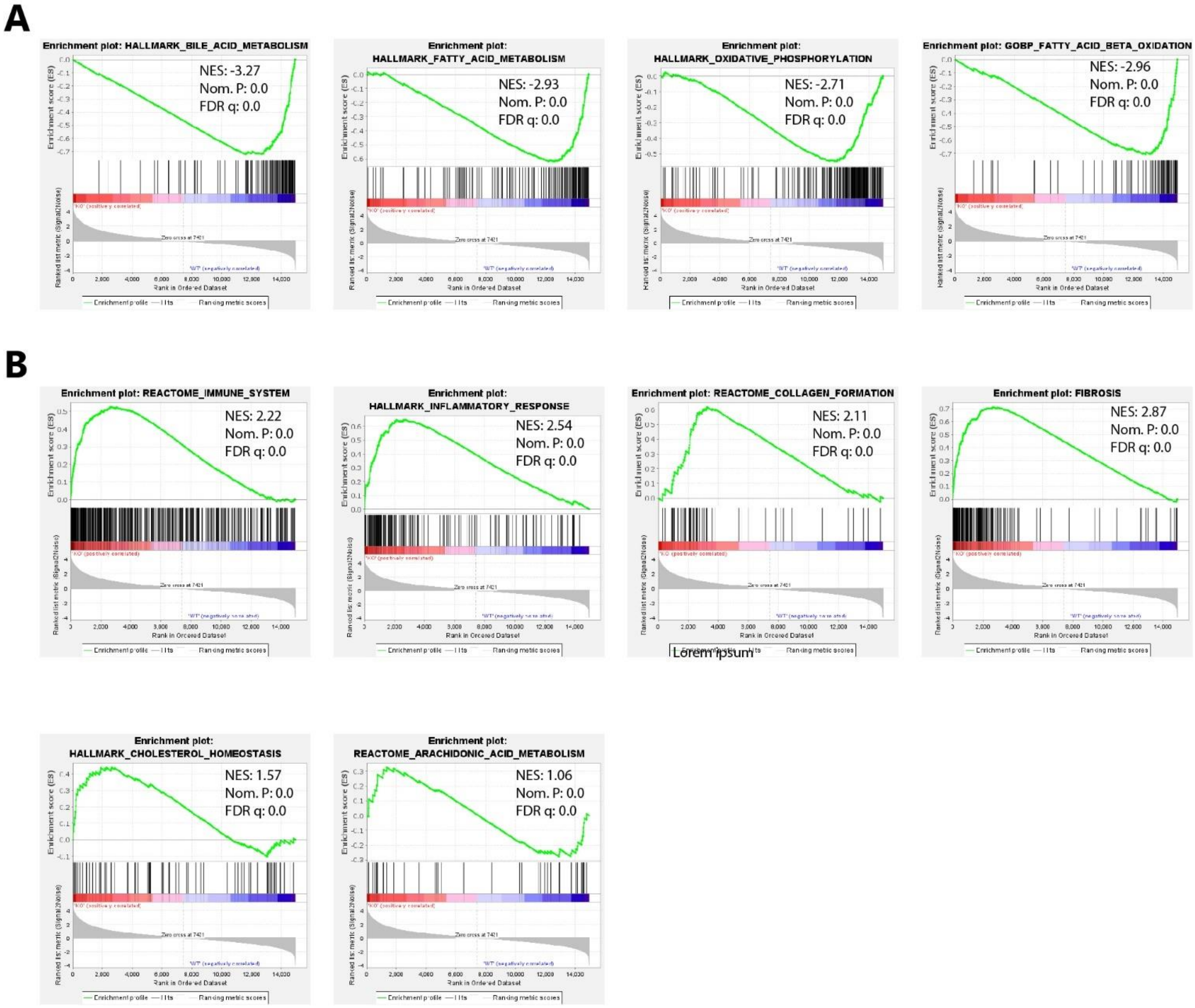


**Figure S20**. **Enrichment plots from Gene Set Enrichment Analysis (GSEA) for the selected functional pathways. (A)** down-regulated pathways (enriched in the WT+Blank group). **(B)** Up-regulated pathways (enriched in the LAL-/-+Blank group).

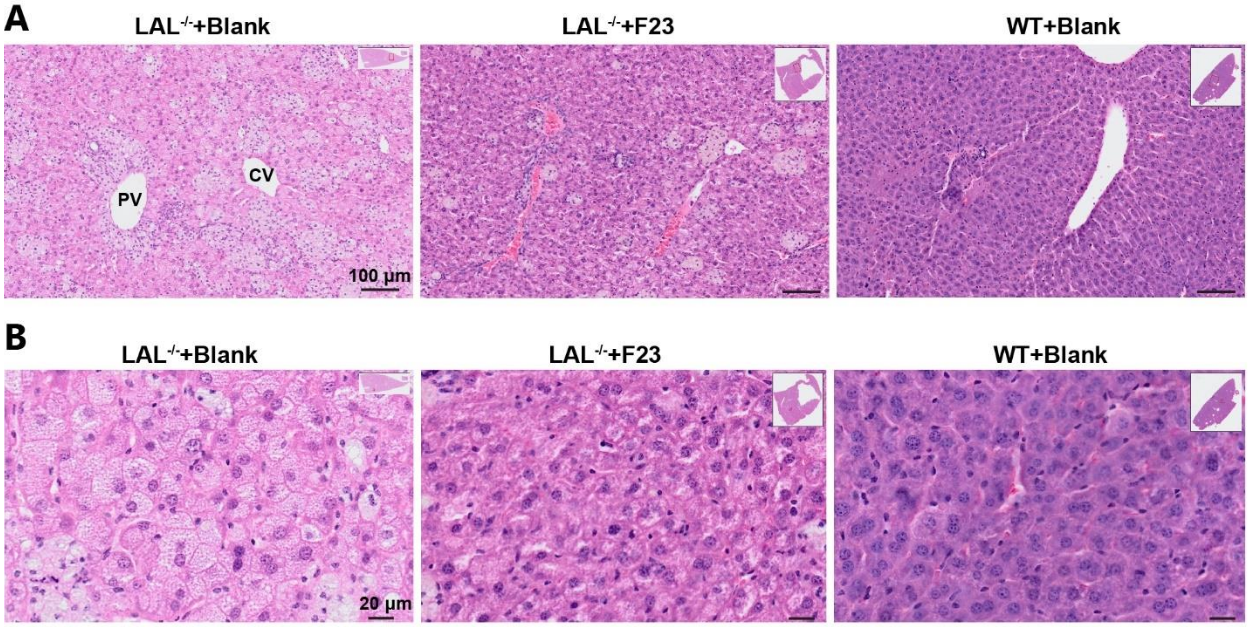


**Figure S21. H&E staining of liver tissue. (A)** Representative image with lower power view depicting the portal vein (PV) and centrilobular vein (CV) locations in the liver showing the presence of necrotic hepatocytes, microvesicular steatosis, and immune infiltrates, which are reduced in F23-treated $LAL^{-/-}$ mice and not present in WT mice. Scale bar, 100 μm. **(B)** Magnification to better visualize the microvesicular steatosis spots in hepatic tissues and ballooning in hepatocytes, which is reduced by F23 treatment. Scale bar, 20 μm.

| Form. | Ionizable lipid Mol. ratio / Mol. percentage [%] | Helper lipid Mol. ratio / Mol. percentage [%] | Cholesterol Mol. ratio / Mol. percentage [%] | DMG-PEG Mol. ratio / Mol. percentage [%] | Size [nm] (N=3) | PDI (N=3) | Zeta potential [mV] (N=3) | EE [%] (N=3) |
|---|---|---|---|---|---|---|---|---|
| F01 | DLin-MC3-DMA 50/50 | DSPC 10/10 | Cholesterol 38.5/38.5 | DMG-PEG 1.50/1.50 | 101.04 ± 11.84 | 0.179 ± 0.078 | -1.54 ± 2.77 | 91.08 ± 3.45 |
| F02 | DLin-MC3-DMA 50/48 | DSPC 15/14 | Cholesterol 38.5/36.67 | DMG-PEG 1.50/1.43 | 115.50 ± 15.84 | 0.242 ± 0.125 | -2.20 ± 5.22 | 89.38 ± 2.14 |
| F03 | DLin-MC3-DMA 50/45 | DSPC 20/18 | Cholesterol 38.5/35 | DMG-PEG 1.50/1.36 | 99.49 ± 22.02 | 0.236 ± 0.053 | -0.40 ± 6.92 | 89.46 ± 2.31 |
| F04 | DLin-MC3-DMA 40/44 | DSPC 15/16 | Cholesterol 35.0/38.25 | DMG-PEG 1.50/1.64 | 88.35 ± 7.54 | 0.238 ± 0.058 | -2.85 ± 2.24 | 85.03 ± 6.84 |
| F05 | DLin-MC3-DMA 40/38% | DSPC 25/23 | Cholesterol 40.0/37.56 | DMG-PEG 1.50/1.41 | 103.35 ± 16.01 | 0.251 ± 0.022 | -3.35 ± 2.40 | 74.41 ± 11.47 |
| F06 | DLin-MC3-DMA 40/44 | DSPC 10/11 | Cholesterol 38.5/42.78 | DMG-PEG 1.50/1.67 | 76.58 ± 7.07 | 0.175 ± 0.037 | -3.56 ± 7.58 | 92.30 ± 2.91 |
| F07 | DLin-MC3-DMA 25/35 | DSPC 15/21 | Cholesterol 30.0/41.90 | DMG-PEG 1.50/2.10 | 86.84 ± 19.30 | 0.241 ± 0.083 | -1.97 ± 3.16 | 94.31 ± 3.22 |
| F08 | DLin-MC3-DMA 30/32 | DSPC 15/16 | Cholesterol 45.0/47.87 | DMG-PEG 4.00/4.26 | 92.43 ± 28.80 | 0.294 ± 0.102 | +1.26 ± 4.57 | 89.59 ± 6.68 |
| F09 | DLin-MC3-DMA 50/50 | DOPE 10/10 | Cholesterol 38.5 / 38.5 | DMG-PEG 1.50/1.50 | 144.01 ± 20.76 | 0.152 ± 0.008 | -5.13 ± 2.45 | 58.23 ± 5.73 |
| F10 | DLin-MC3-DMA 40/46 | DOPE 15/17 | Cholesterol 30.0/34.68 | DMG-PEG 1.50/1.73 | 111.53 ± 7.42 | 0.158 ± 0.114 | -1.85 ± 2.53 | 79.34 ± 1.92 |
| F11 | DLin-MC3-DMA 35/43 | DOPE 15/18 | Cholesterol 30.0/36.81 | DMG-PEG 1.50/1.84 | 88.88 ± 9.57 | 0.176 ± 0.068 | -2.24 ± 1.73 | 87.18 ± 2.38 |
| F12 | DLin-MC3-DMA 30/35 | DOPE 20/23 | Cholesterol 35.0 / 40.36 | DMG-PEG 1.50/1.64 | 90.35 ± 25.57 | 0.204 ± 0.095 | -1.11 ± 5.32 | 94.41 ± 1.51 |
| F13 | DLin-MC3-DMA 50/50 | DOPC 10/10 | Cholesterol 38.5 / 38.5 | DMG-PEG 1.50/1.50 | 103.27 ± 5.28 | 0.187 ± 0.080 | -9.23 ± 3.87 | 78.02 ± 7.75 |
| F14 | DLin-MC3-DMA 30/39 | DOPC 15/20 | Cholesterol 30.0 / 39.22 | DMG-PEG 1.50/1.96 | 84.89 ± 11.35 | 0.226 ± 0.023 | -0.51 ± 2.30 | 78.70 ± 8.20 |
| F15 | DLin-MC3-DMA 40/42 | DOPC 20/21 | Cholesterol 35.0 / 36.46 | DMG-PEG 1.00/1.04 | 104.28 ± 6.84 | 0.215 ± 0.060 | -8.48 ± 8.55 | 72.13 ± 18.36 |
| F16 | DLin-MC3-DMA 40/39 | DOPC 15/15 | Cholesterol 45.0/ 44.43 | DMG-PEG 1.50/1.57 | 90.07 ± 9.17 | 0.269 ± 0.081 | -0.52 ± 3.05 | 89.71 ± 1.43 |
| F17 | DLin-MC3-DMA 50/50 | DMPC 10/10 | Cholesterol 38.5 / 38.5 | DMG-PEG 1.50/1.50 | 105.05 ± 5.38 | 0.197 ± 0.087 | -0.89 ± 5.54 | 71.29 ± 9.21 |
| F18 | DLin-MC3-DMA 40/42 | DMPC 15/16 | Cholesterol 38.5 / 40.53 | DMG-PEG 1.50/1.58 | 106.47 ± 10.46 | 0.120 ± 0.080 | -1.95 ± 7.60 | 77.10 ± 8.44 |
| F19 | DLin-MC3-DMA | DMPC 20/20 | Cholesterol 40.0/39.41 | DMG-PEG 1.50/1.48 | 97.18 ± 12.31 | 0.200 ± 0.045 | +0.54 ± 4.39 | 74.28 ± 8.51 |

| | | | | | | | | |
|---|---|---|---|---|---|---|---|---|
| | 40/39 | | | | | | | |
| F20 | DLin-MC3-DMA 30/37 | DMPC 20/25 | Cholesterol 30.0 / 36.81 | DMG-PEG 1.50/1.84 | 81.04 ± 5.09 | 0.170 ± 0.092 | -8.85 ± 6.05 | 80.80 ± 8.75 |
| F21 | DLin-MC3-DMA 50/50 | DPPC 10/10 | Cholesterol 38.5 / 38.5 | DMG-PEG 1.50/1.50 | 107.68 ± 11.13 | 0.173 ± 0.082 | -0.55 ± 2.59 | 86.41 ± 9.73 |
| F22 | DLin-MC3-DMA 45/42 | DPPC 20/19 | Cholesterol 40.0 / 37.56 | DMG-PEG 1.50/1.41 | 98.54 ± 10.44 | 0.189 ± 0.044 | +3.25 ± 3.68 | 73.87 ± 8.99 |
| F23 | DLin-MC3-DMA 40/44 | DPPC 15/16 | Cholesterol 35.0 / 38.25 | DMG-PEG 1.50/1.64 | 86.95 ± 2.78 | 0.157 ± 0.075 | +0.06 ± 7.33 | 91.52 ± 1.36 |
| F24 | DLin-MC3-DMA 30/41 | DPPC 10/14 | Cholesterol 30.0/41.10 | DMG-PEG 3.00/4.11 | 87.01 ± 5.08 | 0.173 ± 0.078 | -2.61 ± 5.31 | 83.49 ± 6.68 |
| F25 | 5A2-SC8 50/50 | DSPC 10/10 | Cholesterol 38.5/38.5 | DMG-PEG 1.50/1.50 | 100.20 ± 5.56 | 0.185 ± 0.076 | +2.71 ± 0.87 | 84.62 ± 6.23 |
| F26 | 5A2-SC8 30/39 | DSPC 15/20 | Cholesterol 30.0/39.22 | DMG-PEG 1.50/1.96 | 95.04 ± 11.75 | 0.173 ± 0.060 | +2.33 ± 3.49 | 80.02 ± 15.99 |
| F27 | 5A2-SC8 50/47 | DSPC 20/19 | Cholesterol 35.0/33.02 | DMG-PEG 1.00/0.94 | 104.25 ± 10.25 | 0.214 ± 0.095 | +0.24 ± 2.64 | 88.06 ± 3.06 |
| F28 | 5A2-SC8 50/45 | DSPC 15/13 | Cholesterol 45.0/40.36 | DMG-PEG 1.50/1.35 | 97.26 ± 10.92 | 0.197 ± 0.081 | -7.11 ± 8.33 | 80.99 ± 12.13 |
| F29 | 5A2-SC8 40/44 | DSPC 10/11 | Cholesterol 38.5/42.78 | DMG-PEG 1.50/1.67 | 92.32 ± 6.70 | 0.252 ± 0.096 | +1.10 ± 5.39 | 81.66 ± 11.41 |
| F30 | 5A2-SC8 40/44 | DOPE 10/10 | Cholesterol 38.5.38.5 | DMG-PEG 1.50/1.50 | 97.72 ± 0.93 | 0.214 ± 0.084 | -1.61 ± 5.00 | 90.49 ± 0.61 |
| F31 | 5A2-SC8 40/46 | DOPE 15/17 | Cholesterol 30.0/34.68 | DMG-PEG 1.5/1.73 | 89.54 ± 3.78 | 0.159 ± 0.055 | -7.20 ± 3.00 | 85.79 ± 1.44 |
| F32 | 5A2-SC8 35/43 | DOPE 15/18 | Cholesterol 30.0/36.81 | DMG-PEG 1.5/1.84 | 90.04 ± 1.40 | 0.166 ± 0.067 | -0.99 ± 7.36 | 84.62 ± 0.53 |
| F33 | 5A2-SC8 30/35 | DOPE 20/23 | Cholesterol 35.0/40.46 | DMG-PEG 1.50/1.73 | 98.42 ± 1.89 | 0.212 ± 0.076 | +0.68 ± 4.43 | 85.51 ± 1.27 |
| F34 | 5A2-SC8 50/50 | DOPC 10/10 | Cholesterol 38.5/38.5 | DMG-PEG 1.50/1.50 | 114.47 ± 2.73 | 0.273 ± 0.111 | -1.33 ± 3.68 | 68.87 ± 2.22 |
| F35 | 5A2-SC8 30/39 | DOPC 15/20 | Cholesterol 30.0/39.22 | DMG-PEG 1.50/1.96 | 89.74 ± 0.79 | 0.151 ± 0.062 | +3.03 ± 6.55 | 83.84 ± 1.15 |
| F36 | 5A2-SC8 40/44 | DOPC 20/22 | Cholesterol 30.0/32.97 | DMG-PEG 1.00/1.10 | 96.53 ± 4.42 | 0.210 ± 0.100 | +1.66 ± 6.56 | 69.76 ± 2.42 |
| F37 | 5A2-SC8 40/38 | DOPC 15/14 | Cholesterol 45.0/43.27 | DMG-PEG 4.00/3.85 | 89.84 ± 0.99 | 0.156 ± 0.055 | +1.07 ± 3.10 | 81.07 ± 3.08 |
| F38 | 5A2-SC8 50/50 | DMPC 10/10 | Cholesterol 38.5/38.5 | DMG-PEG 1.50/1.50 | 118.89 ± 0.74 | 0.264 ± 0.109 | -4.21 ± 0.61 | 80.08 ± 1.58 |
| F39 | 5A2-SC8 40/38 | DMPC 15/16 | Cholesterol 38.5/40.53 | DMG-PEG 1.50/1.58 | 101.26 ± 1.40 | 0.168 ± 0.078 | +3.05 ± 4.01 | 80.95 ± 1.21 |
| F40 | 5A2-SC8 40/39 | DMPC 20/20 | Cholesterol 40.0/39.41 | DMG-PEG 1.50/1.48 | 126.99 ± 4.79 | 0.280 ± 0.113 | +0.35 ± 1.55 | 69.56 ± 2.47 |
| F41 | 5A2-SC8 30/39 | DMPC 15/20 | Cholesterol 30.0/39.22 | DMG-PEG 1.50/1.96 | 96.72 ± 2.00 | 0.213 ± 0.097 | +0.50 ± 1.78 | 84.59 ± 1.80 |
| F42 | 5A2-SC8 50/50 | DPPC 10/10 | Cholesterol 338.50.0/38.5 | DMG-PEG 1.50/1.50 | 113.21 ± 2.52 | 0.262 ± 0.112 | +1.86 ± 4.63 | 78.73 ± 7.23 |
| F43 | 5A2-SC8 45/42 | DPPC 20/19 | Cholesterol 40.0/37.56 | DMG-PEG 1.50/1.41 | 128.76 ± 1.12 | 0.260 ± 0.109 | -0.52 ± 3.37 | 67.47 ± 5.88 |
| F44 | 5A2-SC8 40/44 | DPPC 15/16 | Cholesterol 35.0/38.25 | DMG-PEG 1.50/1.64 | 92.73 ± 4.25 | 0.163 ± 0.069 | -3.58 ± 4.67 | 73.14 ± 2.51 |
| F45 | 5A2-SC8 30/41 | DPPC 10/14 | Cholesterol 30.0/41.10 | DMG-PEG 3.00/4.11 | 97.83 ± 4.92 | 0.221 ± 0.097 | +3.80 ± 7.64 | 83.70 ± 3.25 |

| | | | | | | | | |
|---|---|---|---|---|---|---|---|---|
| F46 | DLin-MC3-DMA 50/50 | HSPC 10/10 | Cholesterol 38.5/38.5 | DMG-PEG 1.50/1.50 | 102.34 ± 3.46 | 0.183 ± 0.077 | +7.95 ± 1.34 | 91.65 ± 3.16 |
| F47 | DLin-MC3-DMA 45/47 | HSPC 15/16 | Cholesterol 35.0/36.27 | DMG-PEG 1.50/1.55 | 91.17 ± 6.51 | 0.193 ± 0.083 | +6.01 ± 5.30 | 89.03 ± 4.69 |
| F48 | DLin-MC3-DMA 45/41 | HSPC 20/18 | Cholesterol 45.0/40.54 | DMG-PEG 1.00/0.90 | 114.53 ± 2.38 | 0.172 ± 0.082 | -2.78 ± 7.91 | 89.63 ± 3.76 |
| F49 | DLin-MC3-DMA 35/36 | HSPC 15/16 | Cholesterol 45.0/46.63 | DMG-PEG 1.50/1.55 | 100.15 ± 7.74 | 0.193 ± 0.089 | +2.19 ± 1.22 | 82.87 ± 1.51 |
| F50 | 5A2-SC8 50/50 | HSPC 10/10 | Cholesterol 38.5/38.5 | DMG-PEG 1.5/1.5 | 115.75 ± 5.52 | 0.218 ± 0.128 | +0.16 ± 5.92 | 91.73 ± 6.14 |
| F51 | 5A2-SC8 40/41 | HSPC 15/16 | Cholesterol 40.0/41.45 | DMG-PEG 1.50/1.55 | 115.69 ± 3.69 | 0.224 ± 0.105 | +6.97± 3.03 | 73.25 ± 10.34 |
| F52 | 5A2-SC8 35/43 | HSPC 15/18 | Cholesterol 30.0/36.81 | DMG-PEG 1.50/1.84 | 98.16 ± 3.00 | 0.195 ± 0.073 | +4.45 ± 6.34 | 70.42 ± 12.96 |
| F53 | 5A2-SC8 30/35 | HSPC 20/23 | Cholesterol 35.0/40.46 | DMG-PEG 1.50/1.73 | 115.66 ± 6.16 | 0.217 ± 0.086 | +4.17 ± 8.99 | 68.96 ± 12.99 |

**Table S1: Detailed description of the 53 formulations of the combinatorial h*LIPA* mRNA-LNPs library.** The composition of each lipid is given in molar ratio and molar percentage. The values of Size, PDI, encapsulation efficiency, and zeta potential are the average and standard deviations from 3 independent repeated production of each formulation.

| $P_{org}$ | Ratio | $P_{aq}$ | Flow rate [mL/min] |
|---|---|---|---|
| 200 | 1 | 200 | 4.11.38±0.22 |
| 200 | 1.1 | 220 | 6.63±0.28 |
| 200 | 1.2 | 240 | 7.19±0.36 |
| 200 | 1.3 | 260 | 7.540±05 |
| 200 | 1.4 | 280 | 7.71±0.01 |
| 200 | 1.5 | 300 | 8.37±0.48 |
| 400 | 1 | 400 | 9.25±0.22 |
| 400 | 1.1 | 440 | 10.17±0.45 |
| 400 | 1.2 | 480 | 11.46±0.18 |
| 400 | 1.3 | 520 | 15.12±05 |
| 400 | 1.4 | 560 | 16.34±1.45 |
| 400 | 1.5 | 600 | 19.37±2.56 |

**Table S2: Microfluidic set up.** Gradual increase of the pressure and yielded flow rate used for the optimization of the microfluidic mixing. The highlighted cells represent the chosen setup for the microfluidic mixing. Data show mean ± SD, n=3.

| Formulations | F01 | F23 | F27 | F30 | F46 | F50 |
|---|---|---|---|---|---|---|
| **Total Area** | 20.19±0.441 | 41.43±0.9615 | 20.19±0.4441 | 10.87±0.5285 | 15.98±1.003 | 20.46±0.7635 |
| **95% cConfidence Interval** | 19.31 to 21.06 | 39.54 to 43.31 | 19.31 to 21.06 | 9.838 to 11.91 | 12.01 to 16.94 | 17.96 to 22.95 |

**Table S3: Area under the curve of mRNA-induced LAL activity of the formulations assessed in the *in vitro* kinetic analysis.**

| Formulations | PBS | F01 | F23 | F27 | F50 |
|---|---|---|---|---|---|
| **Total Area (Liver)** | 0.000 | 1.645±0.2228 | 3.917±0.2823 | 1.213±0.2591 | 1.757±0.3559 |
| **Total Area (Spleen)** | 0.000 | 4.195±1.236 | 3.902±0.7724 | 4.513±0.5608 | 4.169±1.066 |
| **Total Area** | 0.000 | 5.84±0.456 | 7.819±0.327 | 5.726±0.785 | 5.926±0.642 |

**Table S4: Area under the curve of mRNA-induced LAL activity in healthy mice of selected mRNA-LNP formulations.** The values represent the MEAN ± SD of 3 AUC profiles per formulation.

| Formulations | PBS | F01,0.5µg/g | F23, 0.5µg/g | F01,1.0µg/g | F23, 1.0µg/g |
|---|---|---|---|---|---|
| **Total Area (Liver)** | 0 | 5.25±2.309 | 7.75±1.225 | 10.92±1l826 | 19.08±1.155 |
| **Total Area (Spleen)** | 0 | 7.083±1.368 | 14.073±1.093 | 16.183±2.281 | 24.393±1.666 |
| **Total Area** | 0 | 12.333±2.546 | 21.823±1.906 | 27.103±2.451 | 43.473±3.782 |

**Table S5: Area under the curve of mRNA-induced LAL activity in LAL$^{-/-}$ mice.** The values represent the MEAN ± SD of 3 AUC profiles per formulation.

| Liver | | Cholesteryl esters | | | Triglycerides | | | Phospholipids + Free Fatty Acids | | |
|---|---|---|---|---|---|---|---|---|---|---|
| LIPIDMaps | Designation | WT+Blank conc. [nmol/mg], n=9 | LAL$^{-/-}$+Blank conc. [nmol/mg], n=9 | LAL$^{-/-}$+F23 conc [nmol/mg], n=10 | WT+Blank conc. [nmol/mg], n=9 | LAL$^{-/-}$+Blank conc. [nmol/mg], n=9 | LAL$^{-/-}$+F23 conc [nmol/mg], n=10 | WT+Blank conc. [nmol/mg], n=9 | LAL$^{-/-}$+Blank conc. [nmol/mg], n=9 | LAL$^{-/-}$+F23 conc[nmol/mg], n=10 |
| C14:0 | Myristic acid | n.d. | 0.55 ± 0.211 | 0.5 ± 0.11 | 0.29 ± 0.24 | 0.28 ± 0.0823 | 0.28 ± 0.091 | 0.05 ± 0.017 | 0.09 ± 0.034 | 0.09 ± 0.02 |
| C15:0 | Pentadecanoic acid | n.d. | 0.34 ± 0.13 | 0.28 ± 0.0696 | 0.90 ± 0.0308 | 0.12 ± 0.0195 | 0.1 ± 0.021 | 0.06 ± 0.008 | 0.06 ± 0.012 | 0.06 ± 0.011 |
| C16:0 | Palmitic acid | n.d. | 25.11 ± 4.14 | 15.67 ± 5.067 | 11.98 ± 5.06 | 11.92 ± 2.05 | 10.80 ± 2.036 | 14.06 ± 1.026 | 10.70 ± 1.214 | 11.92 ± 1.182 |
| C16:1n7 | Palmitoleic acid | 0.06 ± 0.042 | 4.14 ± 0.12 | 3.38 ± 0.648 | 0.78 ± 0.501 | 1.48 ± 0.397 | 1.10 ± 0.191 | 0.24 ± 0.057 | 0.21 ± 0.064 | 0.27 ± 0.076 |
| C17:0 | Margaric acid | 0.12 ± 0.01546 | 1.87 ± 0.6791 | 0.87 ± 0.3796 | 0.15 ± 0.067 | 0.45 ± 0.0766 | 0.38 ± 0.073 | 0.40 ± 0.051 | 0.28 ± 0.039 | 0.31 ± 0.029 |
| C17:1 | cis-10-Heptadecenoic acid | n.d. | 0.54 ± 0.1597 | 0.56 ± 0.095 | 0.09 ± 0.0390 | 0.19 ± 0.0395 | 0.16 ± 0.026 | 0.06 ± 0.006 | 0.05 ± 0.006 | 0.05 ± 0.004 |
| C18:0 | Stearic acid | 1.28 ± 1.61 | 6.778 ± 1.778 | 3.92 ± 0.5071 | 0.96 ± 0.3913 | 3.02 ± 0.332 | 2.29 ± 0.2244 | 17.57 ± 2.74 | 11.48 ± 1.674 | 12.50 ± 1.26 |
| C18:1n9 | Oleic acid | 1.96 ± 1.91 | 84.09 ± 15.18 | 64.03 ± 10.30 | 20.35 ± 10.74 | 41.27 ± 9.15 | 28.44 ± 5.46 | 7.08 ± 0.58 | 8.06 ± 0.874 | 8.81 ± 1.003 |
| C18:1n7 | Cis-vaccenic acid | 0.04 ± 0.0051 | 2.54 ± 0.6821 | 1.94 ± 0.3810 | 0.61 ± 0.2711 | 1.64 ± 0.47 | 1.12 ± 0.16 | 0.94 ± 0.099 | 0.98 ± 0.181 | 1.09 ± 0.161 |
| C19:0 | Nonadecanoic acid | n.d. | n.d. | n.d. | n.d. | n.d. | n.d. | n.d. | n.d. | n.d. |
| C18:2n6 | Linoleic acid | n.d. | 21.5 ± 3.64 | 13.64 ± 5.04 | 14.43 ± 6.29 | 16.73 ± 2.074 | 14.97 ± 2.774 | 10.46 ± 1.240 | 5.28 ± 0.549 | 6.52 ± 0.811 |
| C18:3n6 | gamma-Linoleic acid | n.d. | 0.25 ± 0.101 | 0.28 ± 0.1808 | 1.65 ± 1.2881 | 1.58 ± 0.3857 | 1.13 ± 0.125 | 0.51 ± 0.097 | 0.15 ± 0.024 | 0.22 ± 0.042 |
| C18:3n3 | alpha-Linolenic acid | n.d. | 1.96 ± 1.2323 | 0.66 ± 0.2763 | 1.82 ± 0.9974 | 5.57 ± 1.03 | 3.20 ± 0.641 | 0.24 ± 0.048 | 0.11 ± 0.023 | 0.13 ± 0.028 |
| C18:4n3 | Stearidonic acid | n.d. | n.d. | n.d. | 0.99 ± 1.09 | 0.35 ± 0.099 | 0.33 ± 0.086 | 0.04 ± 0.032 | n.d, | n.d. |
| C20:2n6 | Eicosadienoic acid | n.d. | n.d. | n.d. | n.d. | n.d. | n.d. | n.d. | n.d. | n.d. |
| C20:3n9 | Mead acid | n.d. | 0.03 ± 0.0121 | 0.04 ± 0.0198 | 0.01 ± 0.0064 | 0.17 ± 0.0599 | 0.12 ± 0.033 | 0.03 ± 0.011 | 0.05 ± 0.011 | 0.06 ± 0.02 |
| C20:3n6 | Dihomo-gamma-Linolenic cid | n.d. | 0.05 ± 0.0282 | 0.06 ± 0.0471 | 0.05 ± 0.0186 | 0.40 ± 0.088 | 0.3 ± 0.039 | 0.41 ± 0.080 | 0.27 ± 0.051 | 0.37 ± 0.068 |
| C20:3n3 | Eicasatrienoic acid (ETE) | n.d. | n.d. | n.d. | n.d. | n.d. | n.d. | 0.04 ± 0.003 | n.d. | n.d. |
| C20:4n6 | Arachidonic acid | n.d. | 0.27 ± 0.128 | 0.18 ± 0.2011 | 0.17 ± 0.0795 | 2.34 ± 0.4467 | 1.49 ± 0.274 | 0.99 ± 0.177 | 1.93 ± 0.147 | 2.21 ± 0.214 |
| C20:4n3 | Eicosatetraenoic acid (ETA) | n.d. | n.d. | n.d. | n.d. | n.d. | n.d. | n.d. | n.d. | n.d. |
| C20:5n3 | eicosapentaenoic acid | n.d. | n.d. | n.d. | 0.05 ± 0.0232 | 0.11 ± 0.0359 | 0.08 ± 0.024 | 0.05 ± 0.013 | 0.02 ± 0.005 | 0.03 ± 0.004 |
| C22:5n3 | DPA | n.d. | 0.01 ± 0.0011 | n.d. | 0.08 ± 0.0306 | 0.08 ± 0.0280 | 0.08 ± 0.030 | 0.08 ± 0.019 | 0.08 ± 0.015 | 0.1 ± 0.02 |
| C22:6n3 | Docosapentaenoic Acid | n.d. | 0.05 ± 0.0066 | 0.03 ± 0.0095 | 0.38 ± 0.1499 | 1.41 ± 0.4231 | 0.73 ± 0.206 | 2.82 ± 0.264 | 1.37 ± 0.347 | 2.02 ± 0.396 |
| C20:0 | Arachidic acid | n.d. | 0.08 ± 0.0272 | 0.08 ± 0.0162 | 0.03 ± 0.0156 | 0.07 ± 0.0119 | 0.06 ± 0.012 | 0.09 ± 0.041 | 0.05 ± 0.006 | 0.06 ± 0.016 |
| C22:0 | Behenic acid | n.d. | n.d. | n.d. | n.d. | n.d. | n.d. | 0.33 ± 0.0127 | 0.24 ± 0.0072 | 0.23 ± 0.080 |
| C22:1 | Erucic acid | 0.01 ± 0.0029 | 0.04 ± 0.0067 | 0.03 ± 0.0091 | 0.02 ± 0.0048 | 0.04 ± 0.0052 | 0.04 ± 0.009 | 0.02 ± 0.002 | 0.04 ± 0.006 | 0.02 ± 0.011 |
| C24:0 | Lignoceric acid | n.d. | 0.02 ± 0.0012 | n.d. | n.d. | 0.02 ± 0.0029 | 0.02 ± 0.002 | 0.23 ± 0.061 | 0.24 ± 0.049 | 0.20 ± 0.060 |
| C24:1 | Nervonic acid | n.d. | 0.04 ± 0.0221 | 0.05 ± 0.043 | n.d. | 0.03 ± 0.009 | 0.03 ± 0.009 | 0.30 ± 0.058 | 0.77 ± 0.159 | 0.63 ± 0.206 |
| C26:0 | Cerotic acid | n.d. | n.d. | n.d. | n.d. | n.d. | n.d. | n.d. | n.d. | n.d. |

**Table S6: Targeted lipidomic analysis of hepatic samples from the efficacy study.** Cholesteryl esters (CEs), triglycerides (TGs), and phospholipids + free fatty acids (PLs+FFAs) fractions were obtained by the separation of the liver lipid extracts of LAL$^{-/-}$+Blank (n=9), LAL$^{-/-}$+F23 (n=10), and WT+Blank (n=9) mice. The values depict the average ± SD for each species in the associated cohort. N.d.: not detected.

| Spleen | | Cholesteryl esters | | | Triglycerides | | | Phospholipids + Free Fatty Acids | | |
|---|---|---|---|---|---|---|---|---|---|---|
| LIPIDMaps | Designation | WT+Blank conc. [nmol/mg], n=9 | LAL-/-+Blank conc. [nmol/mg], n=9 | LAL-/-+F23 conc [nmol/mg], n=10 | WT+Blank conc. [nmol/mg], n=9 | LAL-/-+Blank conc. [nmol/mg], n=9 | LAL-/-+F23 conc [nmol/mg], n=10 | WT+Blank conc. [nmol/mg], n=9 | LAL-/-+Blank conc. [nmol/mg], n=9 | LAL-/-+F23 conc[nmol/mg], n=10 |
| C14:0 | Myristic acid | n.d. | 0.11 ± 0.03 | 0.05 ± 0.02 | n.d. | 0.08 ± 0.04 | 0.05 ± 0.01 | 0.06 ± 0.007 | 0.12 ± 0.044 | 0.1 ± 0.0213 |
| C15:0 | Pentadecanoic acid | n.d. | 0.03 ± 0.004 | n.d. | n.d. | 0.01 ± 0.001 | n.d. | 0.04 ± 0.008 | 0.05 ± 0.009 | 0.05 ± 0.00659 |
| C16:0 | Palmitic acid | n.d. | 1.96 ± 0.31 | 0.83 ± 0.33 | 0.74 ± 0.4 | 1.65 ± 0.3 | 0.95 ± 0.19 | 6.41 ± 0.904 | 8.03 ± 0.989 | 7.93 ± 0.575 |
| C16:1n7 | Palmitoleic acid | n.d. | 0.29 ± 0.12 | 0.12 ± 0.04 | 0.07 ± 0.042 | 0.18 ± 0.05 | 0.11 ± 0.04 | 0.07 ± 0.012 | 0.11 ± 0.022 | 0.11 ± 0.022 |
| C17:0 | Margaric acid | n.d. | 0.06 ± 0.03 | 0.03 ± 0.02 | n.d. | 0.07 ± 0.01 | 0.03 ± 0.01 | 0.11 ± 0.016 | 0.12 ± 0.111 | 0.12 ± 0.011 |
| C17:1 | cis-10-Heptadecenoic acid | n.d. | 0.06 ± 0.01 | 0.04 ± 0.002 | n.d. | 0.04 ± 0.01 | 0.03 ± 0.01 | n.d. | n.d. | n.d. |
| C18:0 | Stearic acid | n.d. | 0.19 ± 0.03 | n.d. | n.d. | 0.17 ± 0.04 | n.d. | 1.54 ± 0.185 | 1.87 ± 0.2286 | 1.91 ± 0.2460 |
| C18:1n9 | Oleic acid | n.d. | 7.01 ± 1.73 | 3.37 ± 1.02 | 1.40 ± 1.00 | 5.61 ± 0.92 | 3.33 ± 1.43 | 2.61 ± 0.360 | 5.03 ± 0.3152 | 3.98 ± 0.4840 |
| C18:1n7 | Cis-vaccenic acid | n.d. | 0.2 ± 0.05 | 0.11 ± 0.03 | 0.06 ± 0.02 | 0.16 ± 0.03 | 0.12 ± 0.05 | 0.35 ± 0.058 | 0.53 0± .0515 | 0.49 ± 0.04585 |
| C19:0 | Nonadecanoic acid | n.d. | n.d. | n.d. | n.d. | n.d. | n.d. | n.d. | n.d. | n.d. |
| C18:2n6 | Linoleic acid | n.d. | 1.73 ±0.88 | 0.75 ± 0.25 | 0.42 ± 0.253 | 2.23 ± 1.01 | 1.23 ± 0.73 | 1.78 ± 0.324 | 2.28 ± 0.2945 | 1.96 ± 0.3682 |
| C18:3n6 | gamma-Linoleic acid | n.d. | 0.01 ±0.002 | 0.01 ± 0.002 | 0.01 ± 0.002 | 0.02 ± 0.004 | 0.02 ± 0.004 | 0.01 ± 0.002 | 0.02 ± 0.0034 | 0.02 ± 0.0055 |
| C18:3n3 | alpha-Linolenic acid | n.d. | 0.03 ± 0.01 | 0.02 ± 0.004 | 0.02 ± 0.005 | 0.06 ± 0.01 | 0.04 ± 0.007 | 0.02 ± 0.004 | 0.02 ± 0.002 | 0.02 ± 0.0023 |
| C18:4n3 | Stearidonic acid | n.d. | n.d. | n.d. | n.d. | n.d. | n.d. | n.d. | n.d. | n.d. |
| C20:2n6 | Eicosadienoic acid | n.d. | n.d. | n.d. | n.d. | n.d. | n.d. | n.d. | n.d. | n.d. |
| C20:3n9 | Mead acid | n.d. | n.d. | n.d. | n.d. | 0.01 ± 0.003 | n.d. | 0.01 ± 0.002 | 0.01 ± 0.0033 | 0.01 ± 0.0021 |
| C20:3n6 | Dihomo-gamma-Linolenic cid | n.d. | 0.02 ±0.01 | 0.01 ± 0.002 | 0.02 ± 0.002 | 0.06 ± 0.02 | 0.03 ± 0.011 | 0.19 ± 0.041 | 0.25 ± 0.0255 | 0.27 ± 0.0380 |
| C20:3n3 | Eicasatrienoic acid (ETE) | n.d. | n.d. | n.d. | n.d. | n.d. | n.d. | n.d. | n.d. | n.d. |
| C20:4n6 | Arachidonic acid | n.d. | n.d. | n.d. | n.d. | 0.29 ± 0.08 | 0.12 ± 0.054 | 1.00 ± 0.253 | 1.42 ± 0.1526 | 1.26 ± 0.149 |
| C20:4n3 | Eicosatetraenoic acid (ETA) | n.d. | n.d. | n.d. | n.d. | n.d. | n.d. | n.d. | n.d. | n.d. |
| C20:5n3 | eicosapentaenoic acid | n.d. | n.d. | n.d. | 0.01 ± 0.003 | 0.01 ± 0.005 | 0.01 ± 0.001 | 0.01 ± 0.002 | 0.01 ± 0.002 | 0.01 ± 0.0015 |
| C22:5n3 | DPA | n.d. | n.d. | n.d. | 0.01 ± 0.002 | 0.01 ± 0.003 | 0.01 ± 0.005 | 0.04 ± 0.008 | 0.04 ± 0.0078 | 0.04 ± 0.0098 |
| C22:6n3 | Docosapentaenoic Acid | n.d. | n.d. | n.d. | 0.02 ± 0.006 | 0.07 ± 0.016 | 0.05 ± 0.021 | 0.17 ± 0.021 | 0.22 ± 0.0371 | 0.23 ± 0.0425 |
| C20:0 | Arachidic acid | n.d. | n.d. | n.d. | 0.02 ± 0.003 | 0.03 ± 0.011 | 0.02 ± 0.002 | 0.06 ± 0.002 | 0.07 ± 0.0197 | 0.06 ± 0.0106 |
| C22:0 | Behenic acid | n.d. | n.d. | n.d. | n.d. | n.d. | n.d. | 0.09 ± 0.016 | 0.13 ± 0.0298 | 0.11 ± 0.0147 |
| C22:1 | Erucic acid | n.d. | n.d. | n.d. | n.d. | n.d. | n.d. | n.d. | n.d. | n.d. |
| C24:0 | Lignoceric acid | n.d. | n.d. | n.d. | n.d. | n.d. | n.d. | 0.12 ± 0.022 | 0.24 ± 0.0667 | 0.19 ± 0.0340 |
| C24:1 | Nervonic acid | n.d. | n.d. | n.d. | n.d. | n.d. | n.d. | 0.18 ± 0.043 | 0.34 ± 0.0896 | 0.29 ± 0.0409 |
| C26:0 | Cerotic acid | n.d. | n.d. | n.d. | n.d. | n.d. | n.d. | n.d. | n.d. | n.d. |

**Table S7: Targeted lipidomic analysis of splenic samples from the efficacy study.** Cholesteryl esters (CEs), triglycerides (TGs), and phospholipids + free fatty acids (PLs+FFAs) fractions were obtained by the separation of the spleen lipid extracts of $LAL^{-/-}$+Blank (n=9), $LAL^{-/-}$+F23 (n=10), and WT+Blank (n=9) mice. The values depict the average ± SD for each species in the associated cohort. N.d.: not detected.
